# Mapping the Human Ghost Proteome: Classification and Experimental Detection Biases in the Identification of Alternative Microproteins

**DOI:** 10.64898/2026.07.31.741785

**Authors:** Ana Montero-Calle, Alberto Peláez-García, Antonio J. Martín-Galiano, Rodrigo Barderas

**Affiliations:** Chronic Disease Programme, UFIEC, Instituto de Salud Carlos III, Majadahonda, E-28220 Madrid, Spain; Proteomics Unit, Core Scientific and Technical Units, Instituto de Salud Carlos III, Majadahonda, E-28220 Madrid, Spain; Centro Nacional de Microbiología, Instituto de Salud Carlos III, Majadahonda, E-28220 Madrid, Spain; CIBER Frailty and Healthy Aging, Madrid, E-28029 Madrid, Spain

**Keywords:** Alternative proteins, Ghost proteome, microproteins, colorectal cancer, OpenProt, proteomics

## Abstract

The discovery of alternative proteins (AltProts), translated from non-canonical ORFs, has expanded the human proteome and revealed a hidden layer known as the “ghost proteome”. Despite increasing evidence, AltProts detection remains challenging due to their small size, physicochemical heterogeneity, and lack of annotation. Here, we developed an integrated bioinformatic and proteomic workflow to benchmark the detection of reference proteins (RefProts), isoforms, and alternative microproteins (MicroAltProts) in colorectal cancer cells using four extraction protocols—HCl, RIPA buffer, RIPA with chloroform, and RIPA followed by 30 kDa filtration—combined with high-resolution data-independent acquisition mass spectrometry. We identified and quantified using the Orbitrap Astral mass spectrometer a total of 66,438 peptides corresponding to 12,584 different protein groups across methods, with RIPA-based extraction approaches providing the most comprehensive coverage. To reduce redundancy in the OpenProt database and focus on MicroAltProts, we curated the dataset by removing known isoforms and long proteins, yielding a non-redundant set of 183,937 MicroAltProts. K-means clustering based on eight ProtParam-derived features grouped MicroAltProts into four physicochemical clusters. Among them, 43 MicroAltProts (<200 amino acids) were experimentally validated by mass spectrometry and classified into tiers following recent recommended international guidelines. Cluster assignment of detected MicroAltProts revealed that HCl extraction favored disordered, alkaline proteins, while RIPA-based protocols enabled the identification of membrane-associated and amphipathic α-helical MicroAltProts. Structural prediction indicated the presence of diverse folding determinants, including transmembrane helices, disordered regions, and nucleic acid-binding-like motifs. Altogether, this study provides a roadmap framework for the unbiased simultaneous detection of RefProts, isoforms, and AltProts, and supports a broader functional role for MicroAltProts.

## Background

In recent years, the classical view of the human proteome has been largely expanded with the discovery of alternative proteins (AltProts), translated from small open reading frames (sORFs) and alternative reading frames (altORFs) (1–4). These previously unattended proteins have cast doubt on the long-held assumption that only canonical ORFs encode biologically relevant polypeptides. Advances in next-generation sequencing, high-throughput ribosome profiling (Ribo-seq), and mass spectrometry (MS) have revealed a hidden layer of proteomic complexity, now referred to as the “ghost” proteome (2,3,5).

AltProts are often characterized by their short length together with high isoelectric point (pI), intrinsic disorder, or high hydrophobicity in comparison with reference proteins (RefProts) (6,7). This singular pattern has historically led to their exclusion from traditional gene annotation pipelines. However, dedicated databases such as OpenProt, SmProt, and sORFs.org now curate hundreds of thousands of AltProts predicted *in silico* or validated experimentally (8–11). For instance, OpenProt alone contains over 300,000 protein sequences, a majority of which exhibit minimal sequence similarity to canonical proteins (10,11).

Identifying AltProts remains a significant technical challenge (7,9). Ribo-seq has provided strong evidence of their translation, but only MS-based proteomics currently offers direct validation at the protein level itself (7,9). The development of high-resolution MS instruments, such as the Orbitrap Astral, in combination with data-independent acquisition (DIA), has significantly improved the detection sensitivity for low-abundance peptides (12), and thus can also include the identification of AltProts. Nonetheless, proteomic workflows must be carefully designed to avoid technical biases introduced during sample preparation and protein extraction (6,7,9,13). This consideration is particularly critical for MicroAltProts, defined as intrinsically short (<200 amino acids), often represented by only one or two detectable and flyable proteotypic peptides, and generally low abundant. In addition, MicroAltProts may exhibit even higher hydrophobicity, pI or intrinsic disorder —that differ substantially from those of most RefProts. As a consequence, their detection is especially sensitive to extraction chemistry, precipitation or partitioning steps, and downstream cleanup procedures, potentially leading to biased or misleading conclusions in AltProt research (6,7,9,14). Therefore, benchmarking extraction strategies is not merely a technical optimization but directly determines which regions of the alternative microproteome might become experimentally observable.

Despite growing evidence of their existence, the biological relevance of AltProts remains largely unexplored. Their roles in processes such as cell signaling, differentiation, malignancy, and stress response are only beginning to be understood (15–18). (15–18). Given the limiting applicability of homology-based approaches, classification of AltProts based on their predicted biochemical and structural properties would offer a valuable framework in practice to rationalize extraction biases and guide protocol selection. To address this knowledge gap, leveraging curated databases, classifying AltProts by their properties, and developing proteomic pipelines are needed.

In this context, we aimed here at identifying AltProts simultaneously with RefProt and isoform proteins in the KM12C colorectal cancer (CRC) cell line. Using four protein extraction methods combined with high-resolution mass spectrometry, we investigated how different extraction strategies influence the detection of proteins with distinct physicochemical properties represented in the OpenProt database. To support our analysis, the OpenProt database was computationally filtered and curated, identifying 183,937 MicroAltProts shorter than 200 amino acids and classifying them into four feature-based clusters as a surrogate for their potential cellular functions or subcellular localization. Based on our results, we propose a framework for the simultaneous identification of MicroAltProts, RefProts, and protein isoforms using conventional sample preparation methods, high-throughput, high-resolution MS DIA workflows, and the curated here OpenProt database, while taking into account the biophysical properties and flyability of MicroAltProts.

## Materials and methods

### Experimental Design and Statistical Rationale

The study was designed to i) compare how four extraction protocols (HCl, RIPA, RIPA + chloroform, and RIPA + 30 kDa filtering) affect recovery and detectability of RefProts, isoforms, and MicroAltProts (<200 aa) by DIA–MS in a CRC cell context, and ii) derive a property-based classification of MicroAltProts after database curation to reduce homology-driven redundancy. All extractions were performed from the same human CRC cell line to control for biological background.

For each extraction protocol, biological replicates (n = 3) were prepared independently from separate cell culture plates. Each biological replicate was injected once (no technical replication was used to avoid pseudo-replication). System suitability runs (HeLa digest) were conducted on batches of samples, which were used to monitor liquid chromatography coupled to tandem mass spectrometry (LC–MS/MS) stability (peak width, retention time (RT) drift, and precursor intensity coefficients of variation (CVs).

For counts (peptides/proteins by category), differences among methods were assessed by χ² tests or Fisher’s exact where appropriate. For continuous physicochemical properties (molecular weight (MW), pI, GRAVY) of method-specific proteins, distributions were compared using Kruskal–Wallis with Benjamini–Hochberg false discovery rate (FDR) (q = 0.05) across multiple features. Because the primary goal was method profiling rather than differential expression at the protein level, we emphasized effect sizes and overlap analyses (Venns) over classical fold-change testing.

To classify MicroAltProt_NR90, we computed eight global features per sequence (MW, pI, GRAVY, aromaticity, flexibility, instability index, α-helix content, and β-turn content). Features were z-scored prior to unsupervised learning. K-means clustering was run for k=2–20; the inertia (elbow) criterion, complemented by silhouette inspection, supported k=4 as the optimal natural grouping. PCA was used for visualization and to estimate variance explained (PC1+PC2 ≈ 56%), with loadings used to interpret latent axes (α-helix content, MW, flexibility). Cluster differences were summarized with interquartile ranges (boxplots) and mean ± standard deviation (SD) (**Table 1**). All summary statistics are reported as mean ± SD.

**Table 1.**
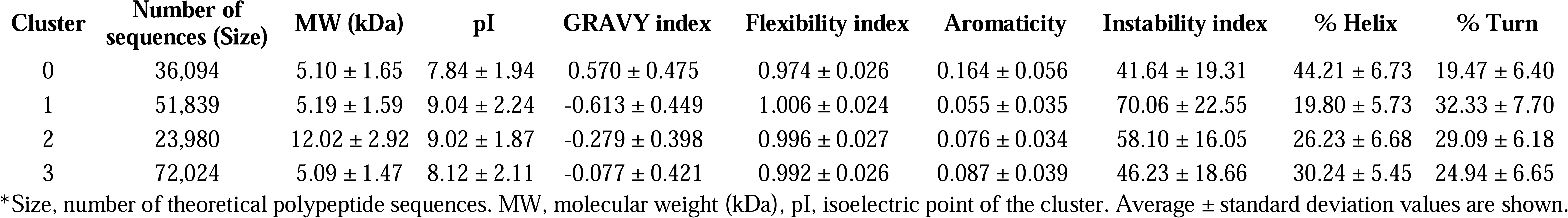
Physicochemical characteristics of the observed MicroAltProts clusters using the curated OpenProt database.

### Protein Extraction

The CRC cell line KM12C (RRID:CVCL_9547) from I. Fidler’s laboratory (MD Anderson Cancer Center) was used. CRC cells were cultured at 37°C and 5% CO_2_ in Dulbecco’s Modified Eagle Medium (DMEM, Lonza, Basel) supplemented with 10% fetal bovine serum (FBS, Sigma Aldrich), 1× L-glutamine (Lonza), and 1× penicillin/streptomycin (Lonza). Cells were routinely inspected for mycoplasma contamination.

For the experiments, KM12C cells were growth at 90% confluence in cultured treated T75 flask (Falcon). Then, cells were washed three times with 1x PBS (Corning) and incubated in culture medium without FBS for 1 h at 37°C and 5% CO_2_. After that, cells were washed three times with 1x PBS and incubated in culture medium without FBS for 48 h at 37°C and 5% CO_2_ to remove any trace of FBS. Next, cells were washed three times with 1x PBS and harvested with 1x PBS containing 4 mM EDTA (Carl Roth). After centrifugation at 1,200 rpm for 5 min, cell pellets were collected for protein extraction.

Four different strategies were used for protein extraction: i) RIPA, ii) RIPA with chloroform, iii) HCl, and iv) 30 kDa filtering, according to previous publications (19). Briefly, for RIPA extraction, cell pellets were resuspended in 500 µL of RIPA buffer (Sigma-Aldrich) and manual disaggregated using 16G and 18G needle syringes until homogeneity was observed. Then, samples were centrifuged at 12,000 g and 4°C for 30 min, and supernatants stored in a new tube. For RIPA with chloroform extraction, supernatants obtained after centrifugation following RIPA extraction protocol were incubated with 125 µL of chloroform (Sigma-Aldrich) and 500 µL of H_2_Omq, and centrifuged at 12,000 g at room temperature (RT) for 10 min. Supernatants were then dried under vacuum in a Speed-Vac (Eppendorf) and resuspended in 700 µL of 50 mM ammonium bicarbonate (ABC, Thermo Fisher Scientific) buffer pH 8.0. For HCl extraction, cell pellets were incubated with 200 µL of boiling water at 100°C for 10 min. Then, after sample centrifugation at 12,000 g and 4°C for 10 min, the aqueous phase was stored in a separate tube, and the cellular pellet was incubated with 400 µL of 50 mM HCl 0.5% DTT (Sigma-Aldrich) and manual disaggregated using 16G and 18G needle syringes until homogeneity was observed. After centrifugation at 12,000 g for 30 min at 4°C, supernatants were incubated with the aqueous phase previously stored and with 125 µL of chloroform and 500 µL of H_2_Omq. Finally, samples were centrifuged at 12,000 g and RT for 10 min, and supernatants were dried under vacuum in a Speed-Vac and resuspended in 700 µL of 50 mM ABC buffer pH 8.0. Finally, for the 30 kDa filtering strategy, 5 mg of protein extracts (supernatants) obtained after centrifugation following RIPA extraction protocol were diluted to a final volume of 4 mL with 50 mM ABC buffer pH 8.0, maintaining the same final RIPA buffer concentration for all three replicates, and filtered using 30 kDa AcroPrep 96-well filter plates (Cytiva) by centrifugation at 1,500 × g for 1 h at RT. The filtrate, containing low-MW proteins, was collected as the Filtered fraction. In parallel, the material retained on the top of the filter (high-MW proteins) was also recovered as the Non-Filtered fraction and used as a control.

All extraction buffers contained 1:100 diluted protease and phosphatase inhibitors (MedChemExpress). Protein concentration was measured by tryptophan quantification (20,21), and three biological replicates were analyzed for each extraction method.

The quality of the protein extracts was evaluated by Coomassie blue or silver staining following 10-17% PAGE-SDS gels (Fig. S1a).

### Proteomics Sample Preparation

After quantification, 15 µg of each protein extract were used for the MS analysis. Samples were incubated in a final volume of 500 µL of 1x PBS with 10 mM TCEP (Sigma-Aldrich) and 40 mM chloroacetamide (Sigma-Aldrich) for reduction and cysteine alkylation at 37°C, for 45 min, in rotation, and in the dark. Then, protein samples were incubated with hydrophilic and hydrophobic magnetic beads (Cytiva) as previously described (22–25), for 35 min, at RT, and in rotation. Once proteins were bound to the beads, beads were washed twice with 70% ethanol and once with 100% acetonitrile (ACN). For digestion, bead samples were incubated with trypsin (Thermo Fisher Scientific) at 37°C, overnight (O/N), and in rotation, diluted 1:20 (trypsin:protein) in 100 µL of 50 mM ABC buffer pH 8.0. After digestion, beads were sonicated for 2 min and supernatants stored in a new tube. Finally, beads were sonicated again for 2 min in 100 µL of 50 mM ABC buffer pH 8.0, and the second supernatant was stored with the previous one. Then, peptide samples were dried under vacuum in a Speed-Vac. Samples were resuspended in 15 µL 0.1% formic acid (FA) and 1 µL was used for LC-MS/MS analysis.

### DIA-LFQ LC-MS/MS

For LC-MS/MS, peptides were analyzed in an Orbitrap Astral mass spectrometer coupled to a Vanquish Neo UHPLC System (Thermo Fisher Scientific). Peptide samples were loaded into the precolumn PepMap Trap Catridge 5 µm, 300 µm x 5 mm (Thermo Fisher Scientific) and eluted in an Easy-Spray PepMap RSLC C18 3 µm, 75 µm x 15 cm (Thermo Fisher Scientific) heated at 50°C. The mobile phase flow rate was 300 nL/min and 0.1% FA in LC/MS Grade and 0.1% FA in 80% ACN were used as elution buffers A and B, respectively. The 15 min elution gradient was: 4%-10% buffer B for 2 min, 10%-40% buffer B for 11 min, 40%-99% buffer B for 0.5 min, and 99% buffer B for 1.5 min. Samples were analyzed in data independent acquisition (DIA) mode. For ionization, 1900 V of liquid junction voltage and 280°C capillary temperature was used. The full scan method employed a m/z 380-980 mass selection, an automatic gain control (AGC) value of 500%, and maximum injection time (IT) 5 ms. The MS/MS was performed with the Astral mass analyzer, using an AGC of 500%, an IT of 3 ms, and a normalized collision energy (NCE) of 25 for fragmentation of precursors. The scan range was set from 380 to 980 m/z, with an isolation window of 2 m/z, and window placement optimization was enabled. Thus, a total of 299 windows were analyzed in each cycle.

Raw data were analyzed with Spectronaut (version 21.0.260609.94842) using standardized workflows. DIA raw data and the human database from OpenProt, containing Reference proteins (RefProts), Isoforms, and Alternative proteins (AltProts) sequences, (November 2022, 309,678 protein entries, https://api.openprot.org/api/2.0/HS/downloads/human-openprot-2_0-refprots+altprots+isoforms-GRCh38.p13-uniprot2022_06_01.fasta.zip) were used for the construction of the spectral library by directDIA and for subsequent analysis. Additionally, NextProt (September 2023, 42,382 protein entries) and UniProt (February 2026, 20,416 entries) databases of canonical proteins were used for the construction of the canonical revised RefProts spectral library by directDIA and subsequent analysis (26,27). Trypsin/P was selected as the digestion enzyme and a maximum of 2 missed cleavages were allowed. Carbamidomethylation of cysteines was set as a fixed modification, and methionine oxidation and N-terminal acetylation were set as variable modifications. For DIA analysis, standard workflow was used. The maximum FDR for peptide and global protein (experiment) identifications was set at 0.01, and at 0.05 for run-wise protein identifications, automatic cross-run normalization was enabled, and all identified peptides were used for protein quantification. MicroAltProts identified were classified into evidence tiers (reported in Table 2), following recommended recent guidelines (4,14). Tier 1A MicroAltProts (n = 4) were supported by i) ≥2 unique, non-nested peptides of ≥9 amino acids meeting stringent thresholds, or ii) 1 unique peptide with 100% coverage. Tier 2A MicroAltProts (n = 26) were supported by single-peptide of length ≥9 amino acids that warrant orthogonal confirmation. Finally, MicroAltProts (n=14) not meeting Tier 1A or 2A criteria were classified as low-confidence evidence. Universal Spectrum Identifiers (USIs) are provided for all MicroAltProt precursors-supporting XICs to enable independent inspection (**Table S6**). Protein inference was performed using the IDPicker algorithm. Non imputation was performed during peptide and protein identification and quantification with Spectronaut.

**Table 2.**
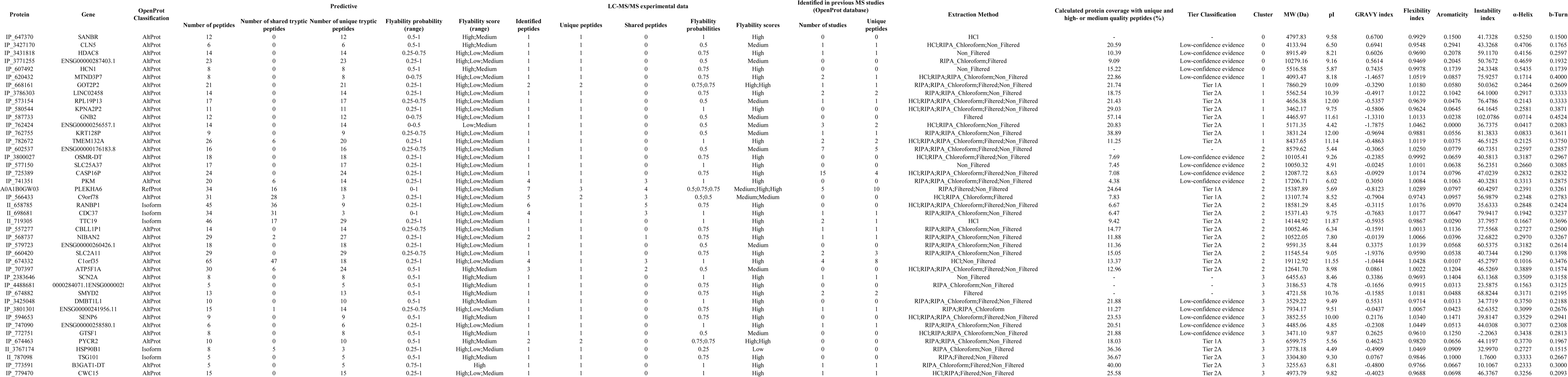
List of the 43 MicroAltProts identified in the CRC KM12C cell line with unique peptides across all extraction methods and their physicochemical properties.

The OpenProt database used in this study is composed of systematically translated annotated transcripts in three reading frames, and retains all ORFs longer than 30 codons that start with an AUG and end with a stop codon, independently of the transcript biotype or the presence of a canonical coding sequence. These predicted ORFs are then categorized based on their similarity to known protein sequences. Sequences identical to annotated proteins are classified as reference proteins (RefProts), while those with partial similarity to known proteins from the same gene are labeled as isoforms. In contrast, sequences showing no significant similarity to any known protein are annotated as alternative proteins (AltProts), potentially originated from previously overlooked or unconventional translation events.

To ensure consistent annotation, all sequences undergo intra-species homology searches. Based on this, isoforms and AltProts are distinguished and assigned unique accession numbers: RefProts retain their standard identifiers, isoforms are labeled with the prefix II_, and AltProts with IP_.

Furthermore, the OpenProt database does not contain the total number of reviewed reference proteins. Thus, to ensure the identification of unique tryptic peptides of AltProts, we integrated two major curated repositories of reviewed human canonical RefProts, NextProt and UniProt (42,736 different canonical RefProts). Theoretical tryptic peptides of the OpenProt and canonical RefProt entries was simulated *in silico* with R Studio (version 4.3.1), allowing up to two missed cleavages. To account for potential incomplete cleavage events at lysine–proline (KP) or arginine–proline (RP) sites, the digestion was performed considering both possible outcomes—cleaved and uncleaved peptides at these positions and both cutting after R or K, with and without exception rules for P.

To curate the OpenProt database, protein sequences were downloaded from OpenProt resource (https://www.openprot.org/downloads; human-openprot-2_0, July 2024). AltProt proteins similar to known proteins were identified by comparing AltProt with the reference human SwissProt proteome available in Uniprot resource (26), using the easy-search mode of MMseq2 (28). Only unique proteins at 50% identity over 50% alignment length over the AltProts from OpenProt database were selected. Next, proteins above 200 residues were discarded. In this sense, we defined MicroAltProts as alternative proteins with a length of ≤200 amino acids. This threshold was selected as an inclusive criterion that encompasses canonical microproteins as well as somewhat longer alternative proteins encoded by non-canonical open reading frames. Previous studies have shown that the vast majority of experimentally supported alternative proteins and microproteins cluster below ∼150 amino acids, while definitions in the literature and in reference databases variably extend up to 200 amino acids depending on biological context and annotation strategy (29,30). Using as cut-off ≤200 amino acids, therefore, should allow for a comprehensive exploration of the alternative proteome, while avoiding arbitrary exclusion of validated small alternative proteins reported in prior studies. Then, the resulting MicroAltProts were clustered with CD-HIT at indicated identity and alignment length thresholds (**File S1**) (31).

All proteomics data are available via the PRIDE repository (PXD066336) (32).

### Computational Methods

To develop a flyability predictor, only tryptic peptides from the OpenProt database of at least three amino acids in length were retained. Then, a set of physicochemical properties was calculated for each tryptic peptide using R Studio (version 4.3.1) with the packages “Peptides”, “dplyr”, “tidyr”, and “stringr”. Six physicochemical properties were calculated based on previous publications (33): peptide length, MW, pI, hydrophobicity index (GRAVY index, Kyte–Doolittle scale), aromaticity index, and instability index. From this information, a heuristic flyability predictor was developed to estimate the likelihood of a peptide being efficiently ionized and detected by LC–MS/MS. Finally, four peptide properties were empirically selected based on prior knowledge of peptide ionization efficiency and detection likelihood: peptide length (7-25 amino acids (aa)), pI (5–9), GRAVY (-1 to +1.5), and instability index (<40). Each parameter contributed +1 point to the total score ranging from 0 to 4. The flyability probability was then expressed as a normalized score between 0 and 1 (score / 4), where higher values indicate greater predicted detectability by MS. Peptides were classified according to their flyability score as low (score < 0.25), medium (score = 0.5), or high (score > 0.75). The resulting flyability scores were integrated into the peptide dataset for subsequent analyses, including the assessment of the flyability distribution among unique and shared peptides identified experimentally.

Protein properties were calculated using the Bio.SeqUtils.ProtParam module available in Biopython project (34). The resulting values were normalized using sklearn.preprocessing.RobusScaler method of the scikit-learn project (35).

Proteins were clustered based on feature value profiles by Kmeans using the sklearn.cluster.KMeans method successively applying an increasing number of predefined clusters (n_clusters parameter) from 2 to 20.

Principal component analysis (PCA) was carried out using sklearn.decomposition.PCA method.

Three-dimensional protein structural models were generated homology-free with AlphaFold 2 server (https://www.alphafold.ebi.ac.uk/) (36). Protein disorder, secondary structure, and signal peptide/transmembrane helices were predicted with IUPRED3 (https://iupred3.elte.hu/) (37), PSIPRED 4.0 (https://bioinf.cs.ucl.ac.uk/psipred/) (38,39), and Phobius (https://phobius.sbc.su.se/) (40) servers, respectively. Proteomics data visualization and analysis was performed with the R Studio (version 4.3.1) as previously done (22–24,41), using “tidyverse”, “psych”, “gridExtra”, “scales”, “ggplot2”, “VennDiagram”, “Biostrings”, “Peptides”, and “seqinr” packages. Sample clustering was determined by PCA using the “stats” R package. For the analysis, a protein was considered as identified by a given extraction method if it was quantified in at least two out of the three biological replicates processed with that protocol.

### Statistical Analysis

All statistical analyses were performed using R Studio (version 4.3.1) and Python 3.11. Data distribution and variance were assessed using Shapiro–Wilk and Levene’s tests, respectively. For comparisons between multiple groups, one-way ANOVA followed by Tukey’s post hoc test or Kruskal–Wallis test with Dunn’s correction was applied, depending on normality. For pairwise comparisons, unpaired two-tailed Student’s t-test or Wilcoxon rank-sum test was used as appropriate. PCA and clustering were conducted using the stats and scikit-learn packages. p-values ≤ 0.05 were considered statistically significant unless stated otherwise. Graphs and data visualizations were generated using ggplot2, seaborn, and matplotlib.

## Results

In this work, we have implemented a comprehensive strategy combining experimental set-up, followed by bioinformatic curation to gain insight into nature and simultaneous detectability of RefProts and isoforms, together with alternative microproteins of less than 200 amino acids (MicroAltProts). We evaluated how different extraction methods affected protein recovery and protein detection in the KM12C CRC cell line using a DIA workflow on an Orbitrap Astral mass spectrometer and a curated OpenProt database. Non-redundant MicroAltProts were clustered according to their physicochemical properties, and the structural and biochemical features of the experimentally identified MicroAltProts were characterized to infer their potential biological roles.

### Evaluation of extraction methods for the identification of RefProts, Isoforms, and AltProts

To experimentally assess how four protein extraction methods—HCl, RIPA, RIPA with chloroform, and RIPA followed by 30 kDa filtering (yielding Filtered and Non-Filtered fractions)—affect the recovery and, consequently, the detection of RefProts, isoforms, and AltProts from CRC cells, we set-up a DIA workflow using the high-resolution Orbitrap Astral mass spectrometer and the OpenProt database. These protocols were selected to compare distinct and widely used extraction chemistries that differ in detergent content and precipitation/partitioning behavior while remaining fully compatible with our downstream cleanup and LC-MS/MS workflow. Although chaotrope-based protocols (e.g., urea) are commonly used in proteomics, they were not included in this benchmark to avoid introducing additional variables related to chaotrope removal that could influence the recovery of low-molecular-weight proteins and peptides.

A total of 66,438 peptides corresponding to 12,584 different protein groups were identified and quantified. The interquartile ranges of peptide intensities per sample (Fig. 1a) and PCA (cumulative variance of PC1 and PC2: 54.4%) (Fig. 1b) supported the quality of the proteomics data (Table S1) and high reproducibility for all extraction methods, but for the first replicate of the 30 kDa filtering method and the third replicate of the RIPA with chloroform protocol.

**Fig. 1.**
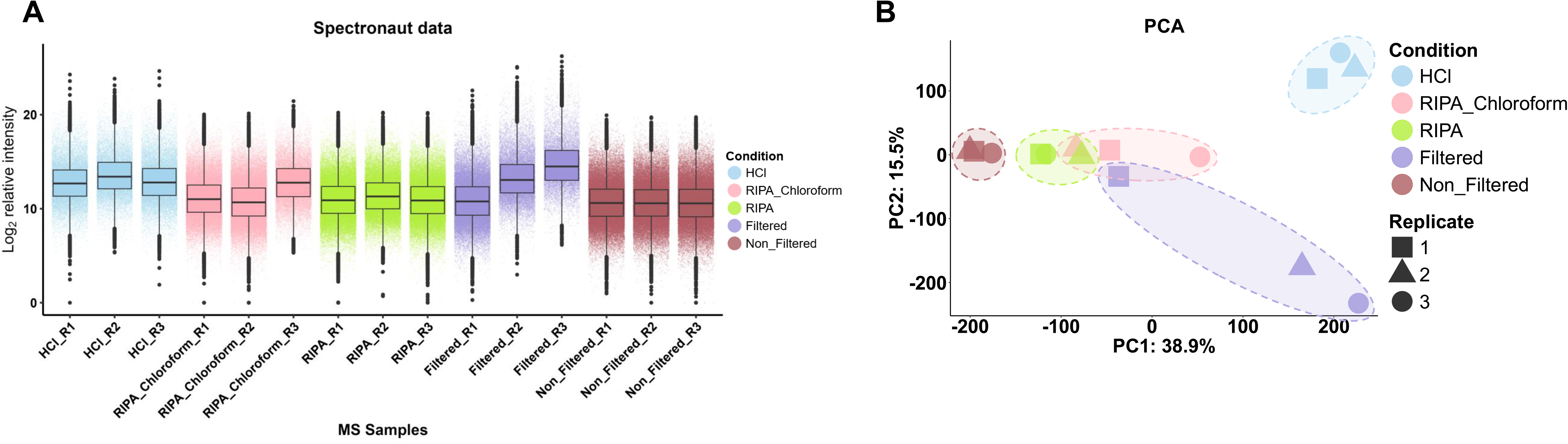
Comparative evaluation of extraction methods by mass spectrometry in KM12C colorectal cancer cells. **a** Boxplots showing the distribution of peptide intensities across replicates for each extraction method (HCl, RIPA, RIPA with chloroform, and RIPA followed by 30 kDa filtration). Median intensities are consistent across replicates, except for two outliers in the RIPA with chloroform and Filtered groups. **b** PCA of peptide-level intensities reveals clustering according to the extraction method used, indicating that each protocol recovers distinct subsets of the proteome.

Then, we focused on the analysis of the data derived from OpenProt to identify RefProts, Isoforms, and MicroAltProts. Despite the observed outliers in the PCA, a clear separation of samples according to the extraction method was observed, highlighting the distinct protein recovery profiles of each strategy. Notably, the two RIPA-based extraction methods clustered closely together, indicating highly similar proteomic profiles. Likewise, the fraction of high-MW proteins obtained after 30 kDa filtration (Non-Filtered) grouped in close proximity to both RIPA-based methods, consistent with its enrichment in high-molecular-weight proteins that are also efficiently recovered by RIPA extraction. In contrast, the HCl and 30 kDa-Filtered samples each formed distinct clusters. In addition, outliers were further analyzed by examining protein extracts with Coomassie blue or silver-stained, peptide and protein identification counts, total ion current (TIC), and pump pressure profiles during LC-MS/MS analysis (Fig. S1). For the 30 kDa filtering replicate, protein staining revealed the presence of high-molecular-weight protein bands that were largely absent from the other two filtered replicates, indicating incomplete removal of proteins above the filter cut-off. This observation was further supported by PCA, where this replicate clustered close to the RIPA-based samples rather than with the remaining filtered replicates. Consistent with this observation, this replicate yielded a markedly higher number of peptide and protein identifications, resembling the profiles obtained with the RIPA extraction methods rather than the remaining filtered samples (Fig. S1a-b). In contrast, the third replicate of the RIPA with chloroform extraction displayed a reduced number of peptide and protein identifications despite showing comparable protein profiles by protein staining (Fig. S1a-b). Importantly, TIC and pump pressure profiles were comparable across all injections (Fig. S1c-d), indicating that MS performance was not compromised. Therefore, these deviations were most likely originated from issues associated with sample preparation or extraction rather than LC-MS/MS acquisition. Importantly, downstream analyses were performed using cross-run normalization in Spectronaut, which compensates for global intensity differences across runs resulting in an overall higher signal intensity for RIPA with chloroform replicate 3 sample and Filtered samples replicate 2 and 3 despite fewer identifications.

Due to the high number of precursors and proteins identified, strict control of the FDR at both levels is required to limit false-positive identifications and ensure robust and reliable proteomic results. In this study, highly stringent FDR thresholds were applied: 1% at the precursor level and for experiment-wide (global) protein identifications, and 5% for run-wise (local) protein identification. Under these criteria, peptides were identified with FDR values ranging from 0 to 0.01, while proteins showed FDR values between 0 and 0.0099 at the global level and between 0 and 0.0499 at the run-wise level (Table S2 containing 3,379,043 precursor entries). Consistent with this stringent filtering, the estimated number of false-positive identifications was below 3,200 peptides and 620 proteins, with an experiment-wide expectation of 664 false peptides and 126 false proteins (Fig. S1b).

Using a filtering criterion of valid quantification in at least two biological replicates per extraction method, a total of 11,206 peptides and 3,370 protein groups were identified and quantified across the five sample groups (Fig. 2a). Peptides (n = 960) and proteins (n = 68) lacking valid values in all extraction methods were excluded from subsequent analysis. Notably, over 68% of peptide and 59% of protein identifications were exclusively detected using RIPA-based protocols (Fig. 2a and Fig. S2a). Nevertheless, fractionation of RIPA extracts using 30 kDa filter plates increased proteome depth primarily through the high-molecular-weight (Non-Filtered) fraction, which yielded the largest number of peptide and protein identifications. This fraction also showed the greatest overlap with both RIPA and RIPA with chloroform extracts, indicating that it retained most proteins recovered by these extraction methods while contributing additionally with a substantial number of unique identifications. In contrast, the low-molecular-weight (Filtered) fraction contained fewer total identifications but provided a complementary subset of peptides and proteins not detected in the other extraction strategies (Fig. 2a and Fig. S2a).

**Fig. 2.**
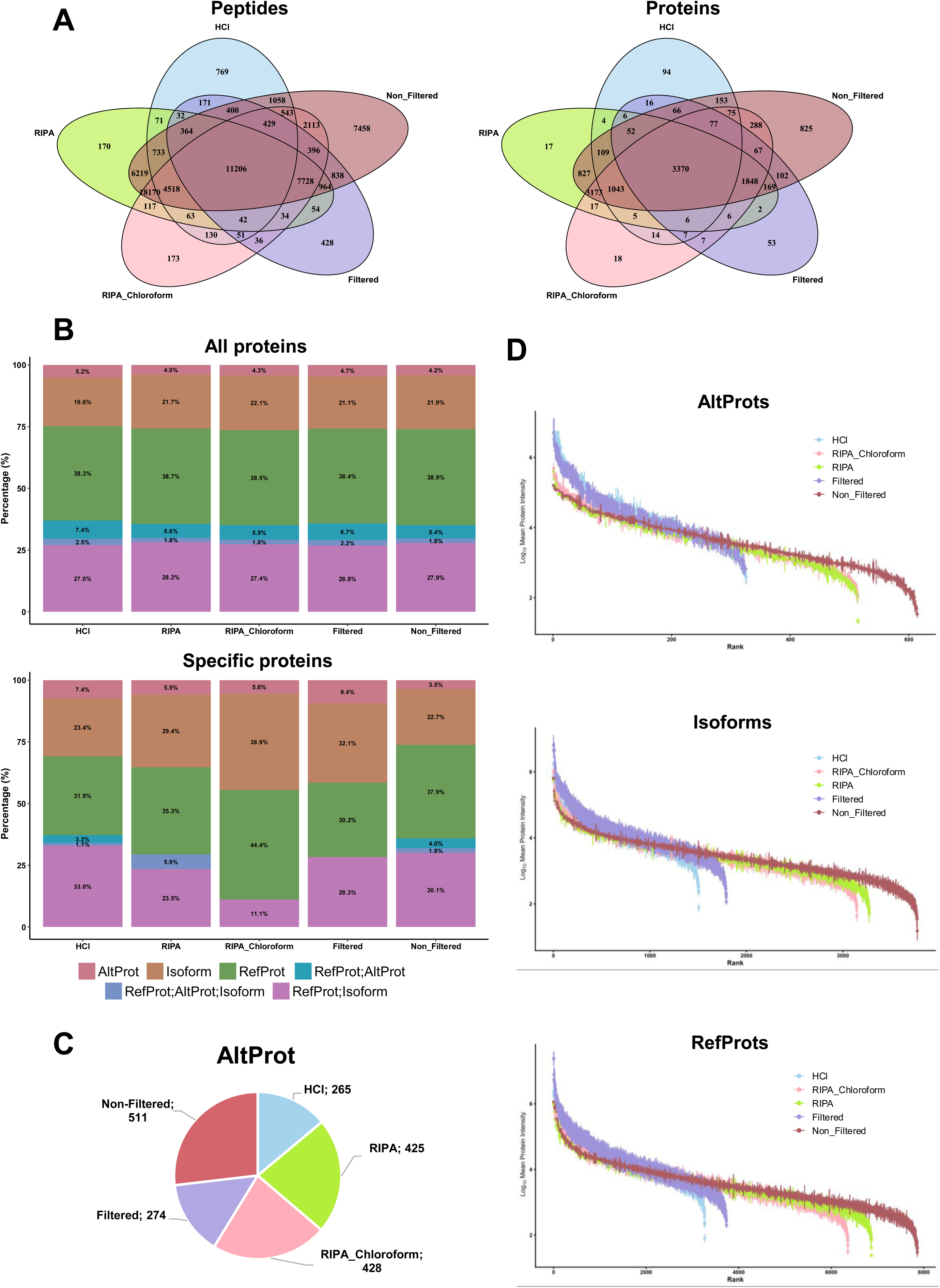
Impact of the tested extraction methods on the simultaneous identification of RefProts, Isoforms, and AltProts. **a** Total number of identified peptides and proteins per category (RefProts, isoforms, AltProts) using the OpenProt database across extraction methods. Non-Filtered samples yield the highest counts. **b** Proportion of protein identifications by category and method, highlighting method-specific biases. HCl and 30 kDa filtering favors AltProt- specific identifications, while RIPA methods enrich isoform identifications. **c** AltProt identifications among all extraction methods. The depicted pie chart shows the number of identified AltProts in each method-specific subset. **d** Ranked abundance distribution of AltProts, isoforms, and RefProts across extraction methods.

Importantly, 43 AltProts, 238 isoforms, and 373 RefProts were identified and quantified as specific of any of the different extraction methods without peptide overlap between protein categories in all three biological replicates (Fig. S2a), among the total of 53 AltProts, 2751 isoforms, and 4861 RefProts identified and quantified according to the OpenProt annotation (Fig. S2a). However, in all datasets, more than 26% of the identified proteins shared peptides with isoforms and RefProts, reflecting the high sequence similarity among them (Fig. 2b left). Furthermore, we observed a relatively high percentage of shared peptides between RefProts and AltProts with all extraction methods (over 5%) further illustrating the extensive sequence overlap between AltProts, isoforms, and RefProts (Fig. 2b left). Remarkably, the same proportion of RefProts was identified across all extraction methods (over 38%), whereas a higher percentage of isoforms was observed with the RIPA-based protocols compared to the HCl method (Fig. 2b left).

Importantly, identifications attributed to AltProts reached a non-negligible fraction ranging between 4% and 5.2% of the total proteins identified with the different extraction methods. Although the HCl samples showed a higher percentage of AltProts (Fig. 2b left), Non-Filtered samples showed a higher number of absolute AltProt identifications, followed by RIPA and RIPA with chloroform extraction protocols, owing to their substantially higher overall protein recovery (Fig. 2c). In addition, a similar number of AltProts were identified with the HCl and Filtered strategies. This trend was also observed for both isoforms and RefProts (Fig. S2b). Additionally, although AltProts showed lower overall numbers, a consistent set of 172 AltProts was detected with all extraction methodologies, whereas only 7 AltProts were not observed within the RIPA protocols (Fig. S2A). Similar trends were observed when only method-specific proteins were considered (Fig. 2b right). Interestingly, the RIPA-based protocols yielded a higher proportion of isoform-derived peptides in specific proteins, compared to the HCl protocol, whereas the HCl samples possessed a higher proportion of proteins with shared peptides between RefProts and isoforms.

Next, the influence of extraction methods on the abundance and the physicochemical properties of all identified and quantified proteins was evaluated. HCl and Filtered samples resulted in the identification of fewer proteins, mainly those that are highly abundant, thereby compromising the dynamic range and detection depth by MS (Fig. S2c). In contrast, RIPA, RIPA with chloroform, and Non-Filtered methods facilitated the identification of a substantially greater number of proteins, allowing for superior depth of proteome coverage (Fig. S2c).

Furthermore, we analyzed the abundance of the proteins specifically identified as AltProts, isoforms, or RefProts according to their annotation in the OpenProt database to investigate potential identification biases (Fig. 2d). As previously observed, RIPA, RIPA with chloroform, and, particularly, Non-Filtered fractions increased the identification of all three protein types, with a greater number of RefProts identified compared to isoforms or AltProts across all extraction methods. AltProts were found to be less abundant than RefProts and isoforms, whereas slight differences in abundance were observed among isoforms and RefProts (Fig. S2d). Additionally, the HCl-based and Filtered extraction methods yielded a narrower dynamic range dominated by highly abundant proteins, whereas all the other methods displayed a broader distribution encompassing a greater number of low-abundant proteins for all three protein types (Fig. 2d).

Collectively, these results demonstrate that RIPA-based extraction methods, particularly when combined with molecular-weight fractionation, provide broader proteome coverage, greater recovery of low-abundance proteins, and improved representation of RefProts, isoforms, and AltProts compared with the HCl protocol.

Regarding their physicochemical properties (Fig. S3a-b and Table S3), proteins specifically identified in the Filtered and RIPA with chloroform samples were generally smaller in size, with a mean MW of 16.65±14.26 kDa and 17.45±13.20 kDa, respectively. The Non-Filtered samples contained high MW proteins (49.37±48.90 kDa) and displayed the broadest MW distribution, whereas proteins specifically identified in the HCl samples spanned both low- and high-molecular-weight ranges, resulting in an intermediate mean MW of 44.59±43.36 kDa. RIPA- specific proteins displayed an intermediate MW distribution (20.59±9.75 kDa), shifted towards large proteins. Additionally, proteins specifically identified in the RIPA, RIPA with chloroform, and Filtered samples exhibited higher GRAVY values (-0.23±0.21, -0.40±0.49, and -0.46±0.33, respectively), indicating an increased recovery of relatively more hydrophobic proteins compared with the HCl (-0.69±0.32 GRAVY) and Non-Filtered (-0.49±0.43 GRAVY) methods. In contrast, proteins uniquely identified in the Filtered fraction displayed the highest isoelectric points (8.29±1.65), whereas RIPA and RIPA with chloroform samples were enriched in proteins with lower pI values (6.43±1.56 and 6.19±1.62, respectively). Furthermore, HCl and Non-Filtered specific proteins showed broader distributions for both GRAVY and pI, reflecting a more heterogeneous physicochemical composition.

Collectively, these findings demonstrate the existence of both numeric and qualitative sampling biases regarding the extraction methodology. RIPA-based extraction methods, particularly when combined with chloroform, favor the recovery of more hydrophobic proteins, whereas molecular-weight fractionation further enriched distinct subsets of proteins according to their physicochemical properties. Additionally, the selected method also influences the detection of specific proteoforms due to redundancy of the OpenProt database.

### Characterization of the OpenProt database used in the study

Given the observed extraction bias affecting the redundant detection of various proteoforms— especially within the low-MW range where most AltProts derived from alternative ORFs can be detected— it was essential to characterize the OpenProt database used in this study to ensure accurate interpretation of the proteomic landscape.

The OpenProt database version used in the study contains a total of 309,678 predicted protein sequences, enabling the detection and classification of non-canonical proteins that may play important roles in physiological and pathological contexts. According to its description, the dataset is broken down into 206,471 AltProts, 48,405 isoforms, and 54,802 RefProts (Fig. 3a). However, the database does not fully capture the annotated landscape of reference proteins, which may artificially inflate the apparent uniqueness of peptides assigned to alternative proteins. To overcome this limitation, we systematically integrated the two major curated repositories of reviewed human canonical RefProts, NextProt and UniProt (26,27). Together, these RefProt databases encompassed 42,736 different annotated proteins, of which only 3,763 were represented in OpenProt, leaving 38,973 actual RefProts from NextProt and UniProt absent in the OpenProt database, and being the remaining 51,039 entries unreviewed proteins (Fig. 3a). These observations highlight that peptide uniqueness is database-dependent and highly influenced by incomplete representation of the canonical proteome. Integration of all annotated AltProts, isoforms, and reviewed RefProts from OpenProt, UniProt, and NextProt generated a theoretical proteomic search space comprising 348,651 protein entries, including 206,471 AltProts, 48,405 isoforms, and 93,775 RefProts, encompassing both unique and common tryptic peptides across alternative and reference proteomes (Fig. 3a). In addition, two protein entries shared the same protein ID despite having different amino acid sequences, whereas 242 amino acid sequences were identical but associated with different protein IDs, indicating the presence of multiple isoform annotations.

**Fig. 3.**
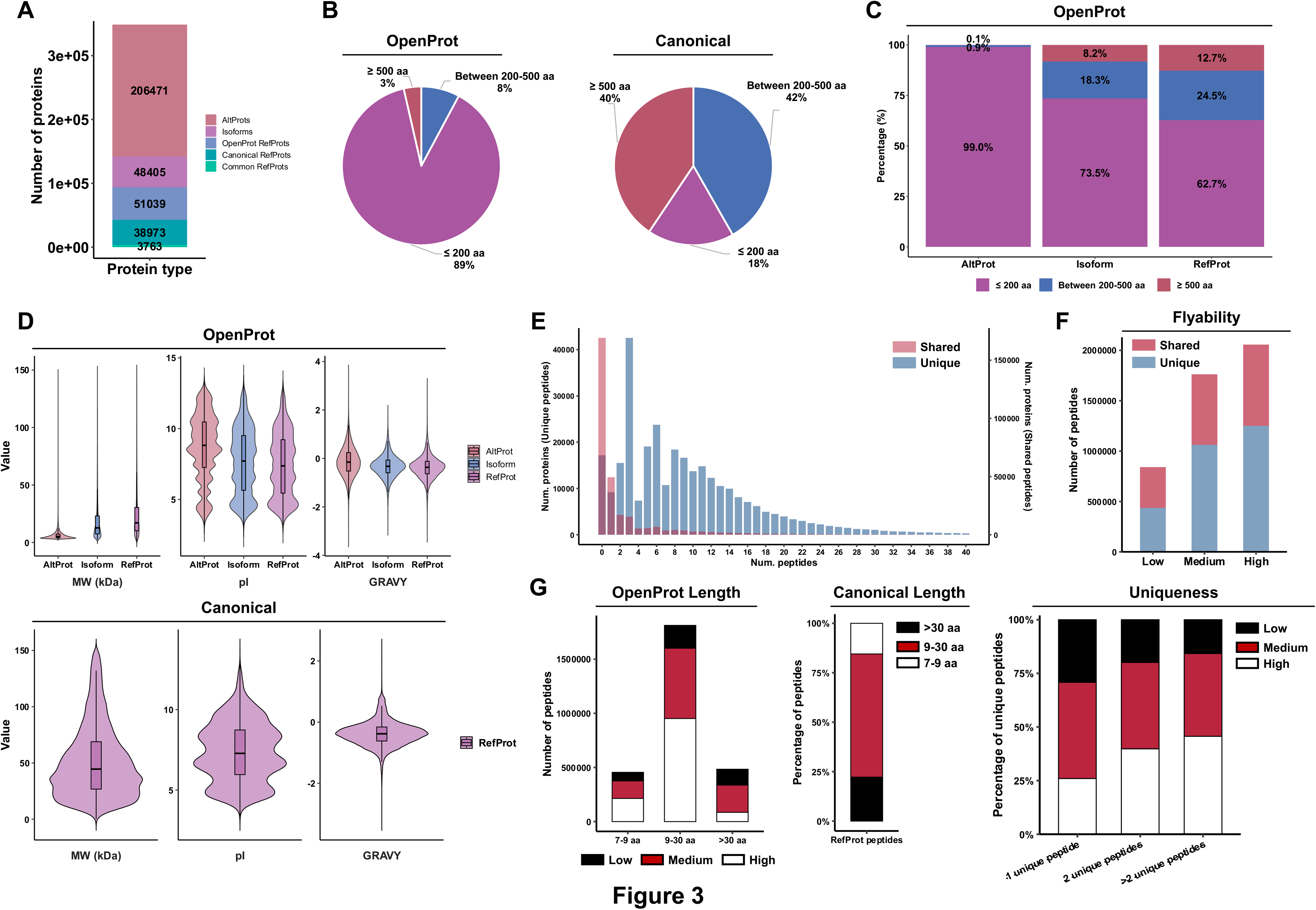
Physicochemical characterization of proteins from the OpenProt database. **a** Number of AltProts, Isoforms, and RefProts included in OpenProt and in the revised databases of canonical proteins, NextProt and UniProt. **b** Pie charts showing the distribution of RefProts, Isoforms, and AltProts from OpenProt, as well as canonical proteins from NextProt and UniProt, according to their amino acid length. **c** Stacked bar plot representing the relative proportion of proteins according to their length for AltProts, Isoforms, and RefProts included in OpenProt. **d** Violin plots comparing the MW, pI, and GRAVY score distributions between RefProts, Isoforms, AltProts, and canonical proteins from neXtProt and UniProt. **e** Distribution of OpenProt proteins according to the predicted number of tryptic unique (blue) or shared (red) peptides. Proteins containing unique peptides are shown in the left y-axis, whereas proteins containing shared peptides are shown in the right y-axis. **f** Distribution of OpenProt unique (blue) and shared (red) peptides across flyability score categories. **g** Distribution of unique peptides across low, medium, and high flyability score categories according to peptide length (left). Overall percentage of canonical protein peptides according to their length (Middle). Distribution of flyability score for peptides derived from proteins with one, two, or more than two unique peptides, illustrating the expected detectability of these proteins by mass spectrometry (right).

Most of the predicted sequences in the OpenProt database corresponded to short proteins, with 89% of the entries being ≤ 200 amino acids in length. Only 8% of entries fall within the intermediate size range of 200–500 amino acids, and 3% of entries with 500 amino acids or more (Fig. 3b). In contrast, canonical proteins displayed a markedly different size distribution, with 82% of proteins exceeding 200 amino acids and only 18% being ≤200 amino acids in length. Furthermore, clear differences in size were observed across the three OpenProt protein categories (RefProts, isoform, and AltProts) (Fig. 3c-d and Table S4). RefProts displayed the most heterogeneous distribution, with a substantial proportion of proteins longer than 500 amino acids (Fig. 3c). In contrast, isoforms and AltProts entries were predominantly composed of short sequences, with over 70% of isoforms and nearly all AltProts being shorter than 200 amino acids in length (Fig. 3c). Moreover, AltProts showed a strong bias towards small proteins below 10 kDa, in agreement with the enrichment in short proteins previously observed in MS experiments, whereas RefProts exhibited a significantly higher MW distribution consistent with their standard and often full-length structure, and isoforms showed intermediate MW values (Fig. 3d). Notably, when compared with the complete canonical human proteome, the RefProt subset represented in OpenProt was still enriched in smaller proteins, displaying a substantially lower MW distribution than canonical proteins as a whole (Fig. 3d). This observation indicates that the RefProts included in OpenProt database are not fully representative of the size distribution of the canonical proteome and are biased toward shorter protein species. This discrepancy demonstrates that the composition of the reference proteome embedded within OpenProt differs from that of comprehensive canonical protein databases, potentially influencing protein classification, peptide uniqueness, and the apparent contribution of alternative proteins in proteogenomic studies.

These findings reflect the fundamental distinctiveness of alternative proteins and suggest that many of them are likely to belong to the rough category of microproteins (17), which have been historically underrepresented and underexplored in proteomic studies.

Furthermore, we examined the distribution of pI and GRAVY physicochemical properties across all proteins in these databases, which are critical determinants of protein function, stability, and subcellular localization. Thus, their distribution may also provide insights into the distinct biological roles and characteristics of conventional and alternative proteins (Fig. 3d and Table S4). The distribution of pI values revealed a broad range, with most proteins ranging between 6 and 10, and a mean of 8.3. However, we could observe differences in the distribution of these properties across the three protein types (Fig. 3d). In terms of pI, AltProts showed a shift toward higher values (8.69±2.32) compared to isoforms, RefProts, and canonical proteins (7.67±2.30, 7.43±2.24, and 7.37±1.84, respectively), which reflect obvious differences in amino acid composition and, probably, in functional roles. Hydrophobicity profiles were similar between RefProts, isoforms, and canonical proteins, with AltProts displaying slightly higher mean GRAVY score, with a peak slightly below 0 (mean = -0.13±0.59), indicating a relative balance between the occurrence of hydrophilic and hydrophobic residues (Fig. 3d). Additionally, AltProts displayed a wider GRAVY distribution, mainly over the hydrophobic section, in comparison to RefProts, Isoforms, and canonical proteins, underscoring the profound physicochemical properties differences between AltProts and RefProts and isoforms.

Because most of the proteins from the OpenProt database are small proteins (≤ 200 amino acids in length), we simulated the number of potential tryptic peptides per protein and their flyability probability. Therefore, trypsin digestion of the 309,678 proteins from the OpenProt database and of the 38,973 canonical proteins not included in OpenProt was simulated (allowing up to two missed cleavages and both cutting after R or K, with and without exception rules for P), and a flyability predictor was subsequently developed based on previous publications (33). In total, 5,121,573 different tryptic peptides were generated for the OpenProt proteins, and 3,133,019 different tryptic peptides were generated for the canonical proteins from UniProt and NextProt. Furthermore, 63.69% of OpenProt peptides (3,261,825 peptides, corresponding to 304,749 proteins) were not found in external RefProt databases, and 36.31% (1,859,748 peptides) were shared with canonical proteins from NextProt and/or UniProt databases not included in OpenProt. Of the total of OpenProt specific peptides, a small fraction of proteins (5.63%, 17,168 proteins from 104,176 peptides) did not possess any unique tryptic peptide, and 8.11% of proteins (24,707 proteins) possessed only 1 or 2 unique tryptic peptides, which might hinder their identification by proteomics (Fig. 3e). These findings highlight the high degree of peptide redundancy between alternative and reference proteomes, thereby increasing peptide assignment ambiguity and reducing confidence in the identification of novel alternative proteins by LC-MS/MS.

Then, flyability of simulated tryptic peptides from the OpenProt database was calculated on the base of 7 or more amino acids in length (4,659,530 different peptides; 1,908,898 shared and 2,750,632 unique peptides). Among them, 18.05% exhibited low flyability scores (≤ 0.5), 37.81 % demonstrated medium scores (0.5), and 44.14% showed high flyability scores (≥ 0.75) (Fig. 3f). Focusing on unique peptides, we observed that the majority fell within the optimal range of 9-30 amino acids (65.88%), with more than 80% of them exhibiting high or medium flyability scores, supporting their efficient detection by MS (Fig. 3g). Notably, a substantial fraction of unique peptides (>455,000; 16.54%) was shorter than 9 amino acids. Despite their reduced length, approximately 80% of these peptides were also classified as having high or medium flyability, indicating that a considerable fraction of short peptides still possesses favorable physicochemical properties for MS identification (Fig. 3g). The overall peptide length distribution of OpenProt unique peptides was broadly similar to that observed for RefProt- derived unique peptides, with the 9–30 amino acid range representing the predominant peptide population in both databases. However, OpenProt exhibited a modest enrichment of short peptides (<9 amino acids) and a reduced proportion of long peptides (>30 amino acids) compared with canonical RefProts. Additionally, proteins identified with only 1 or 2 unique peptides (30,693 proteins) were predominantly supported by peptides exhibiting favorable flyability properties, with 78.03% of the corresponding peptides classified as having medium or high flyability scores, which might facilitate their identification by MS, whereas proteins identified with more than two unique peptides exhibited the highest proportion of medium- and high-flyability peptides (Fig. 3g). Thus, only 9.11% (28,200 proteins -5447 AltProts, 12,816 isoforms and 9,937 RefProts-) of OpenProt proteins could not be confidently identified by proteomics due to their short sequence (less than 7 amino acids), the absence of unique peptides, or the low flyability probability of their few unique peptides, which, together with the importance of protein abundance for their identification, make them particularly challenging or even inaccessible to current MS-based detection methods.

Finally, we applied this uniqueness and flyability scoring to the set of peptides identified in our proteomics analysis of KM12C cells using the OpenProt database. A total of 66,438 peptides were identified, of which 0.79% (528 peptides, corresponding to 433 different protein groups) were unique and 99.20% (65,910 peptides) shared among OpenProt and RefProt proteins. Among unique peptides, 61.85% and 29.79% of them showed high and medium flyability probability, respectively, whereas only 8.34% of them showed a low flyability probability, highlighting the strong performance of the flyability predictor developed in this study (Table S1).

Finally, to further assess the quality of the proteomics data we also searched against the canonical proteins databases. A total of 105,780 peptides and 15,071 different protein groups were identified and quantified using UniProt and NextProt databases, which confirmed the good quality of the data. A total of 76,130 peptides, corresponding to 12,463 protein groups, were shared among databases, and 29,650 peptides, corresponding to 5,460 protein groups, were only present in the canonical protein databases.

### Refining OpenProt for accurate identification of Alternative Microproteins

Given the biases and distinctive features observed in the characterization of the OpenProt database—particularly the strong enrichment in short AltProts and the redundancy affecting proteoforms—it became necessary to refine the dataset to improve confidence in protein identification and biological interpretation. While OpenProt offers a remarkable opportunity to explore the alternative proteome, its vast size, potential redundancy, especially among short polypeptides of uncertain origin, and absence of the complete RefProts canonical sequences pose major challenges for accurate analysis. Therefore, to address these issues, we curated the OpenProt database to focus specifically on truly novel and small alternative proteins, less than 200 amino acids —MicroAltProts— (Fig. 4), while excluding RefProts and isoforms.

**Fig. 4.**
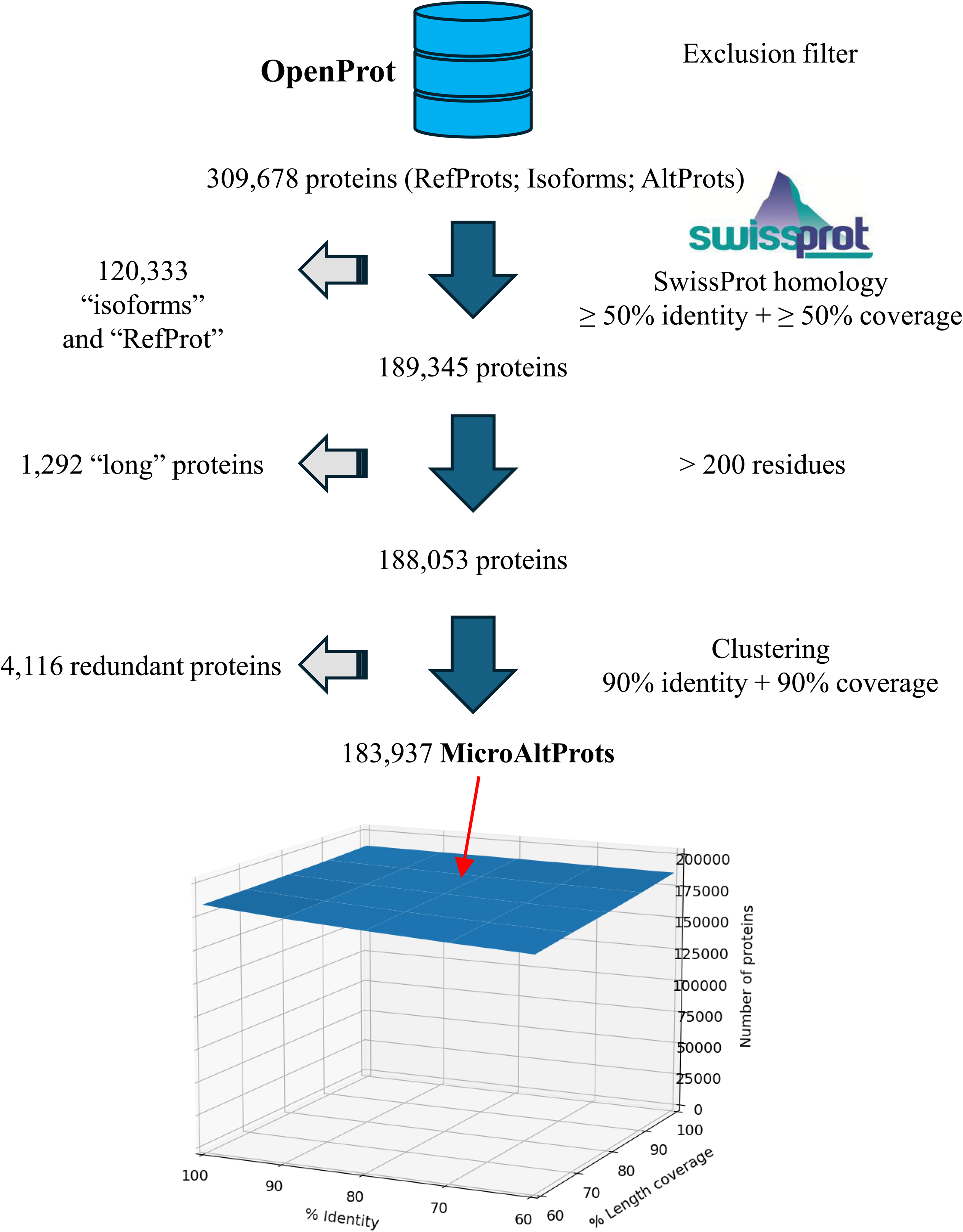
Data Processing Workflow for the OpenProt Protein Sequence Database Focused on Alternative Proteins. The initial OpenProt dataset includes 309,678 protein sequences. To reduce redundancy, sequences longer than 200 amino acids were clustered using 90% identity and 90% coverage, resulting in 189,345 isoform groups. Additionally, 1,292 long protein sequences were retained independently. After applying an exclusion filter that removed sequences with high redundancy (90% identity + 90% coverage), 4,116 redundant entries were discarded. Therefore, the final non-redundant curated dataset comprised 183,937 unique protein sequences. The workflow, and the number of surviving proteins after each step of the sequential application of exclusion filters (SwissProt closeness, length and sequence redundancy) is shown.

To this end, sequences of the original 309,678 proteins OpenProt database were compared with the SwissProt human proteome at 50% identity and 50% sequence coverage levels. Up to 38.86% proteins from the database shared relationships with reviewed human proteins above these thresholds and were deemed isoforms of those with mostly known functions. Next, a small fraction (0.42%) showed more than 200 residues and were also removed. This resulted in 188,053 MicroAltProt entries.

Finally, to assess the degree of redundancy within the dataset, we performed sequence clustering at an array of identity and alignment length thresholds (Fig. 4). Strikingly, grouping by sequence composition only produced minor size reduction, resulting in 183,937 MicroAltProt entries at 90% identity and 90% alignment percent (MicroAltProt_NR90) parameter values (Table S5, and File S1 for the fasta file as an open resource-). This MicroAltProt_NR90 fasta file database was mainly composed of proteins defined as AltProts in the OpenProt database (95.35%, 175,382 proteins), whereas only 3.33% (6,125 proteins) and 1.32% (2,430 proteins) of these MicroAltProts had been defined as isoforms or RefProts, respectively. Additionally, 15% of the proteins previously defined as AltProts by OpenProt were not included in the MicroAltProt_NR90 database after curation due to their length or the high sequence identity or homology with the SwissProt human reference proteome.

Collectively, our curation of the OpenProt database suggests that the theoretically alternative microprotein proteome is extensive—comprising about 180,000 distinct sequences—and highly diverse in sequence and origin.

### The AltProt Microprotein landscape organizes in four feature-based clusters

Next, given the limited sequence similarity among MicroAltProts, traditional protein classification approaches based on sequence alignment were not applicable. To circumvent this limitation, we applied an alternative strategy by means of unsupervised machine learning on MicroAltProts_NR90 based on calculation and normalization of eight physicochemical and structural properties: MW, pI, GRAVY index, aromaticity, flexibility, instability, α-helix composition, and β-turn composition.

Proteins were then clustered according to their normalized property value profiles with the K-means algorithm at a range from 2 to 20 preselected number of clusters. A cluster inertia-based analysis —the average distance of cluster members to their respective cluster centroid— indicated an optimal number of four clusters (Fig. 5a, left). These optimal natural clusters showed sizes ranging from 13 to 39% of the total of MicroAltProts_NR90, showing there is not a strongly dominant featural trend in the MicroAltProt proteome (Fig. 5a, right).

**Fig. 5.**
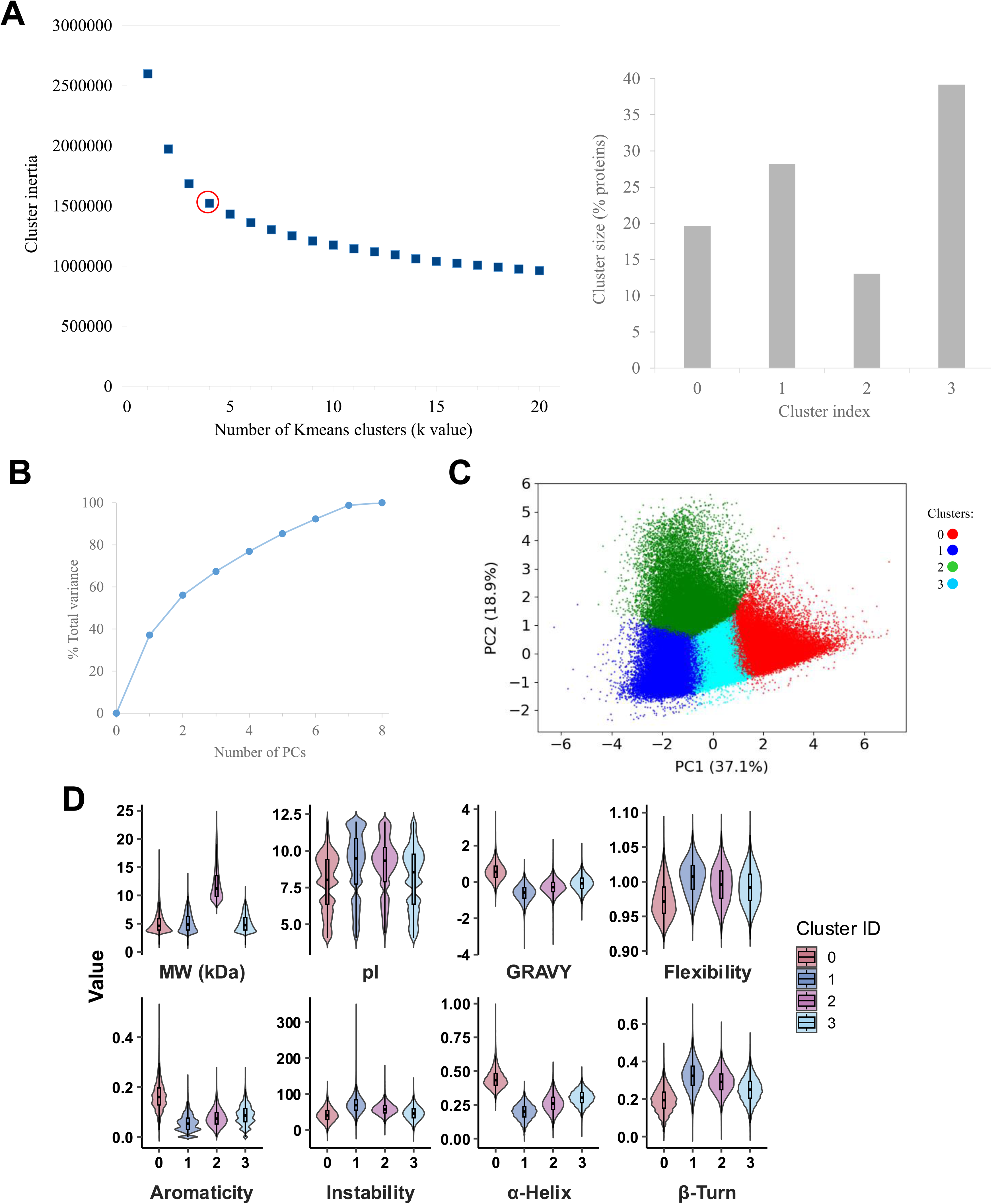
Sequence-based clustering and principal component analysis of alternative MicroProteins. **a** Elbow plot (left) showing the within-cluster sum of squares (inertia) for increasing k-values in K-means clustering. The optimal number of clusters was selected at k = 4 (red circle). Distribution of MicroAltProts across the four clusters (right), represented as the percentage of total entries. **b** Cumulative variance explained by the first 8 principal components (PCs) in the PCA, justifying dimensionality reduction. **c** PCA of protein sequence features reveals segregation according to the 4 clusters of MicroAltProts based on the first two principal components. Two-dimensional PCA plot (PC1 vs. PC2), colored by K-means cluster showed that clusters differed in charge, hydrophobicity, structural order, and molecular size. Cluster 0 (red) contains the highest hydrophobicity, aromaticity, and α-helix content MicroAltProts; Cluster 1 (blue) includes alkaline proteins with higher flexibility and instability indices, and increased β-turn content; Cluster 2 (green) featured the largest MW among MicroAltProts combined with high instability and flexibility; and Cluster 3 (light blue) includes a more balanced group with median values for MW, pI, hydrophobicity, instability, and secondary structure content. **d** Boxplots showing the distribution of molecular and structural properties of MicroAltProts across clusters: molecular weight (MW), isoelectric point (pI), hydrophobicity (GRAVY), aromaticity, flexibility index, instability index, and predicted content of α-helix, and β-turn.

Moreover, dimensionality reduction via PCA showed that PC1 and PC2 accounted for 56% of all data variance (Fig. 5b). This also allowed us to depict a 2D-projection of the AltProt predicted proteome, which cluster members mostly occupy discrete zones (Fig. 5c, Table S5).

Collectively, MicroAltProts within the different clusters generally exhibited varied physicochemical trends (*see* Fig. 5d for interquartile ranges and Table 1 for average and standard deviation values), such as size, hydrophobicity, instability, and α-helix content. Across clusters: Cluster 0 showed the highest hydrophobicity, aromaticity, and α-helix content, suggesting a hybrid population of cytoplasmic and transmembrane helical polypeptides. Cluster 1, in contrast, was characterized by an alkaline pI, markedly higher flexibility and instability indices, and increased β-turn content, supporting their classification as highly disordered MicroAltProts. Cluster 2 displayed the largest average MW combined with high instability and flexibility, pointing to longer but less stable proteins. Cluster 3 represented a more balanced group, with median values for most descriptors—MW, pI, hydrophobicity, instability, and secondary structure content—indicating that this set comprises MicroAltProts without extreme physicochemical biases. Collectively, this clustering scheme emphasizes three salient axes of diversity among MicroAltProts: high hydrophobicity and α-helix content (Cluster 0), basic pI —alkaline— (Clusters 1 and 2), high flexibility, instability, and turn content (Cluster 1) besides increased size (Cluster 2), and with Cluster 3 members showing average values for these features. This scheme for the three salient clusters mirrors the three properties deemed more important for the respective three top PCs of the PCA above: α-helix content, MW, and flexibility index, respectively, which would further likely influence their cellular location and effector functions.

The mostly unexplored nature of MicroAltProts makes them especially attractive as potential biomarkers and therapeutic targets, as they may capture disease-associated biological processes that are not reflected by the canonical proteome. Thus, to gain further insight into their biological significance, we next sought to comprehensively characterize these MicroAltProts. First, by comparing the amino acid frequencies between MicroAltProts and canonical RefProts, we observed a marked enrichment in cysteine (C) and serine (S), and a slight decrease in lysine (K) residues in MicroAltProts, which frequently serve as targets of post-translational modifications (PTMs) (Fig. 6a) (42). S constitutes a major target for phosphorylation-mediated signaling, C plays key roles in redox regulation and protein activity, and K is one of the most extensively modified amino acids, serving as a substrate for a wide range of PTMs, including ubiquitination, acetylation, methylation, SUMOylation, or succinylation, among others. These compositional differences suggest that MicroAltProts may be subjected to extensive post-translational regulation. Interestingly, acidic amino acids, aspartate (D) and glutamate (E), were clearly diminished in MicroAltProts. These two residues play important roles in protein folding, structural stability, and electrostatic interactions through their negatively charged side chains. Their reduced abundance may therefore contribute to the distinct physicochemical properties of MicroAltProts, potentially favoring increased structural flexibility and altered interaction dynamics. Furthermore, acidic residues are frequently involved in mediating protein–protein interactions and in establishing electrostatic contacts with nucleic acids. Consequently, the depletion of D and E may influence the ability of MicroAltProts to interact with DNA and RNA, suggesting that they could engage in molecular recognition and regulatory processes. Interestingly, tryptophan (W) was the most proportionally enriched amino acid in MicroAltProts, despite being the most underrepresented amino acid in the canonical proteome. Given the importance of W in mediating hydrophobic contacts, protein stability through π-π stacking, and molecular interactions, its overrepresentation may contribute to the distinctive structural and functional properties of this hidden layer of the proteome. Furthermore, R residues were also enriched in MicroAltProts. Given that trypsin specifically cleaves at the C-terminus of K and R residues, the increased abundance of R may contribute to the generation of shorter tryptic peptides from MicroAltProts, in concordance with the moderate enrichment of short peptides previously observed *in silico*. This compositional bias suggests that MicroAltProts possess unique physicochemical properties, potentially reflecting differences in structural organization, evolutionary constraints and regulatory potential.

**Fig. 6.**
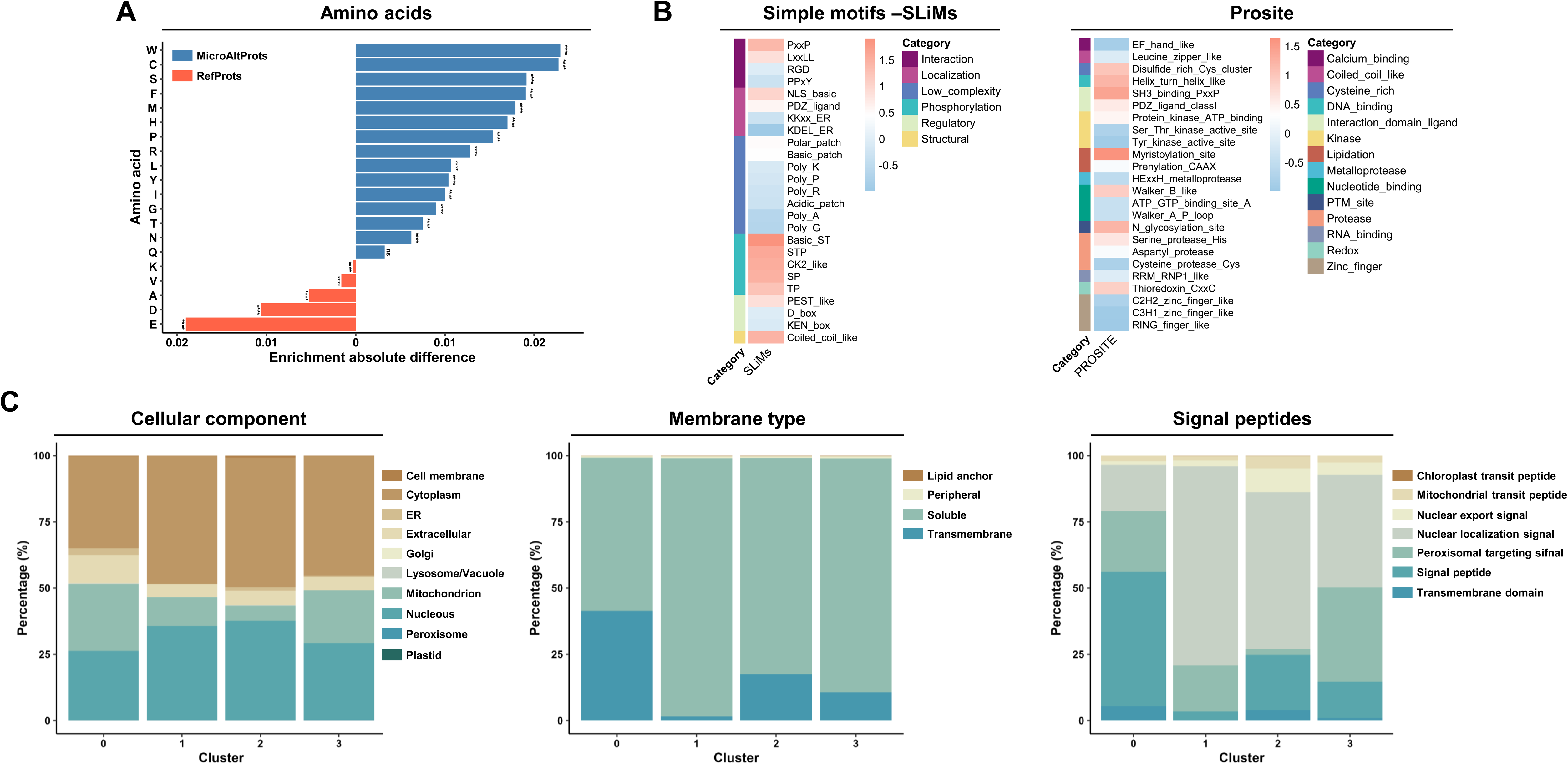
Compositional and functional characterization of MicroAltProts. **a** Absolute differences in amino acid frequency between MicroAltProts and canonical reference proteins (RefProts), highlighting amino acids enriched or depleted in MicroAltProts. **b** Predicted short linear motifs (SLiMs) and protein domains identified with PROSITE in MicroAltProts, grouped according to their functional categories. **c** Distribution of predicted subcellular localizations (left), membrane-associated localizations (middle), and localization signals across MicroAltProt clusters generated by DeepLoc.

To gain further insight into the potential functions of MicroAltProts, we systematically analyzed the prevalence of short linear motifs (SLiMs) and PROSITE patterns across MicroAltProts (Fig. 6b). SLiM analysis revealed a predominance of phosphorylation-related motifs, including Basic_ST, STP, SP, TP, and CK2-like motifs, suggesting that MicroAltProts might be extensively integrated into kinase-dependent signaling networks. Interaction-associated motifs, such as PxxP, LxxLL, and RGD, were also frequently detected, supporting a potential role in protein–protein interactions, particularly with SH3 domain containing proteins, nuclear receptors, and integrin-mediated signaling pathways. Additionally, motifs associated with subcellular targeting and molecular recognition were commonly observed, including basic nuclear localization signals (NLSs), indicative of potential nuclear trafficking, and PDZ-binding motifs, which may facilitate the assembly of membrane-associated signaling complexes. PROSITE analysis further supported a regulatory role for MicroAltProts. The most prevalent signatures included myristoylation sites, SH3-binding motifs, PDZ ligand motifs, and cysteine-rich/disulfide-associated patterns, indicating potential involvement in membrane association, protein interaction networks, and redox-dependent processes. Moreover, the presence of RNA-related motifs, including RRM-like signatures, suggests that a subset of MicroAltProts may participate in RNA-associated regulatory mechanisms. Together, SLiM and PROSITE analyses indicate that MicroAltProts are enriched in motifs associated with signaling, post-translational regulation, membrane targeting, and molecular interactions rather than catalytic or structural functions. These findings support the hypothesis that MicroAltProts primarily act as regulatory molecules, potentially modulating cellular processes through interaction- and signaling-based mechanisms despite their reduced size.

Finally, to predict the subcellular distribution of MicroAltProts we used DeepLoc 2.1 (Fig. 6c and Fig. S4). Cluster 0 was primarily associated with the secretory pathway and membrane-related compartments, including the plasma membrane, Golgi apparatus, endoplasmic reticulum, mitochondrion, and extracellular space. This cluster was highly enriched in soluble and transmembrane proteins, showing the highest probability of containing transmembrane domains among all clusters. Furthermore, it was enriched in proteins harboring signal peptides, suggesting potential roles in intercellular communication, trafficking, and membrane-associated signaling. Cluster 1 displayed a predominantly intracellular distribution, with enrichment in cytoplasmic and nuclear localizations, and was enriched in soluble proteins. Additionally, it showed the highest probability of containing nuclear localization signal peptides, consistent with potential regulatory functions in intracellular signaling, gene expression, or nucleic acid-associated processes. Cluster 2 was enriched in cell membrane, Golgi apparatus, cytoplasmic, and nuclear components. Similar to cluster 0, these proteins are enriched in soluble and transmembrane proteins, but with the highest probability of containing lipid anchor domains among all clusters. Furthermore, nuclear export signals were highly enriched in cluster 2 proteins. Finally, Cluster 3 was enriched in cytoplasmic proteins, mitochondrion, and peroxisome components. In addition, this cluster was predominantly composed of peripheral and soluble proteins, showing the highest probability of containing peroxisomal targeting signal peptides.

### Physicochemical classification of MicroAltProts detected by LC-MS/MS

Next, after establishing a curated MicroAltProt database and defining the sets of shared and unique peptides among AltProts, isoforms, RefProts, and canonical proteins, we focused our analysis on experimentally detected MicroAltProts supported by unique peptides. Additionally, as we were able to distinguish clear clusters associated with specific physicochemical characteristics of MicroAltProts, we classified the experimentally detected MicroAltProts across all extraction methods based on their physicochemical properties. Among the 528 unique peptides identified, only 48 peptides corresponded to 43 different MicroAltProts were detected in at least 70% of the replicates for one or more extraction methods, showing all of them but one medium and high flyability scores (Table 2).

First, we evaluated whether the different extraction methods preferentially enriched specific MicroAltProt clusters and assessed potential biases in MicroAltProt identification across the different clusters (Fig. 7). Proteomic profiling revealed a differential pattern of detection of AltProts using the different extraction protocols. The RIPA-based protocols enabled the identification of 41 MicroAltProts in total, with only 2 MicroAltProts specifically identified with the HCl protocol (Fig. 7a). Additionally, a core of 7 MicroAltProts (16.3%) were identified with all extraction methods, and 4 MicroAltProts (9.30%) with all RIPA-based samples. However, the higher number of MicroAltProts were identified in the Non-Filtered fractions, followed by RIPA with chloroform samples (Fig. 7a).

**Fig. 7.**
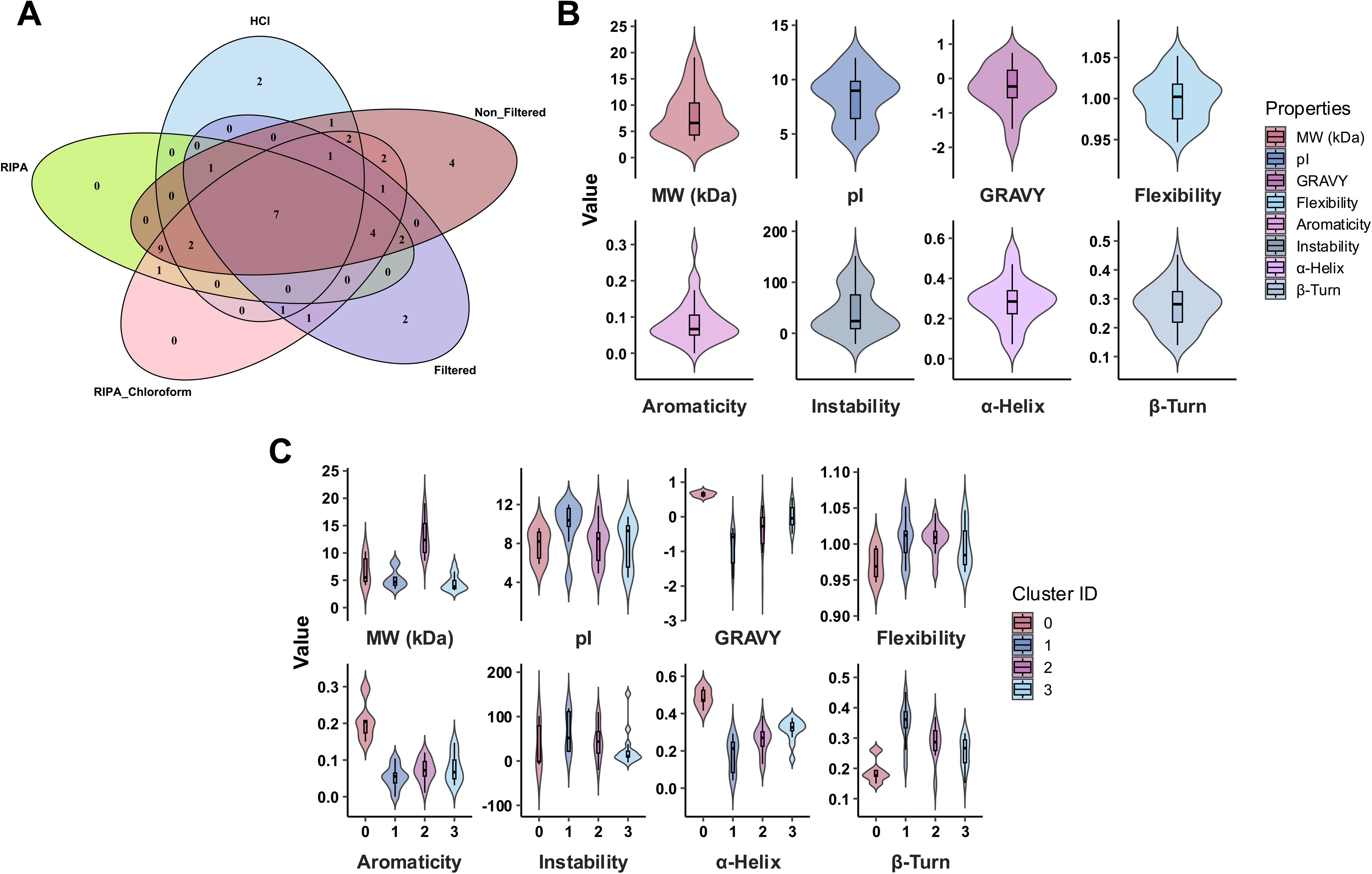
Comparative overlapping of protein extraction methods and physicochemical characteristics of globally detected AltProts. **a** Venn diagram showing the overlap of alternative microproteins identified using four extraction protocols: HCl, RIPA, RIPA–Chloroform, and RIPA followed by 30 kDa filtration protocol. **b** Violin plots displaying the distribution of selected physicochemical properties across all AltProts identified: molecular weight (MW), isoelectric point (pI), hydrophobicity score (GRAVY), flexibility, aromaticity, instability index, and secondary structure propensity (helix, turn, sheet). **c** Violin plots of the same 8 properties of the 44 experimentally validated MicroAltProts grouped according to the defined novel clusters in which they were identified (0–3).

Using the previously developed flyability predictor, we then estimated the flyability score for each of the unique peptides identified from these MicroAltProts (Table 2). The flyability score for most of them was high (34 peptides) or medium (15 peptides), with only 1 peptide showing a low flyability score, highlighting the high confidence of the identifications. Furthermore, 11 out of the 43 MicroAltProts had been identified in previous LC-MS/MS analysis included in the OpenProt database, with 9 of them identified with two or more unique peptides. Additionally, 43.18% of these MicroAltProts had not been previously identified by MS (19 MicroAltProts) and 29.54% had only been reported in a single study and with one unique peptide (13 MicroAltProts), according to the data retrieved from the OpenProt website (Table 2).

Because MicroAltProts are small and low-abundance proteins, as previously observed (Fig. S2D), with often particular physicochemical properties, their identification by MS is especially challenging, making stringent control of false-positive identifications essential to confidently confirm their detection. Proteome-wide FDR control is not enough to ensure high confidence in MicroAltProt identifications, as a global FDR threshold does not constrain the FDR within small and underrepresented protein classes, and further peptide-based investigation is mandatory according to the recently published guidelines for the proteomic detection of human microproteins (14). Therefore, to ensure confidence on the MicroAltProt identifications, extracted Ion Chromatograms (XICs) of the 56 precursors associated with the 48 peptides and 43 identified MicroAltProts were manually inspected by two independent evaluators to assess peptide quality. Only precursors supported by high-quality and well-defined XICs were considered reliable. All MicroAltProt-supporting XICs are provided as USIs together with representative annotated evidence (Table S6 and Fig. S5). Overall, 53.57% (30 precursors sequences) and 32.14% (18 precursors sequences) of precursors displayed a high- and medium-quality XICs, respectively. High-quality precursor XICs corresponded to 27 different peptides of 27 MicroAltPrtos, whereas medium-quality XICs corresponded to 17 different peptides of 15 MicroAltPrtos. In contrast, only a 14.3% of precursors (8 precursors sequences corresponding to 8 different peptides and proteins) displayed poor chromatographic quality. Consequently, protein sequence coverage was recalculated based exclusively on peptides supported by high-quality XICs and avoiding nested peptides (Table S6 and Table 2). Among the identified MicroAltProts, 9 of them showed protein sequence coverage below 10%, 12 displayed coverage between 10–20%, and 17 exhibited coverage above 20%. Only 5 MicroAltProts could not be reliably identified because no high- or medium-quality ion chromatograms were obtained for their unique peptides.

Then, based on the number of reliable peptides per protein, MicroAltProt identifications were subsequently classified into higher-confidence evidence (≥2 unique, non-nested peptides of length ≥9 amino acids meeting stringent thresholds, or 1 peptide with 100% coverage)—Tier 1A—, uncertain-confidence evidence (single unique peptide of length ≥9 amino acids that warrant orthogonal confirmation)—Tier 2A—, or low-confidence evidence (not meeting Tier 2A or 2B criteria but with high quality XICs) (Table 2). Therefore, 4 MicroAltProts were classified as Tier 1A (9.09%) and 20 (59.09%) as Tier 2A. Additionally, 14 MicroAltProts (31.81%) were classified as low-confidence evidence because the only unique peptide identified was shorter than nine amino acids (7 peptides with 7 amino acids and 7 peptides with 8 amino acids) (4), for which further validation is required to confirm their existence as bona fide MicroAltProts. However, 10 of these low-confidence MicroAltProts supported by short peptides (7–8 amino acids), exhibited high-quality XICs and were consistently detected in at least two independent extraction protocols (named low-confidence evidence). Although these low-confidence evidence-peptides fall just below the length required for Tier 2A classification, their reproducible detection across multiple extraction methods strengths confidence in their actual existence and warrants further validation.

Finally, we investigated the distribution of the 43 MicroAltProts identified by MS across the four predefined clusters, including their Tier classification. MicroAltProts analysis revealed a heterogeneous distribution, which displayed broad variability in the eight properties previously evaluated. Additionally, most of them showed very low MW and moderate flexibility, but high instability scores, consistent with a likely disordered nature (Fig. 7b-c). Furthermore, Cluster 2 was overrepresented, with 16 MicroAlprots (2 MicroAltProts in Tier 1A and 9 MicroAltProts in Tier 2A) belonging to this cluster, whereas cluster 0, marked by high hydrophobicity and α-helix content, was represented with only 5 MicroAltProt, probably because of a transmembrane location. In contrast, clusters 1 (9 MicroAltProts; 7 in Tier 2A) and 3 (13 MicroAltProts; 1 in Tier 1A and 4 Tier 2A) presented an equivalent number of MicroAltProts (Table 2).

### Structural-Context Examination of Experimentally Detected MicroAltProts

Finally, to gain insight into the potential functional roles of the 44 MicroAltProts with specific unique peptides (Table 2), we selected MicroAltProts —prioritized based on intensity, recurrence across detection methods, and cluster assignment— for structural characterization. For these MicroAltProts, we generated ab initio three-dimensional models using AlphaFold2, complemented by predictions of secondary structure, intrinsic disorder, and signal peptides or transmembrane helices (Fig. 8). Given the short length of many MicroAltProts and the limited evolutionary information available, AlphaFold2 models in this context should be interpreted as hypothesis-generating structural scenarios rather than definitive functional predictions.

**Fig. 8.**
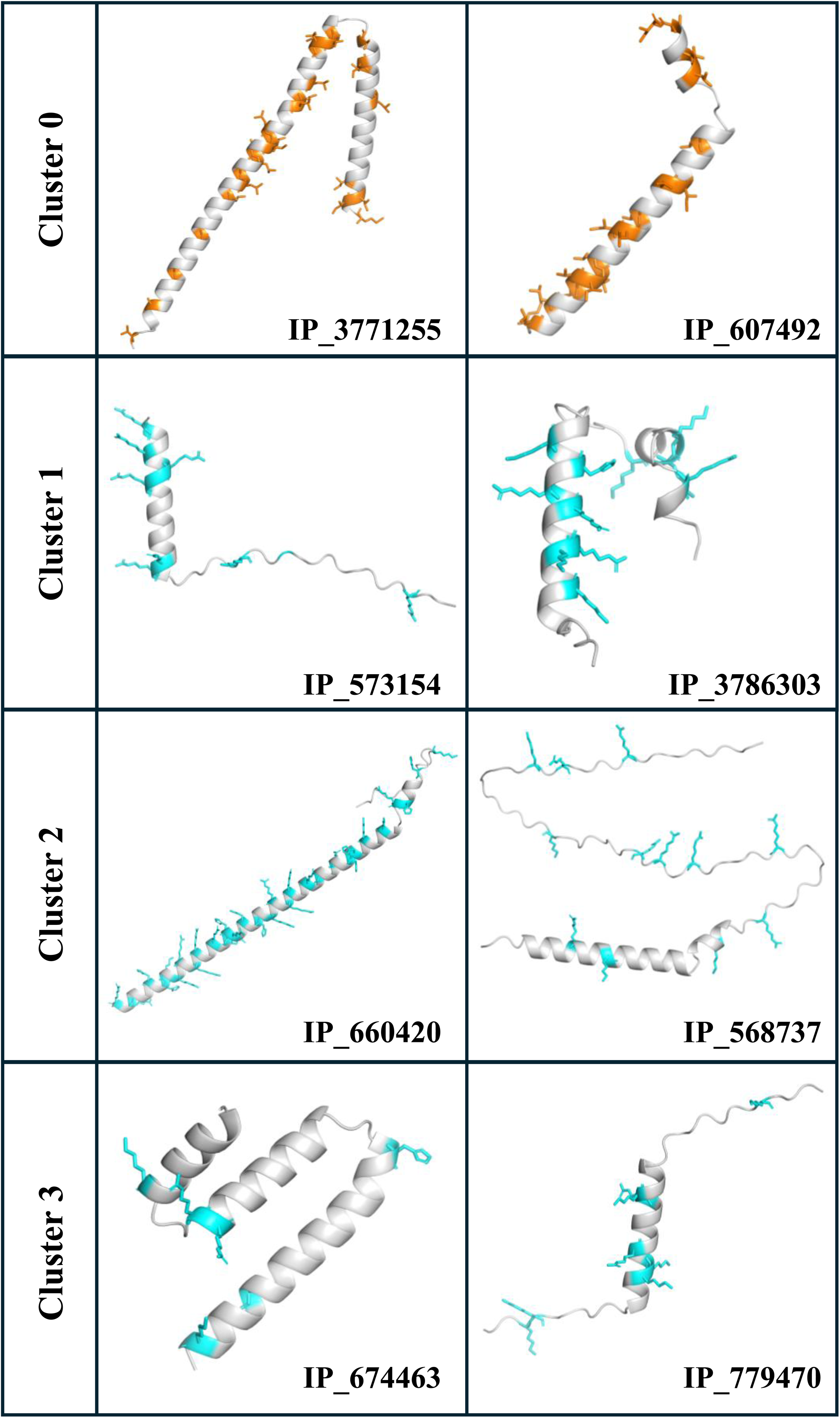
Cluster-based structural models of the verified alternative proteins by proteomics. AlphaFold2 models are shown for hypothesis generation; functional interpretations require orthogonal validation. Side chains of charged residues are shown in sticks and colored for alkaline (cyan), and hydrophobic (orange) residues when appropriate. Predicted structural model through AlphaFold, and contextual analysis of the alternative proteins validated through experimental evidence by proteomics. Two MicroAltProts belonging to each identified cluster were depicted.

Among the few hydrophobic verified MicroAltProts grouped in Cluster 0, IP_3771255 (88 aa; 23 unique peptides; GRAVY: 0.561) and IP_607492 (46 aa; 8 unique peptides; GRAVY: 0.743) each contained a predicted transmembrane α-helix. Their AlphaFold2 structural models further supported the presence of extended α-helical segments enriched in hydrophobic residues. This is consistent with the notion that, beyond their high hydrophobicity, these polypeptides may stably insert into cell membranes.

From Cluster 1, IP_573154 (42 aa; 17 unique peptides; Tier 2A; identified by five methods) and IP_3786303 (48 aa; 14 unique peptides; Tier 2A; identified by three methods) were very alkaline (pI: 12.0 and 10.4, respectively) and were predicted as unstable (76 and 64 instability index, respectively). Structural models for both proteins included α-helices with co-oriented positive-charged residues along at least three consecutive turns, a non-random configuration compatible with binding to polyanionic partners, such as DNA or RNA. This arrangement resembles the charged-aligned recognition helices of the Helix-Turn-Helix superfamily (43,44), suggesting a possible convergent structural relationship that may dictate their functional behavior.

Two of the relatively longer MicroAltProts, Cluster 2 IP_660420 (93 aa; 29 unique peptides; Tier 2A) and IP_568737 (101 aa; 27 unique peptides; Tier 2A), were also predicted to be alkaline (pI: 9.0 and 7.8, respectively). As in Cluster 1 proteins, both polypeptides displayed spatially aligned positive charges. Structural predictions indicated the presence of one α-helix with confident pLDDT values in both proteins. However, while IP_660420 was almost entirely α-helical, IP_568737 contained only a single helical segment within otherwise largely unfolded architecture.

Finally, Cluster 3 IP_674463 (61 aa; 10 unique peptides; Tier 1A) and IP_779470 (43 aa; 15 unique peptides; Tier 2A), also contained α-helical segments with some positively charged residues, but did not meet the criteria required to plausibly interact with nucleic acids, unlike the Cluster 1 and 2 cases.

It should be noted that AlphaFold2 tends to overestimate α-helix content in loop regions (45), a bias that may be accentuated due to the absence of multiple sequence alignments, as is the case for MicroAltProts. Nonetheless, in the cases described above, PSIPRED 4.0 independently corroborated a predominantly α-helical character of the segments involved.

Collectively, the strategy followed here allowed us to reveal a structurally and biochemically diverse landscape of MicroAltProts. Paradigmatic structures exemplify potential functional archetypes across clusters, ranging from highly flexible and disordered peptides with electrostatic patches, to helices resembling nucleic acid–binding modules, or hydrophobic domains compatible with transmembrane segments. Despite current limitations in modeling accuracy for very short sequences, these predictions strongly suggest that MicroAltProts may encode a wide repertoire of molecular behaviors with plausible implications in cellular regulation, signaling, and membrane dynamics. Together, these observations reinforce the notion that the alternative microproteome constitutes not only a vast reservoir of hidden coding potential but also a source of functionally relevant entities whose roles in physiology and disease warrant further experimental validation.

## Discussion

The human proteome, once thought to be essentially characterized (46–48), is being reshaped by the growing recognition of AltProts and, specifically, MicroAltProts, translated from unconventional ORFs (2,3,5,7,17). MicroAltprots represent a largely unexplored component of the human proteome, and most of their components are still functionally uncharacterized, with little or no information available regarding their biological roles, molecular interactions, cellular localization, or involvement in physiological and pathological processes. These entities, collectively termed the "ghost proteome", remain opaque due to limitations in annotation and methodological biases (49). In this study, we present a strategy combining computational curation and vanguardist MS-based proteomic high-sensitive Orbitrap Astral equipment to simultaneously analyze RefProts, isoforms, and AltProts in CRC cells, while assessing the influence of extraction protocols on AltProt detectability.

Our results demonstrate that extraction methodology markedly influences the detectability of the AltProt landscape. While RIPA-based protocols offered broader coverage—including membrane-associated and α-helical MicroAltProts—, HCl extraction favored disordered, and alkaline MicroAltProts. These biases align with the physicochemical nature of the extraction buffers, as RIPA preserves hydrophobic interactions *via* detergents, whereas acid-based solubilization denatures most proteins, favoring hydrophilic and disordered species. Notably, the complementary nature of these approaches underscores the importance of methodological diversity for comprehensive MicroAltProt profiling. Additionally, the enrichment of membrane-associated or highly hydrophobic features in certain clusters suggests that conventional protocols may overlook subsets of structurally atypical MicroAltProts.

In large-scale proteomic datasets, a limited number of false-positive identifications can disproportionately affect low-abundance protein groups, such as microproteins, leading to inflated confidence if class-specific FDR control is not applied. Accordingly, controlling the FDR specifically for unannotated proteins has been recommended to increase confidence in alternative protein identifications (14). While database searches restricted to MicroAltProts may improve class-specific FDR estimates, such approaches require careful evaluation to ensure that the identified peptides are not shared with canonical proteins. The presence of shared peptides may compromise or hinder the confident identification of specific proteins; therefore, focusing on unique peptides minimizes ambiguities increases the reliability of the results. In such cases, to avoid misassignment arising from shared peptides sequences (50), it is mandatory to assess whether peptides assigned to identified microproteins are also attributable to other annotated proteins, using specific tools as PeptideAtlas ProteoMapper,. As an alternative to these post hoc peptide-mapping approaches, we generated a comprehensive *in silico* catalogue of theoretical tryptic peptides from OpenProt, NextProt, and UniProt entries that can be directly integrated into proteomics pipelines, enabling rapid prioritization of class-specific unique peptides and high-confidence identification of AltProts and MicroAltProts. Beyond its application in the present study, this resource can be readily incorporated into alternative proteome workflows to improve peptide assignment, facilitate class-specific FDR strategies, and standardize AltProt identification across studies. Accordingly, when analyzing large databases such as the curated here OpenProt database, it is advisable to prioritize unique peptides for downstream interpretation instead of ProteinGroup quantification, as MicroAltProt-specific low-abundance peptides may be masked by high-abundance peptides shared with RefProts or isoforms.

To address these challenges and facilitate the interpretation of putative human alternative protein identifications in standard tryptic proteomics workflows, specific guidelines for the MS-based detection of MicroAltProts have recently been published as an extension of the Human Proteome Project Mass Spectrometry Data Interpretation Guidelines 3.0 (13,14). Based on these guidelines, here we have paid particular attention to FDR control and to the quality of the XICs associated with each peptide identified from MicroAltProt entries. We did not perform an independent database search using the MicroAltProt restricted dataset for separate FDR estimation to avoid potential misassignment arising from shared peptides. Additionally, the 43 MicroAltProts identified here were classified into Tier 1A, Tier 2A, or low-confidence evidence categories, and strong biological interpretations were restricted to Tier 1 or 2 candidates. In this study, a set of 14 MicroAltProts were classified as low-confidence evidence. However, these MicroAltProts were supported by high-quality XICs of unique peptides, which were only one or two amino acids shorter than the length threshold defined by current guidelines, and all of them but 3 were consistently detected in two or three independent extraction protocols. Although these candidates do not formally meet the criteria for Tier 2A classification, their reproducible detection and peptide-level quality would suggest an increased confidence in their existence and argue against their exclusion as false positives. These observations support the notion that, for MicroAltProts, a modest relaxation of peptide length requirements—when combined with stringent quality control and reproducibility criteria—may be biologically justified and warrants further investigation.

The small size of MicroAltProts, together with the flyability probability of their tryptic peptides, makes it often challenging to identify more than one or two unique tryptic peptides per protein by MS. Considering that in some cases even a single peptide can be difficult to detect, further validation using orthogonal or targeted MS approaches using synthetic peptides—such as parallel reaction monitoring (PRM)—would substantially strengthen the confidence in proteomics-based identifications. Notably, in this study we also provide a curated list of unique tryptic peptides and their predicted MS flyability, which could serve as a valuable resource for future PRM validation experiments, which is particularly relevant given the lack of antibodies against specific MicroAltProts.

By curating the OpenProt database and excluding entries with homology to canonical proteins or exceeding 200 amino acids, we focused here on identifying true actual MicroAltProts. Subsequent clustering based on eight general physicochemical features revealed four distinct natural groups of MicroAltProts. Experimental detection of 43 MicroAltProts confirmed that members from each cluster could be recovered and structurally profiled.

Additionally, 19 of these MicroAltProts lacked previous MS evidence and 13 were identified in one unique study with one unique peptide, highlighting both the novelty of our dataset and the potential biological significance of these newly detected microalternative proteins. These groups likely represent functional archetypes: small, hydrophobic, and compositionally simple α-helical proteins (Cluster 0), disordered and unstable MicroAltProts (Cluster 1), larger but unstable high-content β-turn proteins (Cluster 2), and intermediate amphipathic MicroAltProts with balanced properties (Cluster 3). This classification not only provides a rational framework to guide functional hypotheses but also most likely reflects possible evolutionary constraints or cellular niche specialization. Computational clusters mirror the actual biochemical diversity in experimentally validated MicroAltProts by MS, except for the most hydrophobic, which was probably underrepresented because of membrane association and/or lower abundance than the other ones.

One of the main goals of this study was to challenge the historical exclusion of AltProts from classical proteomic and functional genomics workflows (7,17,49). To explore their potential roles, we systematically characterized their amino acid composition, predicted structural features, subcellular localization, and functional signatures. Because most MicroAltProts are too short to contain large, conserved domains, their biological activity is likely mediated by short functional elements. Accordingly, we investigated the prevalence of SLiMs together with conserved PROSITE signatures that provide clues to specific structural or functional properties. Both analyses revealed an enrichment of motifs associated with protein-protein interactions, post-translational regulation, and subcellular targeting, which suggest that despite their small size, MicroAltProts may primarily function as regulatory molecules or interaction hubs rather than enzymes with complex catalytic domains. In parallel, the enrichment of residues associated with intrinsically disordered regions and post-translational regulation, particularly serine and cysteine, suggests that these proteins may participate in signaling and regulatory pathways despite their small size, while the unexpected enrichment of tryptophan further supports the existence of a unique compositional signature. Consistent with these observations, DeepLoc predictions indicated that MicroAltProts are not restricted to a single cellular compartment but are distributed across diverse subcellular locations, including the cytoplasm, nucleus, endomembrane system, and extracellular secretory pathway. The presence of predicted signal peptides and transmembrane domains in a subset of proteins further supports their potential association with membrane trafficking and cell-cell communication. In addition, the predicted occurrence of amphipathic helices, intrinsically disordered regions, transmembrane segments, and nucleic acid-binding architectures is consistent with potential roles in membrane organization, intracellular trafficking, gene regulation, stress responses, or RNA metabolism (51). These three independent layers of evidence converge to support the view that MicroAltProts are biologically meaningful proteins rather than represent biological noise or truncated versions of canonical proteins. Although these predictions require experimental validation, together they provide a first functional framework suggesting that MicroAltProts represent a specialized and previously overlooked layer of the proteome with potential implications in tumor biology and other physiological and pathological processes (52–57).

This study has several limitations. First, our experimental set-up was restricted to a single CRC cell line (KM12C), which may not capture the diversity of AltProts and MicroAltProts in other cellular scenarios. Second, while *in silico* structural predictions provide valuable insights, they require orthogonal validation through biochemical or biophysical methods, such as for example cross-linking MS (58). Third, a notable observation is the large discrepancy between the number of MicroAltProts annotated in OpenProt (183,937 sequences ≤200 amino acids) and the 43 MicroAltProts experimentally validated in this study. This gap reflects both biological and technical constraints. While the ≤200 amino acid threshold enables an intentionally inclusive representation of the alternative proteome, many predicted altORFs are unlikely to be expressed, stable, or detectable specific cellular contexts. Moreover, proteins at the lower end of this size spectrum often yield few or no unique tryptic peptides, further limiting detectability in standard bottom-up proteomics workflows. These factors, together with conservative evidence thresholds required to control false positives in large search spaces, inevitably result in a relatively small but high-confidence set of experimentally supported MicroAltProts. Future studies should incorporate alternative proteases (i.e. GluC or chymotrypsin) to increase unique peptide yield, MicroAltProt enrichment or fractionation strategies, targeted validation approaches such as PRM or SRM for prioritized candidates, and integration with orthogonal datasets (e.g., ribosome profiling) to reduce search space and strengthen biological confidence. Fourth, functional roles of identified MicroAltProts remain hypothetical at this stage. Functional characterization will further require stable overexpression or CRISPR-based perturbation strategies, as well as consideration of post-translational modifications, which were not explored here but may be critical for MicroAltProt function and stability.

Nevertheless, our findings have broad implications as similar pipelines could be extended to other cell types, tissues, or disease models, including neurodegenerative disorders, where dysregulated non-canonical translation and stress-adaptive proteomes have been described (59,60). Furthermore, the limited sequence similarity of MicroAltProts to known proteins makes them attractive biomedical candidates such as tumor-specific targets or antigens, or modulators of immunogenicity—offering exciting diagnostic or therapeutic opportunities.

Beyond these immediate findings, our study paves the way for systematic exploration of MicroAltProts as potential contributors to human disease. By providing a curated database together with a property-based framework for their classification, we establish a foundation for subsequent studies aimed at testing functional hypotheses, linking structural archetypes with biological processes, and extending these analyses to tumor samples, patient-derived models, and other disease contexts. Importantly, because of their limited sequence similarity to canonical proteins, traditional annotation or homology-based tools fall short in predicting their roles. Instead, clustering by physicochemical properties offers a rational strategy to uncover latent structure–function relationships within the alternative proteome. This paradigm not only enhances interpretation of large-scale proteomic data but also expands our view of the protein universe, where short, diverse, and non-canonical polypeptides may exert tightly regulated and context-specific functions. Despite the remaining challenges—such as validation, functional assays, and the integration of post-translational modifications—our results highlight that diversity itself constitutes a meaningful biological signal. The alternative microproteome may therefore provide novel entry points to dissect human disease mechanisms and develop therapeutic interventions. Indeed, representative members from each cluster already display archetypal features that should influence cellular localization, stability, and molecular interactions. Together, these observations provide a framework to explore the involvement of MicroAltProts in pathological contexts—illustrated here with CRC cells as a model—where non-canonical translation products may contribute to dysregulated signaling, stress adaptation, or cellular plasticity.

## Conclusions

We present in this study a scalable framework combining bioinformatic curation and high-resolution MS to simultaneously analyze RefProts, isoforms, and MicroAltProts. By integrating database refinement, physicochemical clustering, and comparative extraction workflows, we were able to uncover structurally distinct classes of MicroAltProts. Our results demonstrate that the extraction method significantly influences AltProt detectability, with RIPA-based buffers enabling broader recovery. Our study provides a systematic approach to explore the diversity and potential functionality of non-canonical proteins across biological systems. Importantly, the identification and classification of MicroAltProts in a CRC context highlights their relevance in disease biology and suggests their potential as novel regulatory elements or biomarker candidates. Collectively, this study provides not only a workflow for confident MicroAltProt identification but also practical resources—including a curated peptide catalogue and a structure-based classification framework—that can facilitate future investigations of the alternative proteome across biological systems.

## Supplementary data

The online version contains supplementary material available at

## Acknowledgements

Not applicable.

## Authors’ contributions

Conceptualization, A.M-C., AJ. M-G., and R.B.; methodology, A.M-C., A. P-G., AJ. M-G., and R.B.; investigation, A.M-C., A. P-G., AJ. M-G., and R.B.; writing-original Draft, A.M-C., AJ. M-G., and R.B.; writing-review and editing, A.M-C., A. P-G., AJ. M-G., and R.B.; resources, AJ. M-G., and R.B.; supervision, A.M-C. and R.B.; and funding acquisition, AJ. M-G. and R.B. All authors have read and approved the manuscript.

## Conflict of interest

The authors declare no competing interests.

## Funding

The financial support of Grants PID2022-140307OB-I00 and PID2023-151514OB-I00 funded by MCIN/AEI/10.13039/ 501100011033 and by “ERDF A way of making Europe” to R.B. and AJ.M-G., respectively, and PI23CIII/00027 grant from the AES-ISCIII program cofounded by FEDER funds to R.B.

## Data Availability

All data generated or analyzed during this study are included in this published article and its supplementary information files. The Mass Spectrometry data were deposited to the ProteomeXchange Consortium via the PRIDE partner repository with the dataset identifier PXD066336. All MicroAltProt-supporting spectra are referenced by Universal Spectrum Identifiers in the Supplementary Information.

## Declarations

### Ethics approval and Consent to participate

Not applicable.

### Consent for publication

All authors consent for publication.

### Abbreviations

AltProts,: alternative proteins
HCl,: Hydrochloric acid
MicroAltProts,: Alternative microproteins, alternative proteins less than 200 amino acids in length
RefProts,: reference proteins
RIPA,: Radioimmunoprecipitation assay buffer
ORF,: Open reading frame
sORF,: small ORF
AltORF,: alternative ORF
Ribo-Seq,: ribosome profiling
MS,: mass spectrometry
LC-MS/MS,: liquid chromatography tandem
MS RT,: retention time
CVs,: coefficients of variation
DIA,: data independent acquisition
pI,: isoelectric point
FDR,: false discovery rate
CRC,: Colorectal cancer
USI,: universal spectrum identifiers
XIC,: extracted ion chromatogram
HTH,: Helix-Turn-Helix

## Legend to the Supplementary Figures

**Fig. S1.** Quality control assessment of the protein extraction protocols. *A.* Comassie blue and silver staining of protein extracts from KM12C cells used for LC-MS/MS. *B*, Summarized number of Peptides (Top) and Protein groups (Bottom) identified per run (colored bars) and in the global experiment after cross-run integration (black bar). Numbers above the bars indicate the estimated number of false positive identifications after applying the corresponding FDR thresholds (FDR ≤0.01 at the global experiment level; FDR ≤0.05 at run-wise level). Total ion current (TIC) profiles (*C*) and pump pressure profiles (*D*) across the chromatographic gradient for each replicate of each extraction method indicate comparable injection performance and overall MS stability.

**Fig. S2.** Overlap and distribution of the proteins identified across extraction methods. *A*, Venn diagram of common and differential RefProts, isoforms, and AltProts identified per extraction method. A consistent core of 172 AltProts, 638 isoforms, and 1272 RefProts proteins was observed to be shared among methods, with additional method-specific detections that were mitigated in RIPA-based methodologies. *B*, Isoform and RefProt identifications among the four different extraction methods. The depicted pie charts show the number of identified Isoforms or RefProts in each method-specific subset. *C*, Ranked abundance distribution of KM12C proteins across extraction methods. A significant shift in dynamic range and detection depth was observed for the HCl and Filtered samples in comparison to RIPA-based protocols. *D,* Ranked abundance distribution of proteins according to their classification (RefProts, Isoforms, or AltProts) across extraction methods in the KM12C cells.

**Fig. S3.** Influence of extraction protocol on the physicochemical properties of identified proteins. *A*, Violin plots comparing molecular weight (MW, kDa), hydrophobicity score (GRAVY score), and isoelectric point (pI) of proteins extracted with four different protocols: HCl (blue), RIPA (green), RIPA–Chloroform (pink), and RIPA followed by fractionation into High- (Non-Filtered, Maroon) and low- (Filtered, purple) molecular weight proteins. *B*, Density plots for the same properties (MW, GRAVY, and pI), showing the overall distribution trends of each parameter across the different extraction methods.

**Figure S4.** *In silico* characterization of DeepLoc-derived MicroAltProt localization clusters. Heatmaps displaying the relative enrichment of predicted subcellular compartments (left), membrane association classes (middle), and localization signals (right) across the four MicroAltProt clusters identified by DeepLoc. Colors represent normalized Z-scores, with red indicating relative enrichment and blue indicating relative depletion.

**Fig. S5.** Extraction ion chromatograms of the precursor ions from unique peptides of MicroAltProts identified in each sample by LC-MS/MS.

## Notes

### Competing Interest Statement

The authors have declared no competing interest.

