## Supplementary Figures for "Mapping the Human Ghost Proteome: Classification and Experimental Detection Biases in the Identification of Alternative Microproteins": Figure S1.pdf

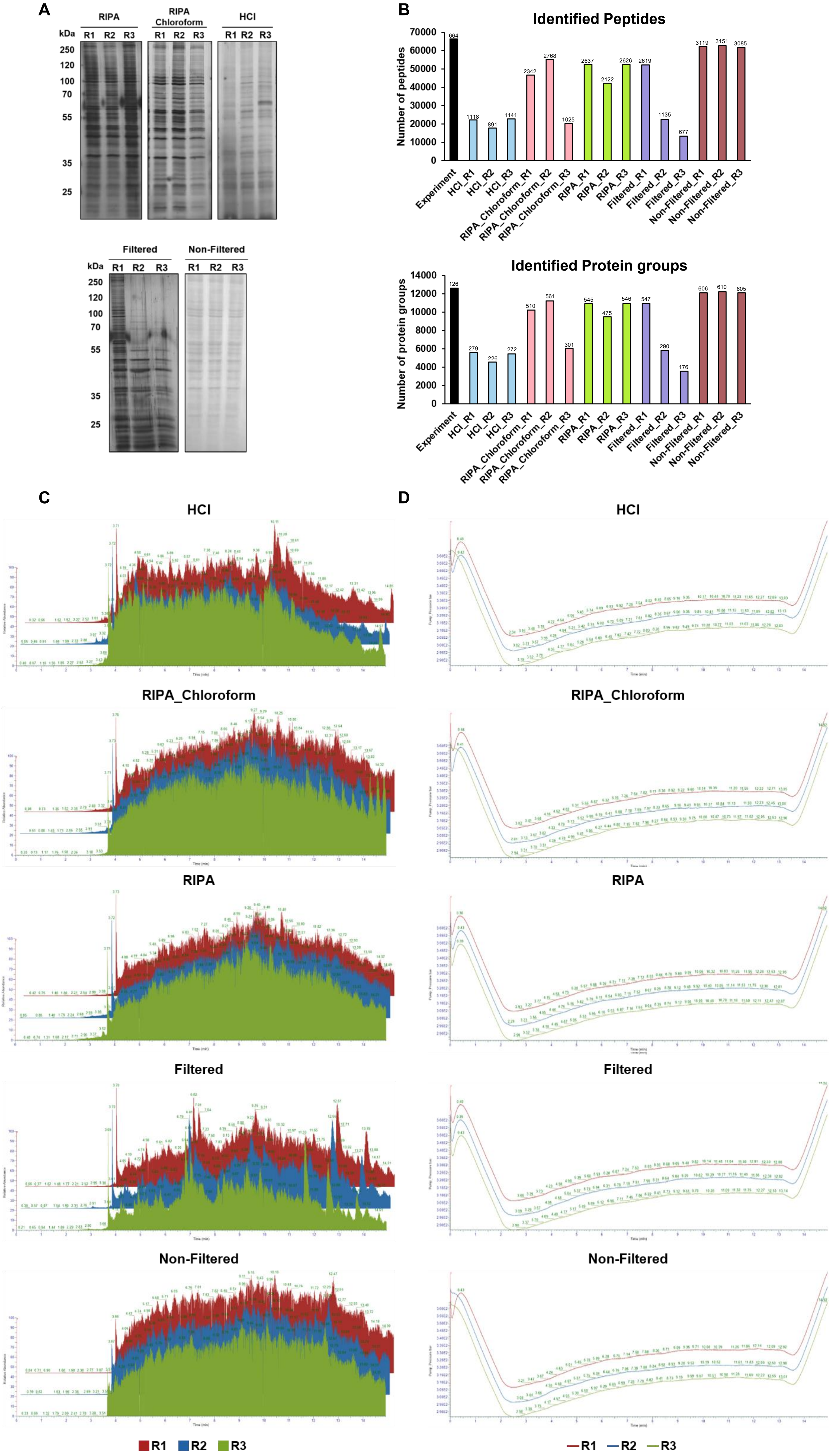


**Fig. S1. Quality control assessment of the protein extraction protocols.** *A*. Comassie blue and silver staining of protein extracts from KM12C cells used for LC-MS/MS. *B*, Summarized number of Peptides (Top) and Protein groups (Bottom) identified per run (colored bars) and in the global experiment after cross-run integration (black bar). Numbers above the bars indicate the estimated number of false positive identifications after applying the corresponding FDR thresholds (FDR ≤0.01 at the global experiment level; FDR ≤0.05 at run-wise level). Total ion current (TIC) profiles (*C*) and pump pressure profiles (*D*) across the chromatographic gradient for each replicate of each extraction method indicate comparable injection performance and overall MS stability.
