## Supplementary Figures for "Mapping the Human Ghost Proteome: Classification and Experimental Detection Biases in the Identification of Alternative Microproteins": Figure S2.pdf

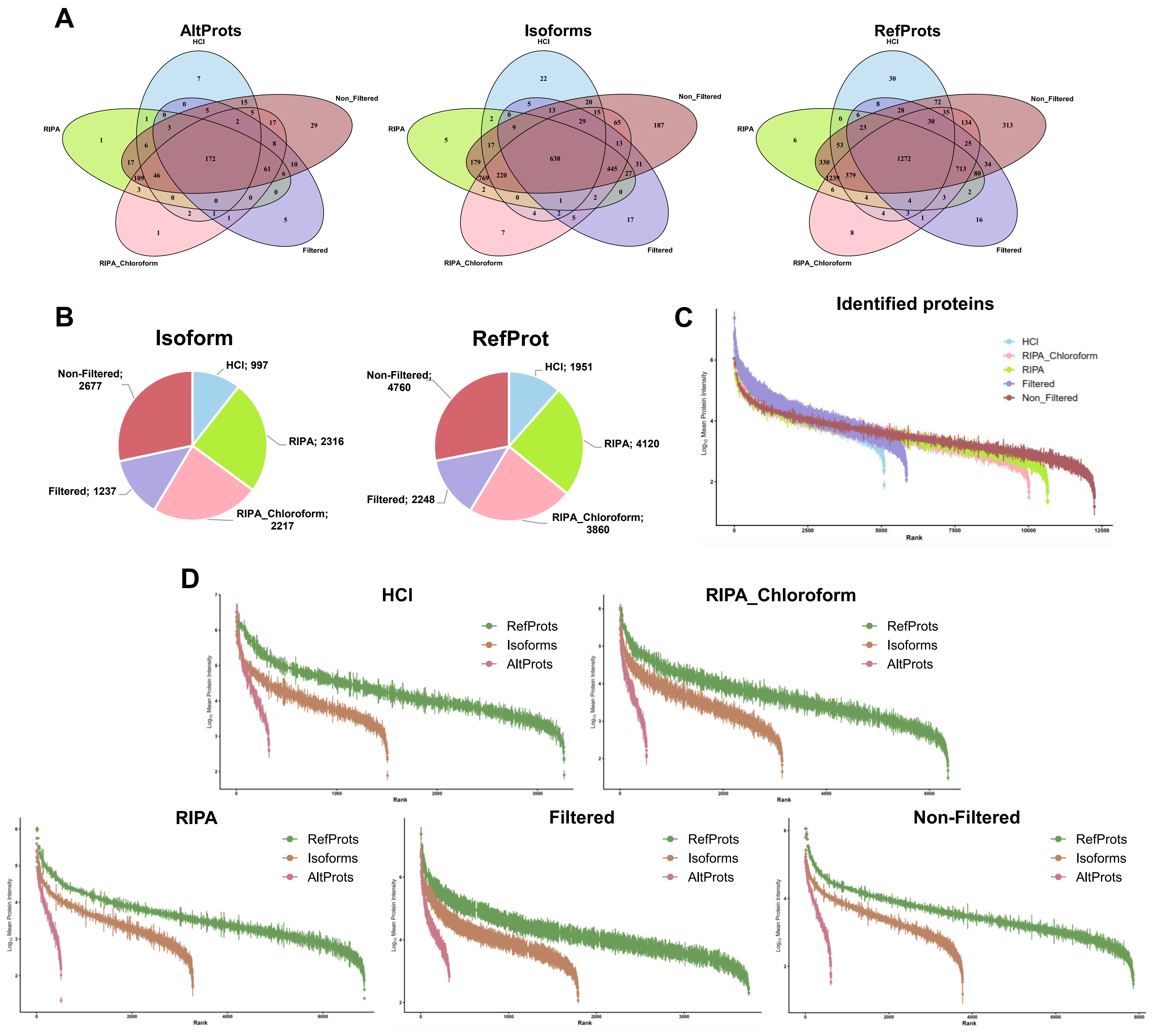


**Fig. S2.** Overlap and distribution of the proteins identified across extraction methods. *A*, Venn diagram of common and differential RefProts, isoforms, and AltProts identified per extraction method. A consistent core of 172 AltProts, 638 isoforms, and 1272 RefProts proteins was observed to be shared among methods, with additional method-specific detections that were mitigated in RIPA-based methodologies. *B*, Isoform and RefProt identifications among the four different extraction methods. The depicted pie charts show the number of identified Isoforms or RefProts in each method-specific subset. *C*, Ranked abundance distribution of KM12C proteins across extraction methods. A significant shift in dynamic range and detection depth was observed for the HCl and Filtered samples in comparison to RIPA-based protocols. *D,* Ranked abundance distribution of proteins according to their classification (RefProts, Isoforms, or AltProts) across extraction methods in the KM12C cells.
