## Supplementary Figures for "Mapping the Human Ghost Proteome: Classification and Experimental Detection Biases in the Identification of Alternative Microproteins": Figure S3.pdf

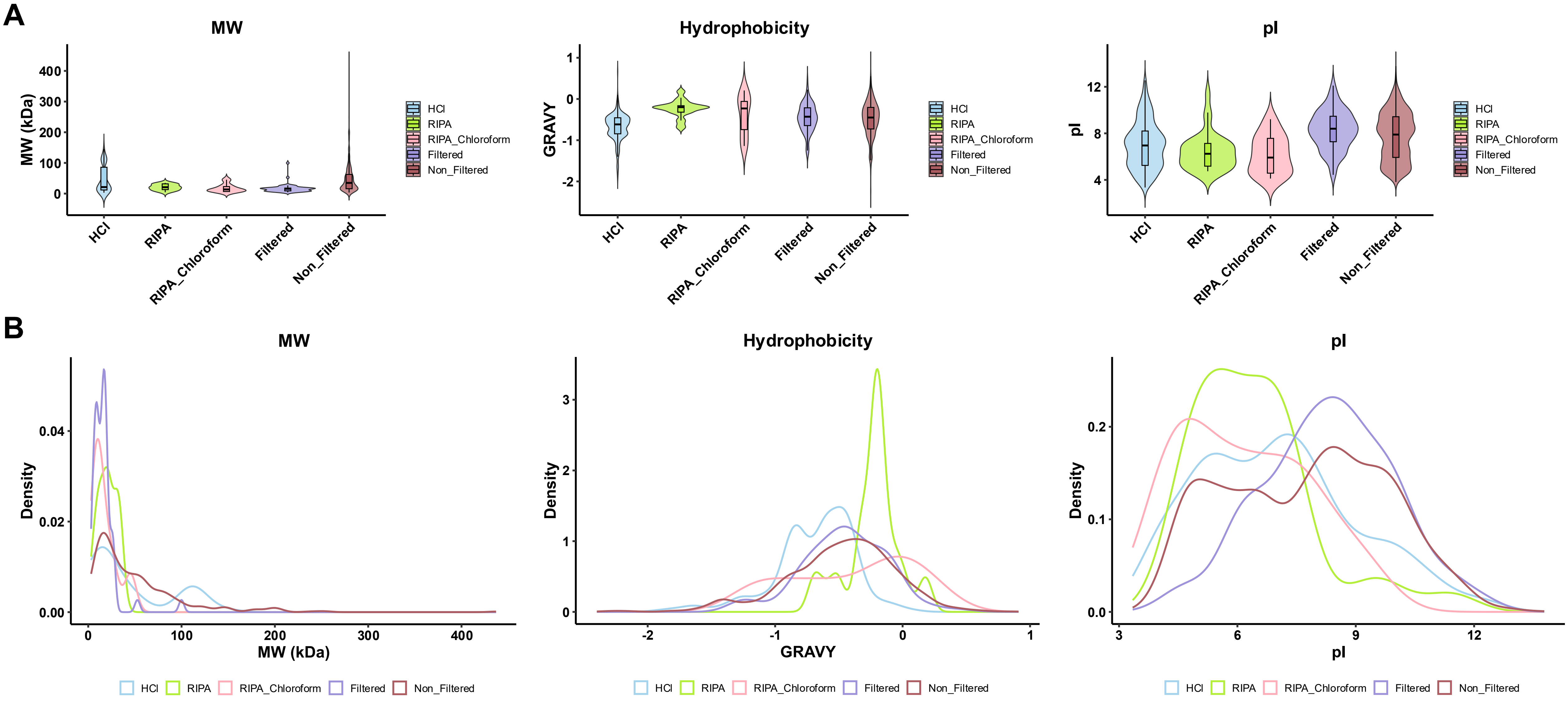


**Fig. S3. Influence of extraction protocol on the physicochemical properties of identified proteins.** *A*, Violin plots comparing molecular weight (MW, kDa), hydrophobicity score (GRAVY score), and isoelectric point (pI) of proteins extracted with four different protocols: HCl (blue), RIPA (green), RIPA–Chloroform (pink), and RIPA followed by fractionation into High- (Non-Filtered, Maroon) and low- (Filtered, purple) molecular weight proteins. *B*, Density plots for the same properties (MW, GRAVY, and pI), showing the overall distribution trends of each parameter across the different extraction methods.
