## Supplementary Figures for "Mapping the Human Ghost Proteome: Classification and Experimental Detection Biases in the Identification of Alternative Microproteins": Figure S4.pdf

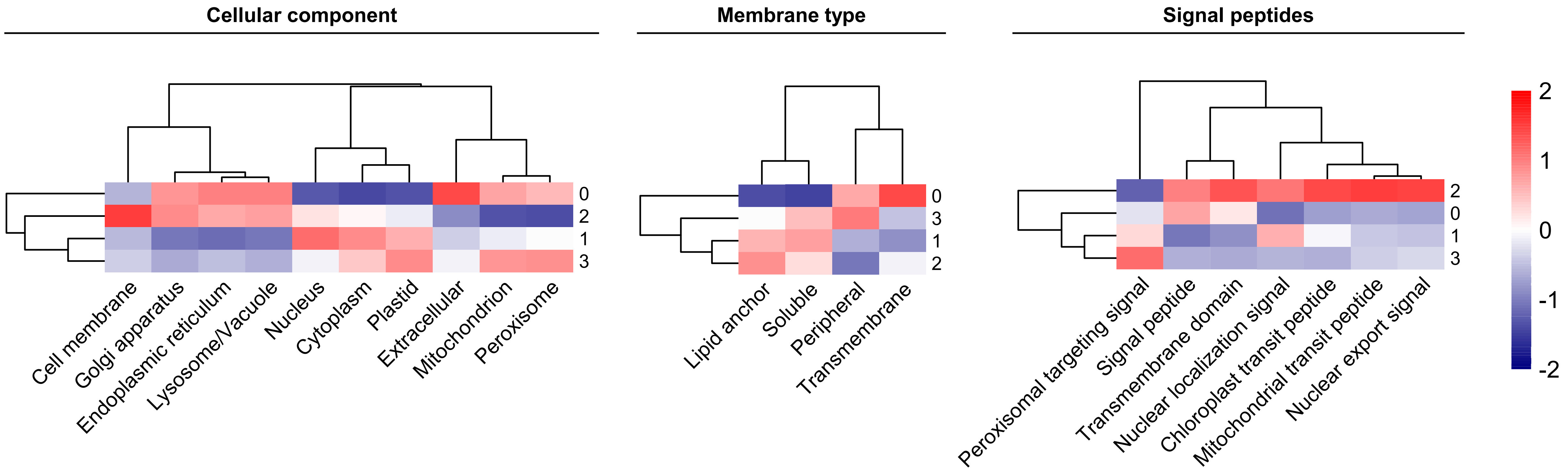


**Figure S4.** *In silico* characterization of DeepLoc-derived MicroAltProt localization clusters. Heatmaps displaying the relative enrichment of predicted subcellular compartments (left), membrane association classes (middle), and localization signals (right) across the four MicroAltProt clusters identified by DeepLoc. Colors represent normalized Z-scores, with red indicating relative enrichment and blue indicating relative depletion.
