## Supplementary Figures for "Mapping the Human Ghost Proteome: Classification and Experimental Detection Biases in the Identification of Alternative Microproteins": Figure S5_REDUCED.pdf

### mzspec_PXD066336_IP_762755_AATAAPLAQSSSSR

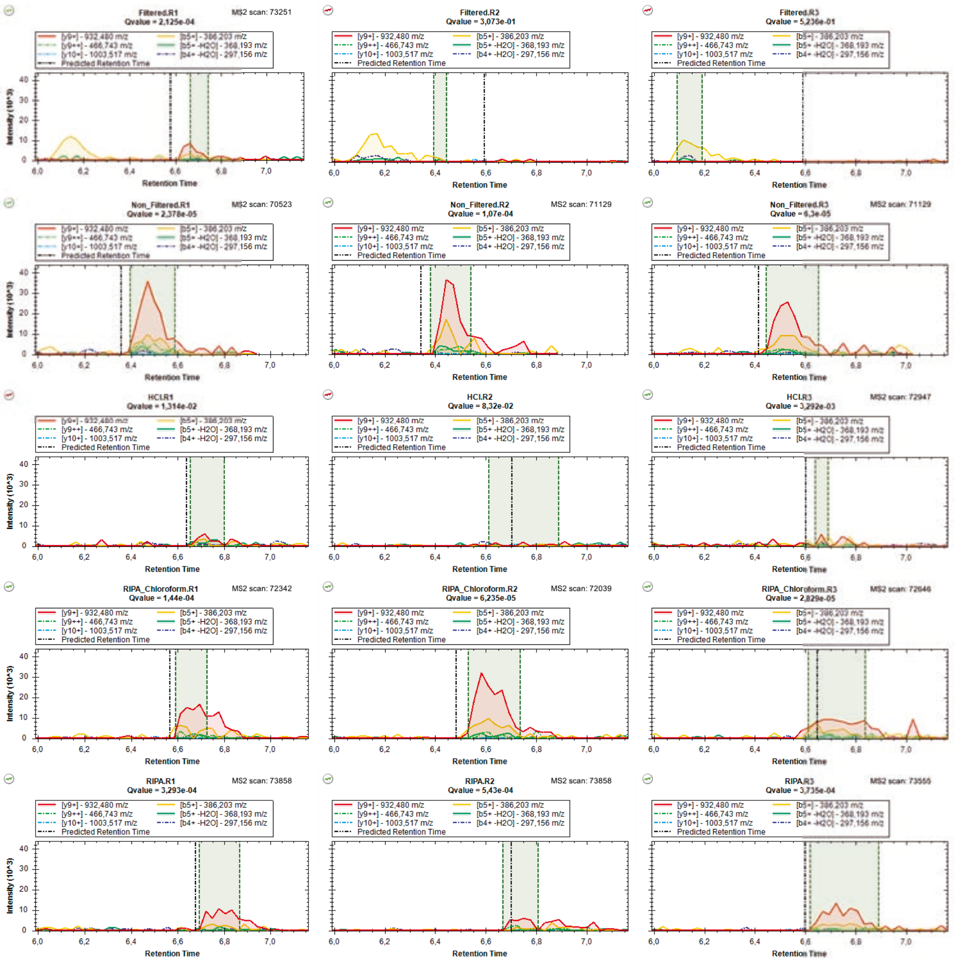

### mzspec_PXD066336_IP_707397_AIEEQVAVIYAGVR_Charge_3

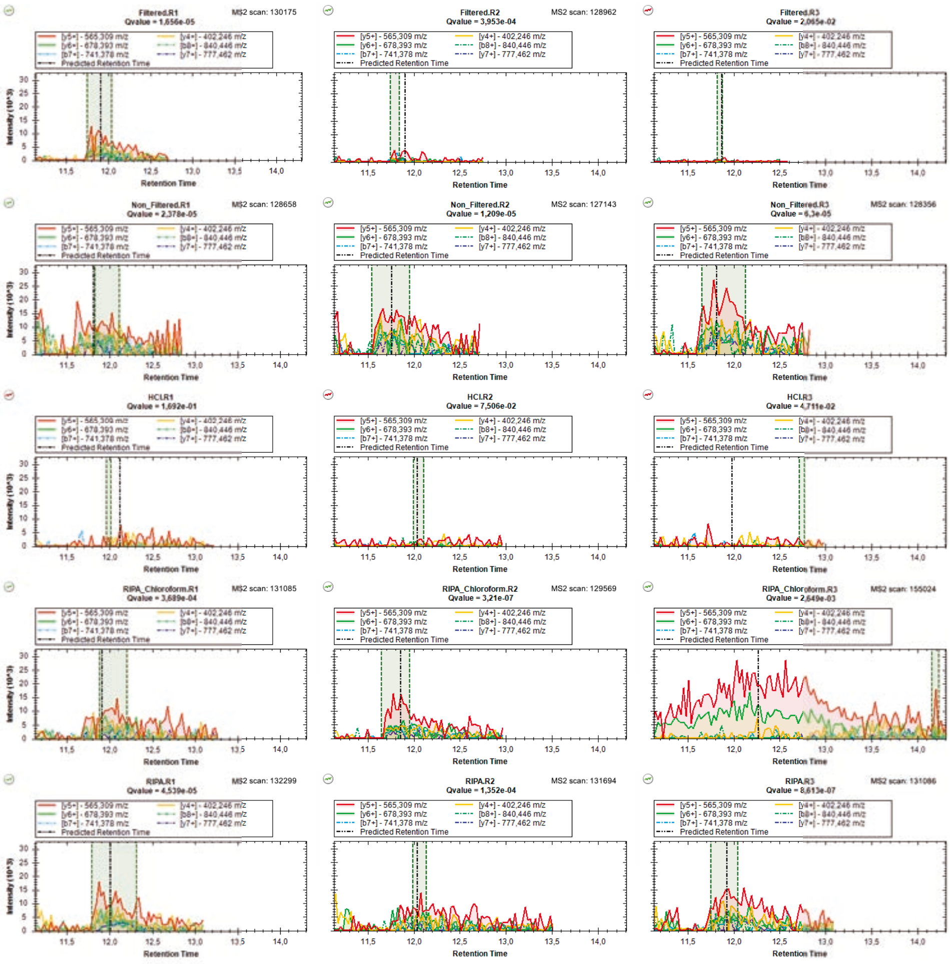

### mzspec_PXD066336_IP_707397_AIEEQVAVIYAGVR_Charge_2

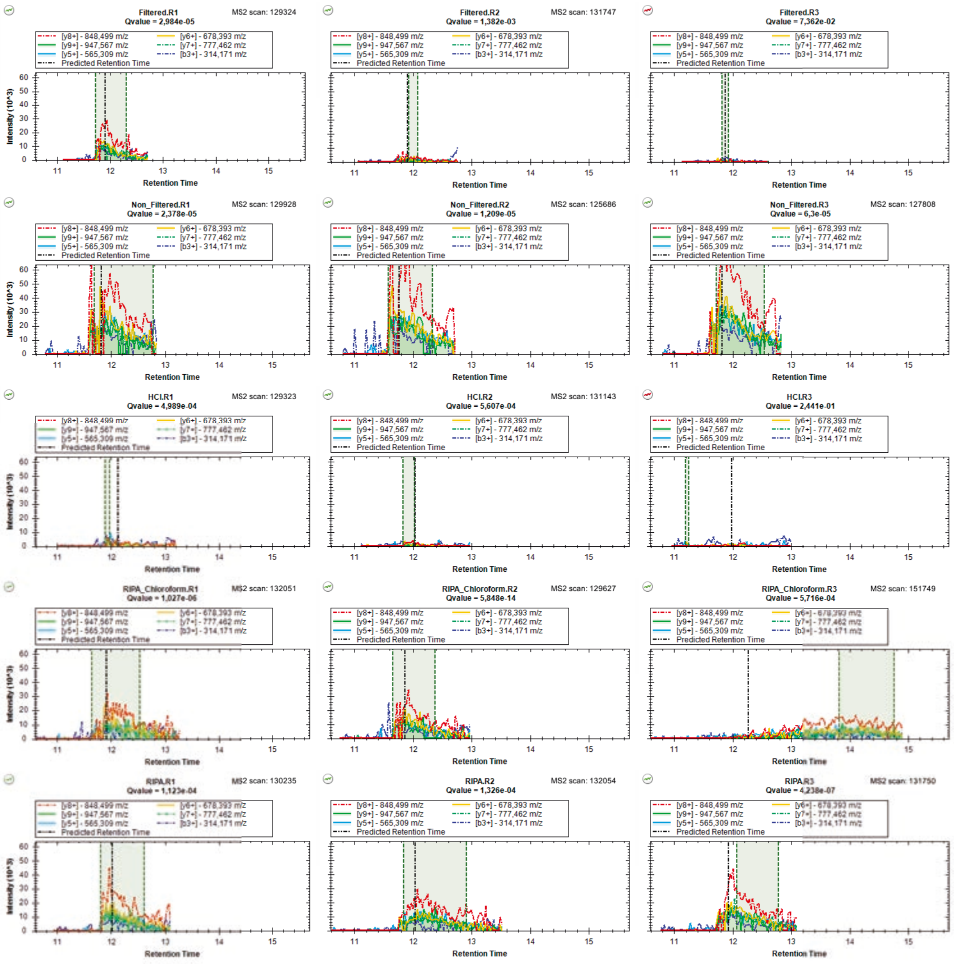

### mzspec_PXD066336_IP_674463_ALDALADGGVK

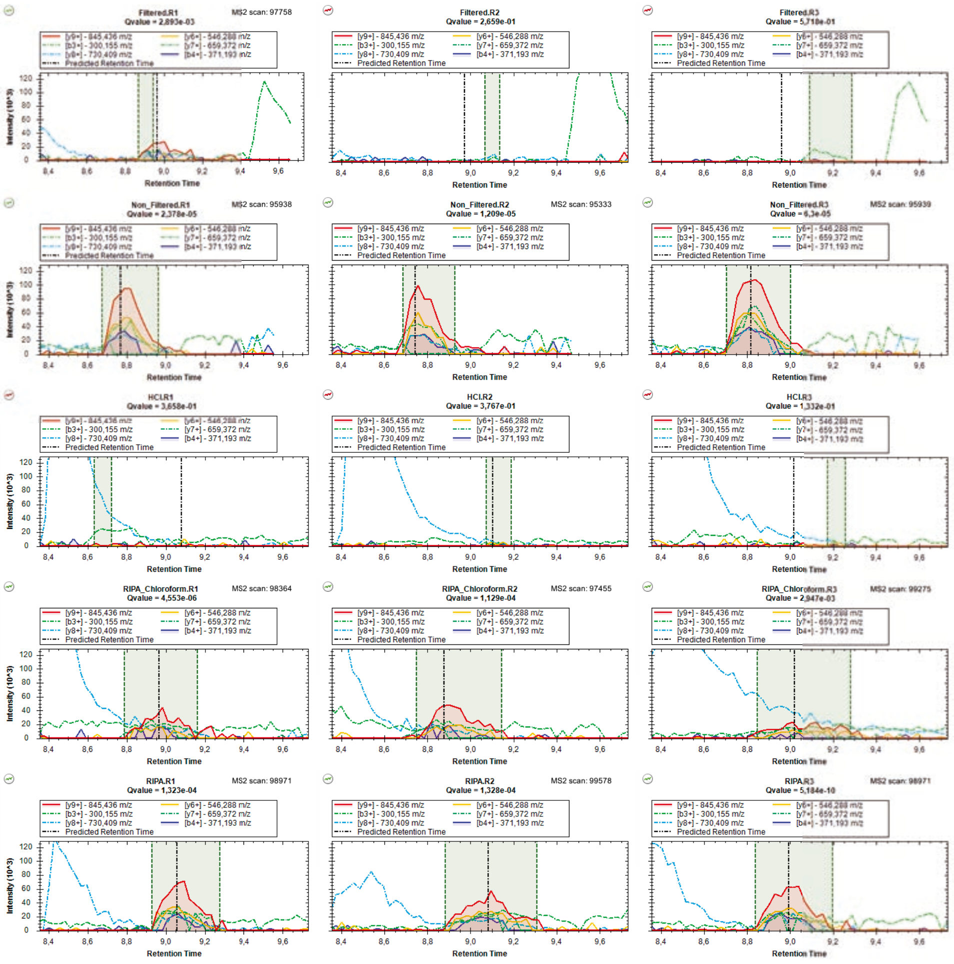

### mzspec_PXD066336_IP_587733_ASGQELPPPAPVPRPPAAPNPAPR

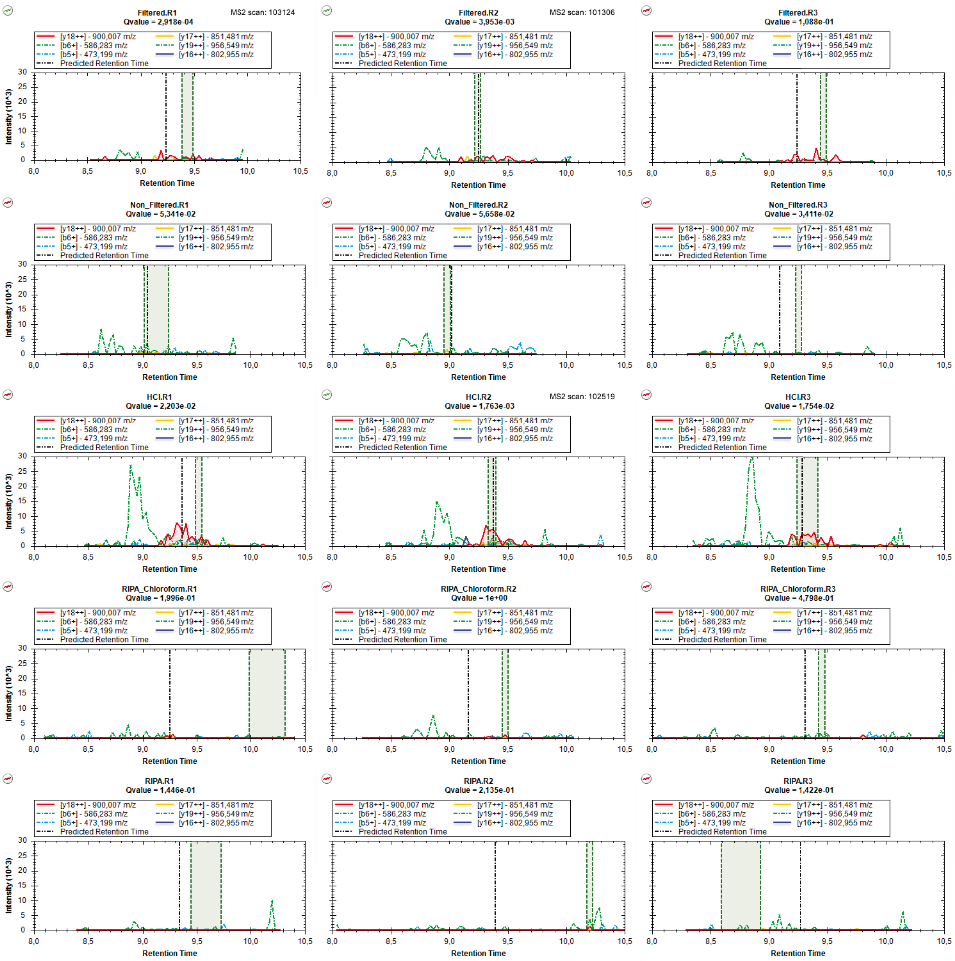

### mzspec_PXD066336_IP_566433_AVNYVQHNR

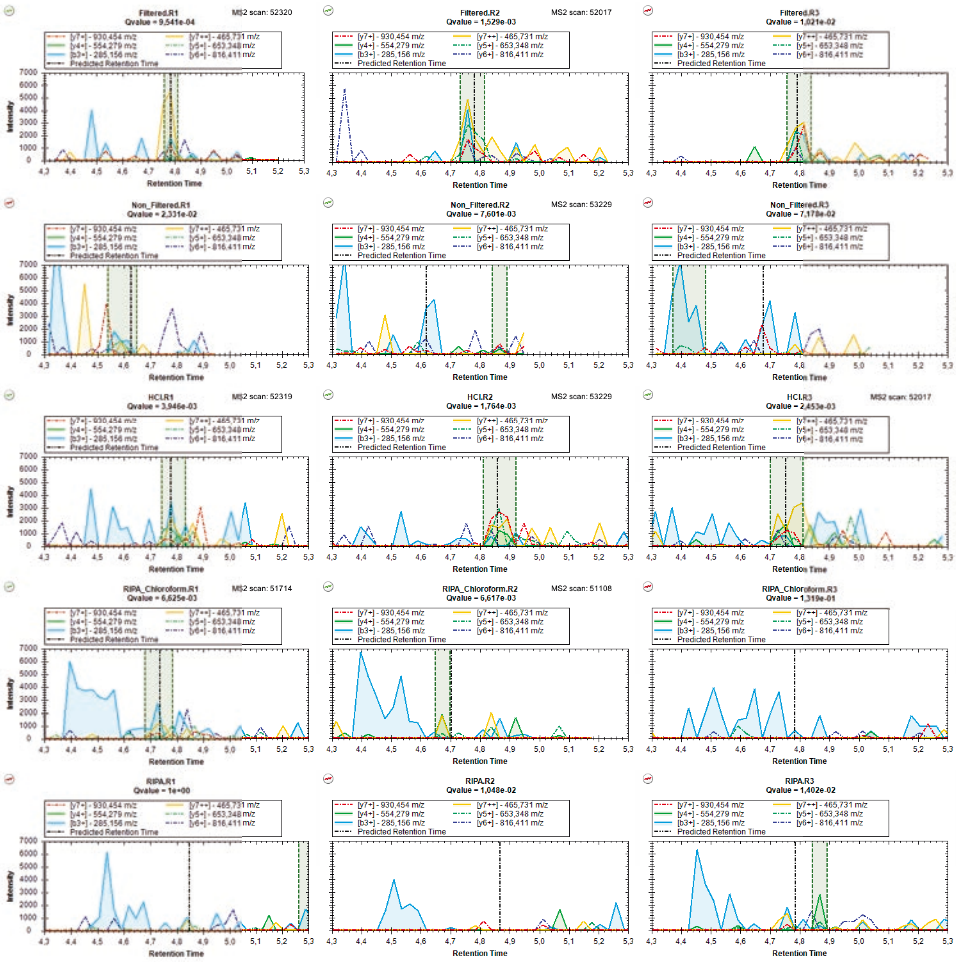

### mzspec_PXD066336_IP_674332_DAEAGQLADGAGRPLAEGPRPHLVR

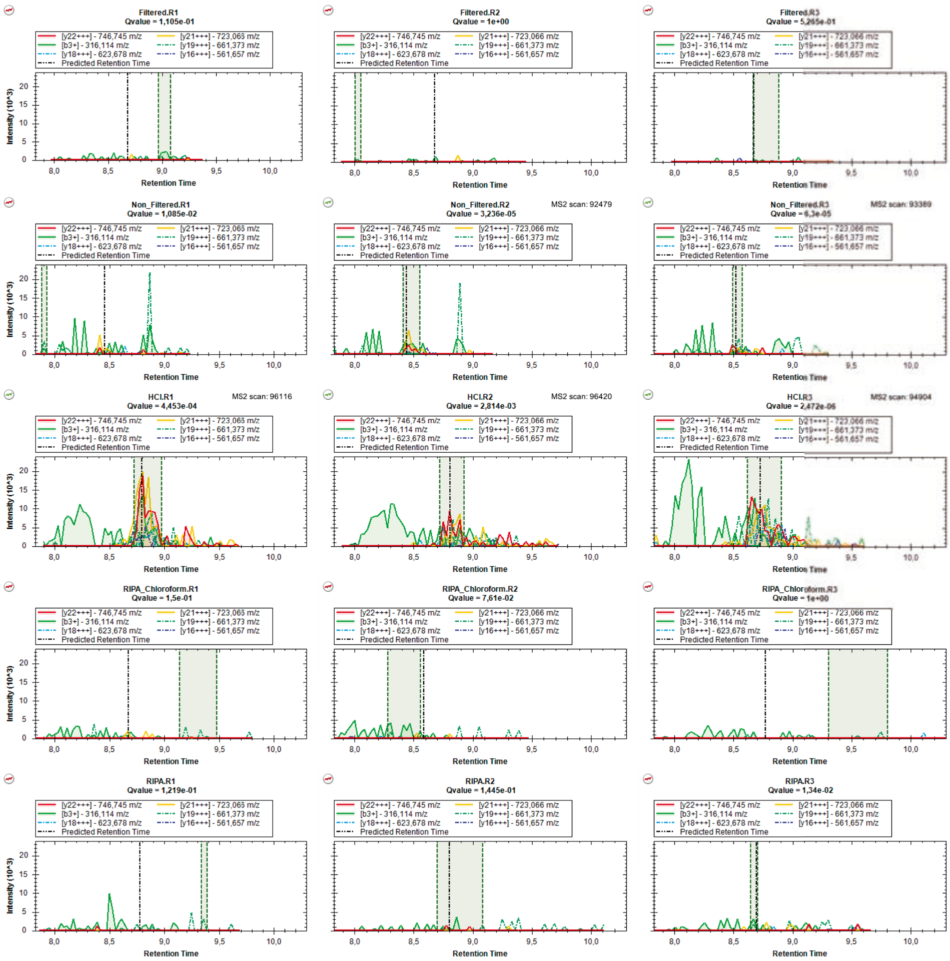

### mzspec_PXD066336_IP_3431818_DILQFYVK

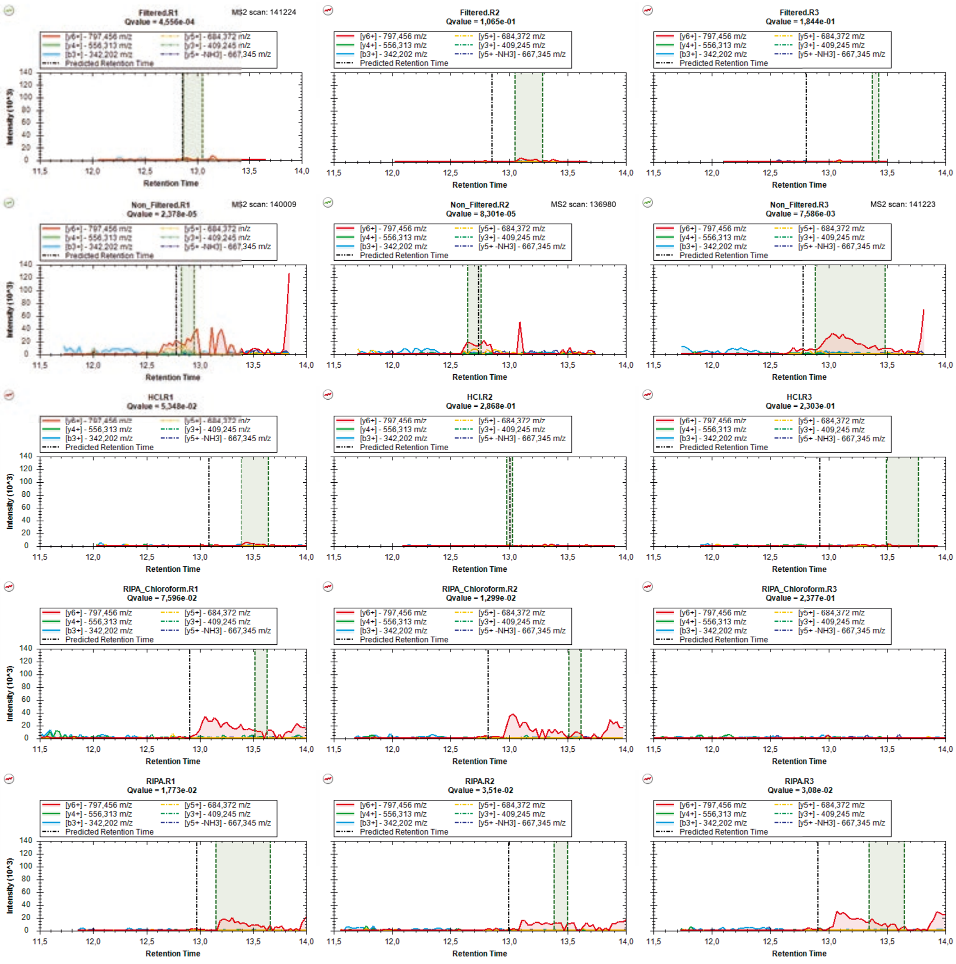

### mzspec_PXD066336_IP_747090_DLLIVLSR

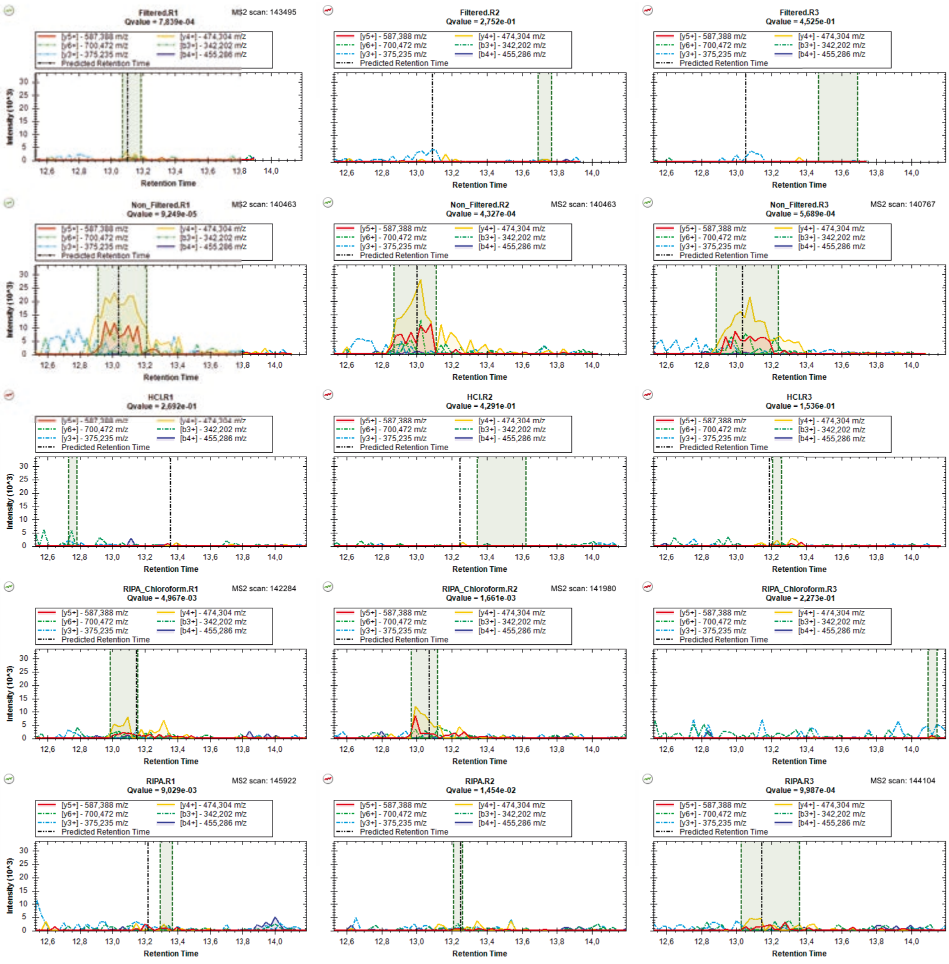

### mzspec_PXD066336_IP_607492_DLNELKK_Charge_2

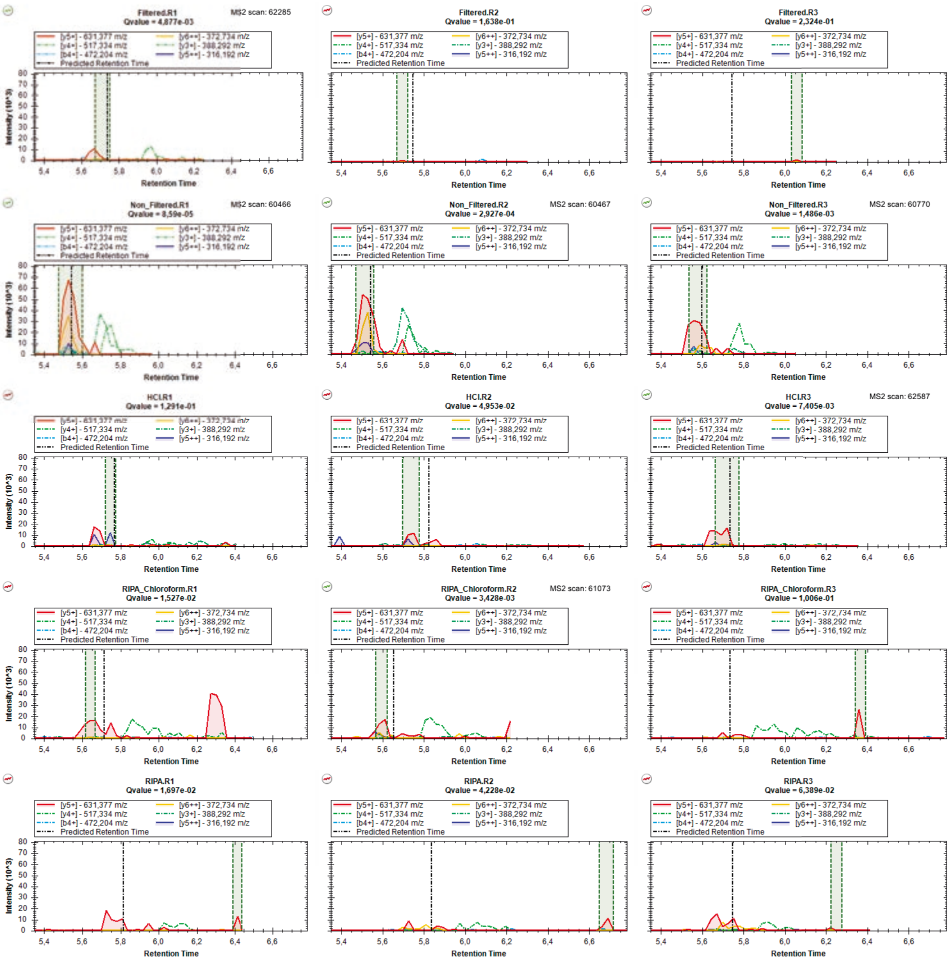

### mzspec_PXD066336_IP_607492_DLNELKK_Charge_1

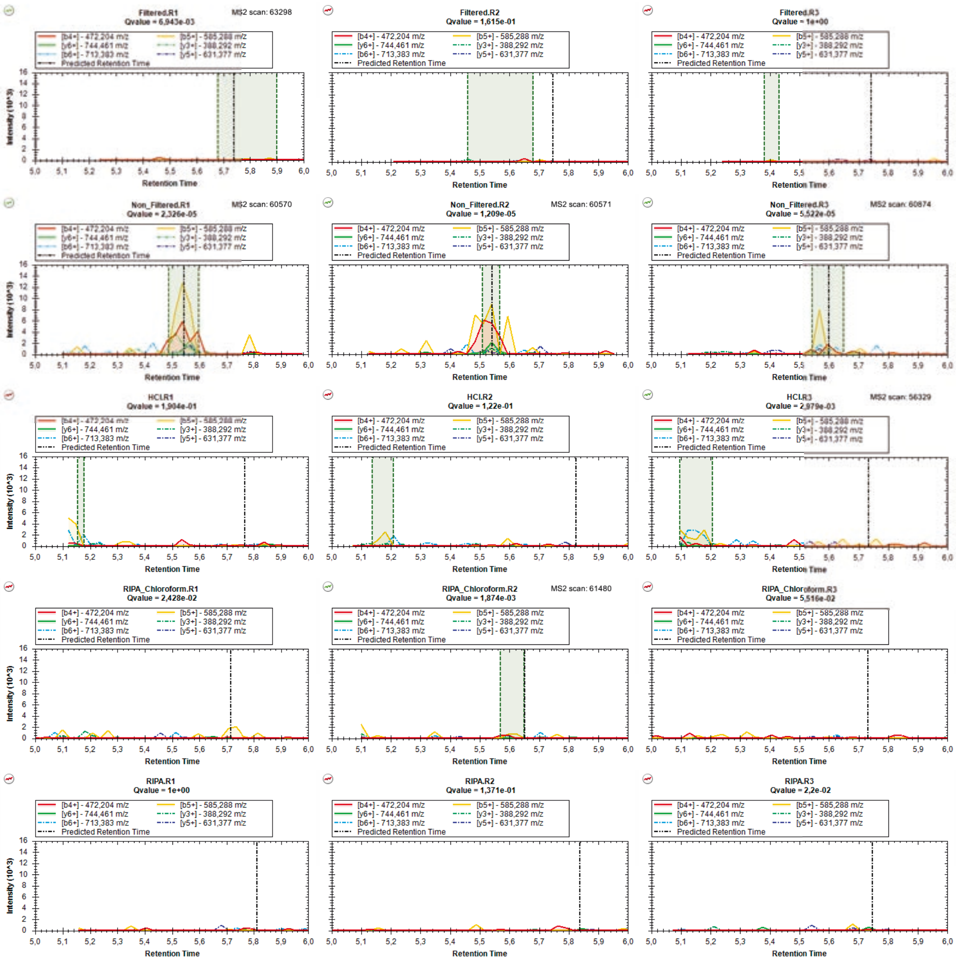

### mzspec_PXD066336_II_3767174_DVGTDEEEETAK

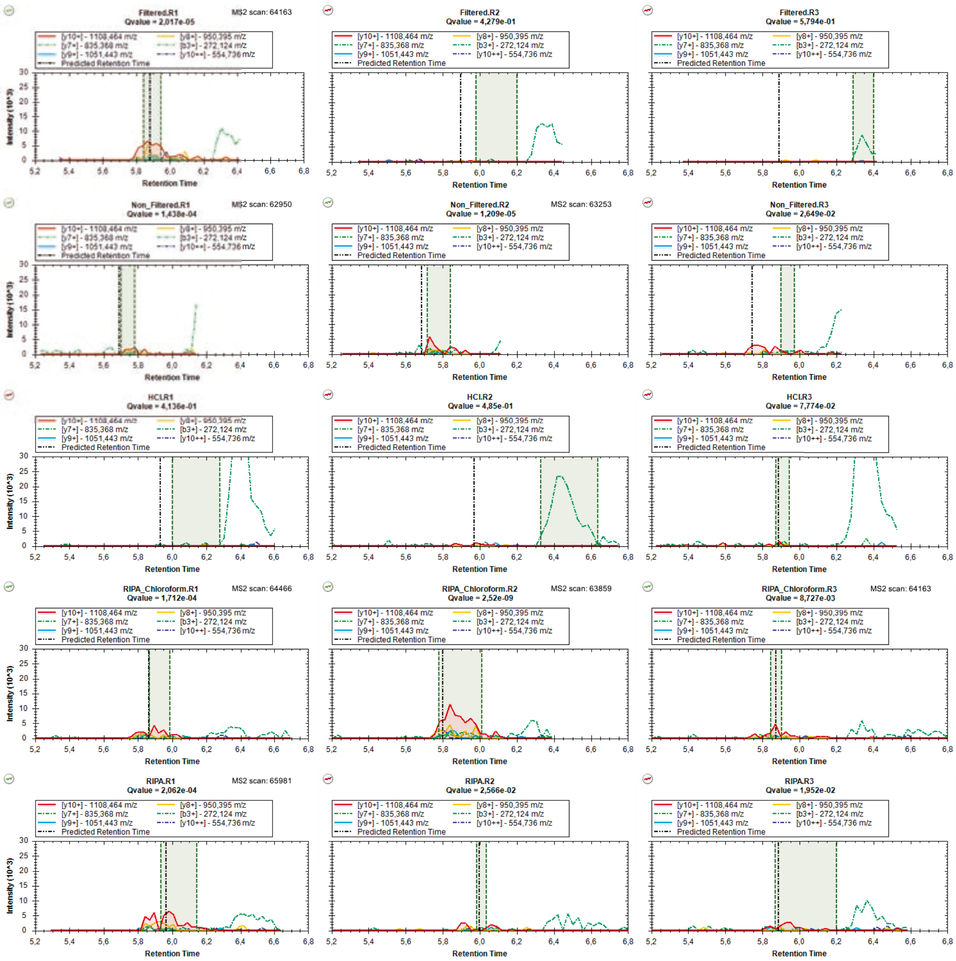

### mzspec_PXD066336_IP_579723_EEVPSGLAPR

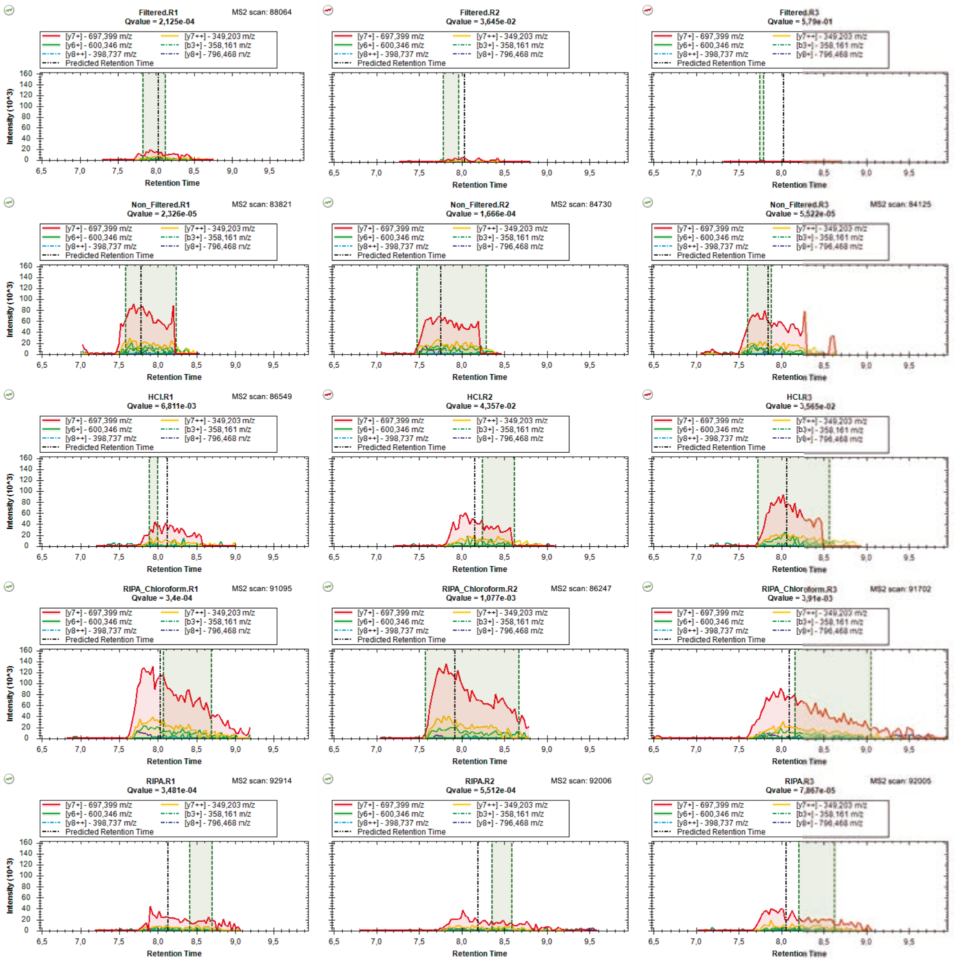

### mzspec_PXD066336_IP_3427170_FGFLFLK

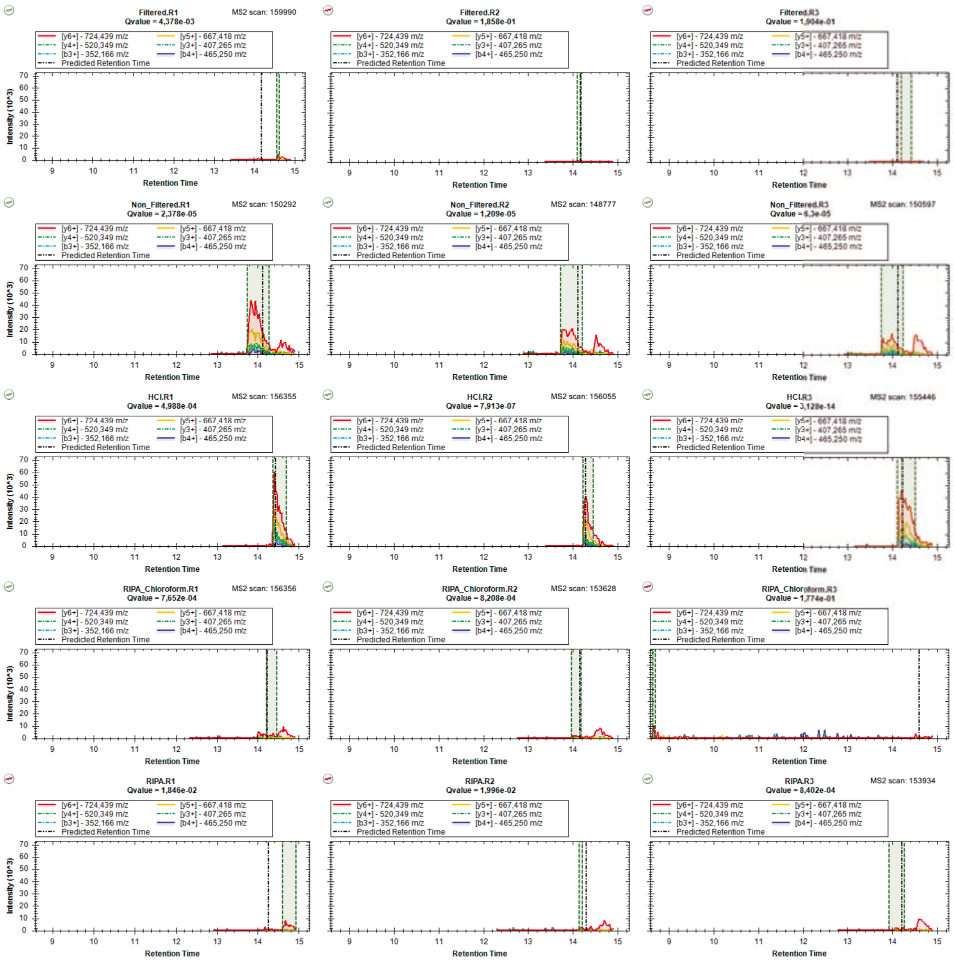

### mzspec_PXD066336_IP_3771255_FILNIYSK

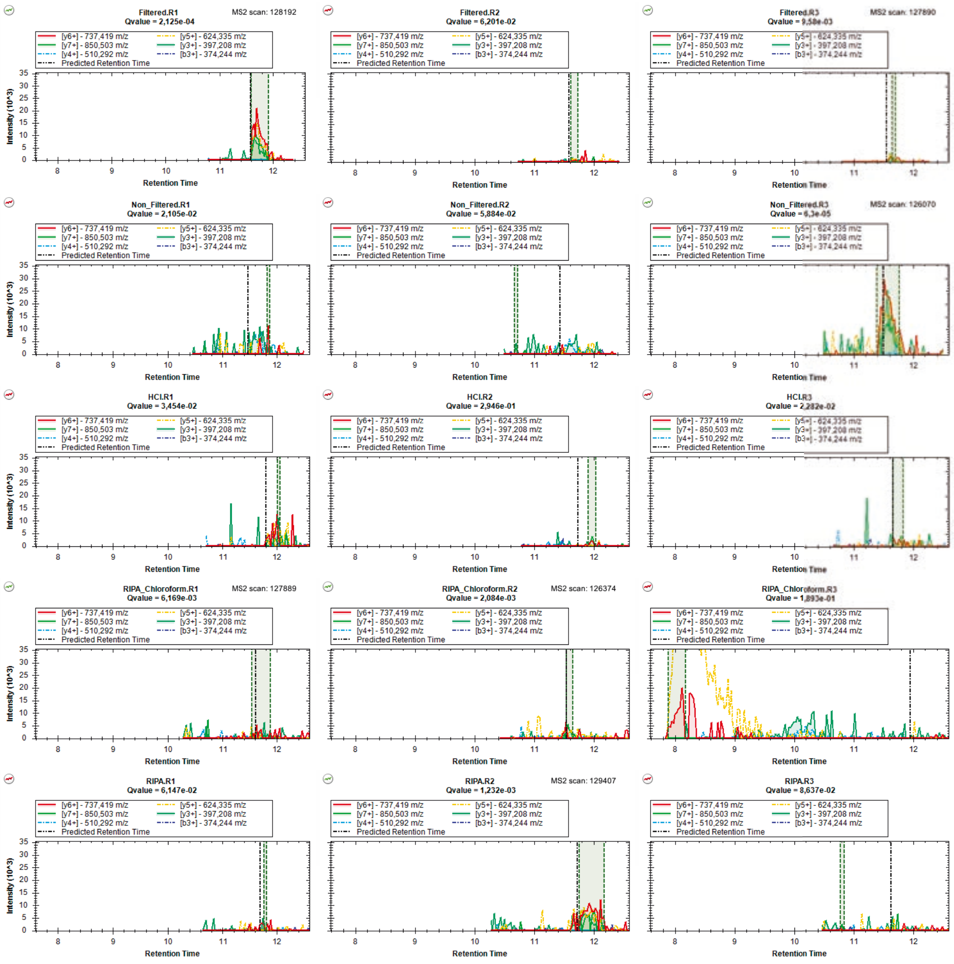

### mzspec_PXD066336_IP_2383646_FLPFHTVSEFK

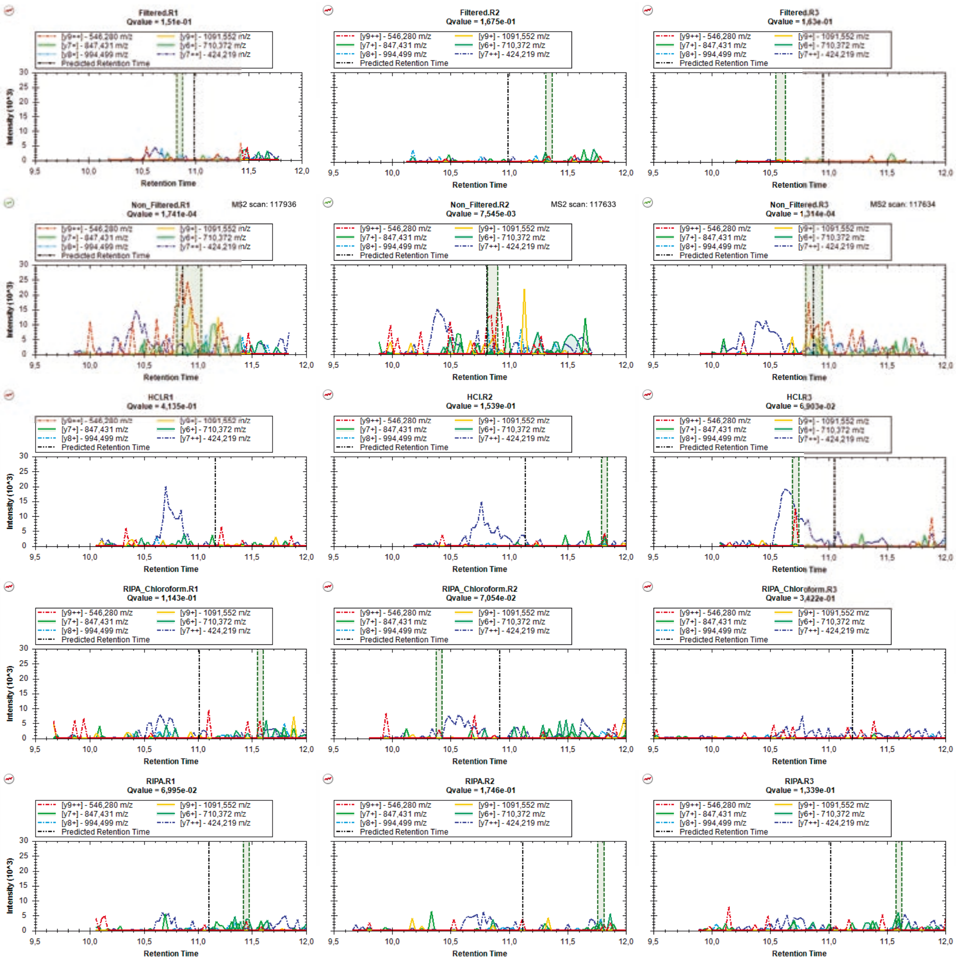

### mzspec_PXD066336_IP_779470_FVTVTSGESWK

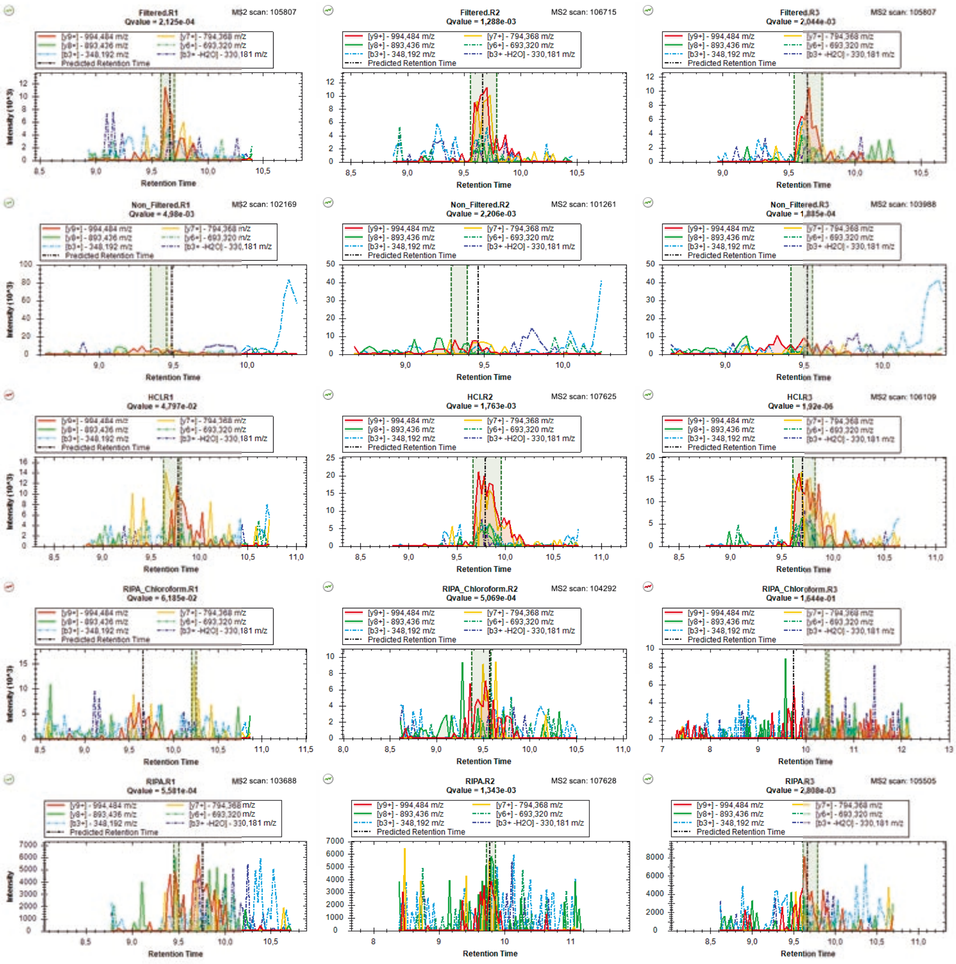

### mzspec_PXD066336_IP_772751_GFFVFLK_Charge_2

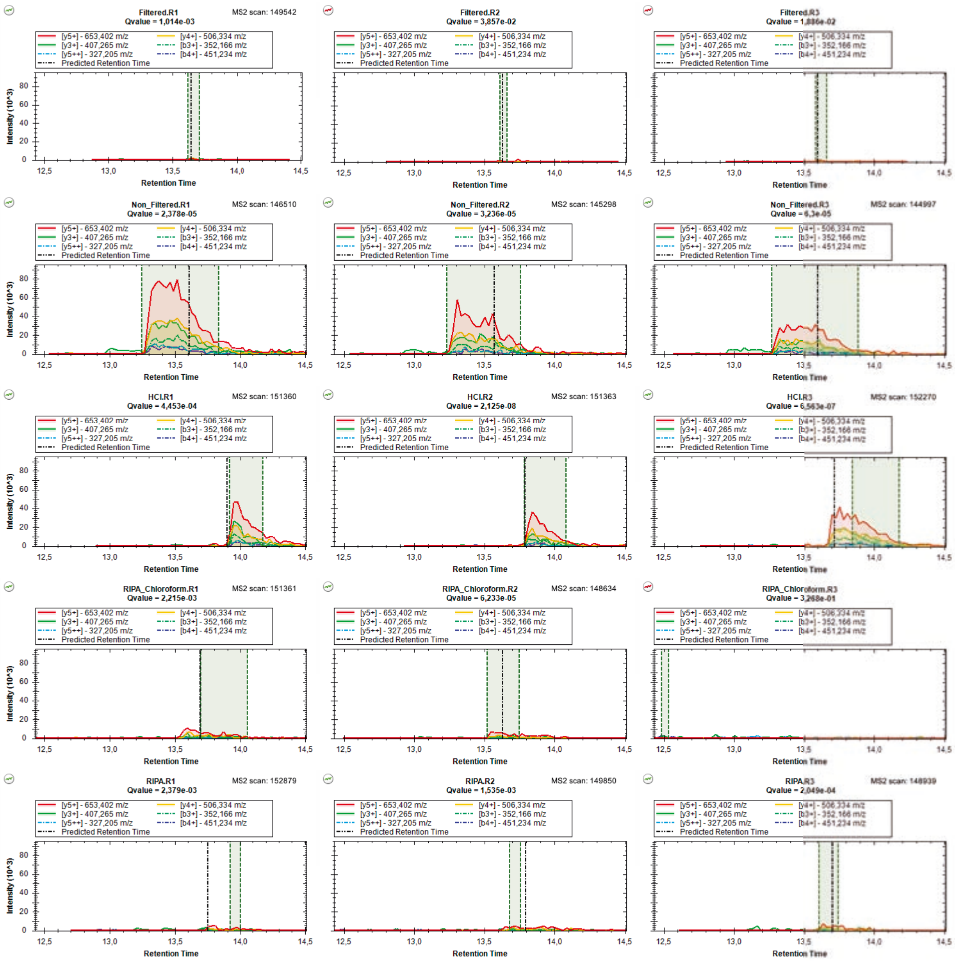

### mzspec_PXD066336_IP_772751_GFFVFLK_Charge_1

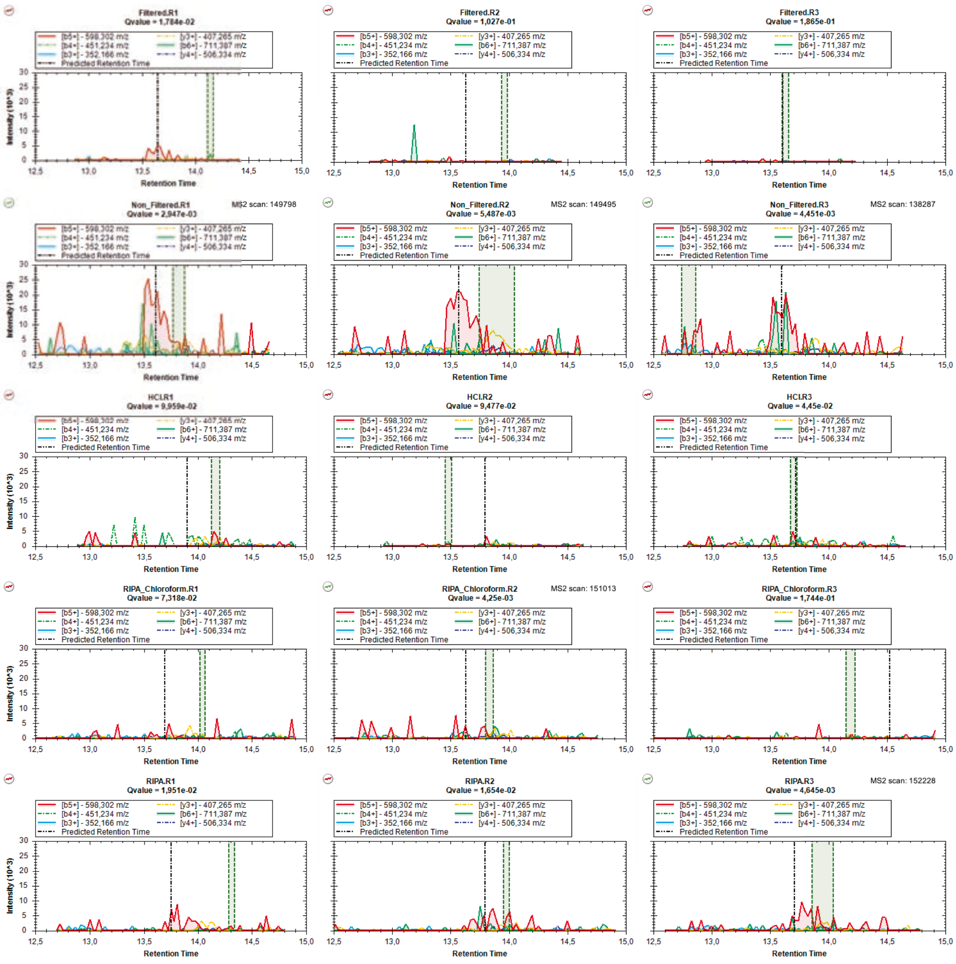

### mzspec_PXD066336_IP_594653_GIGILSLK_Charge_1

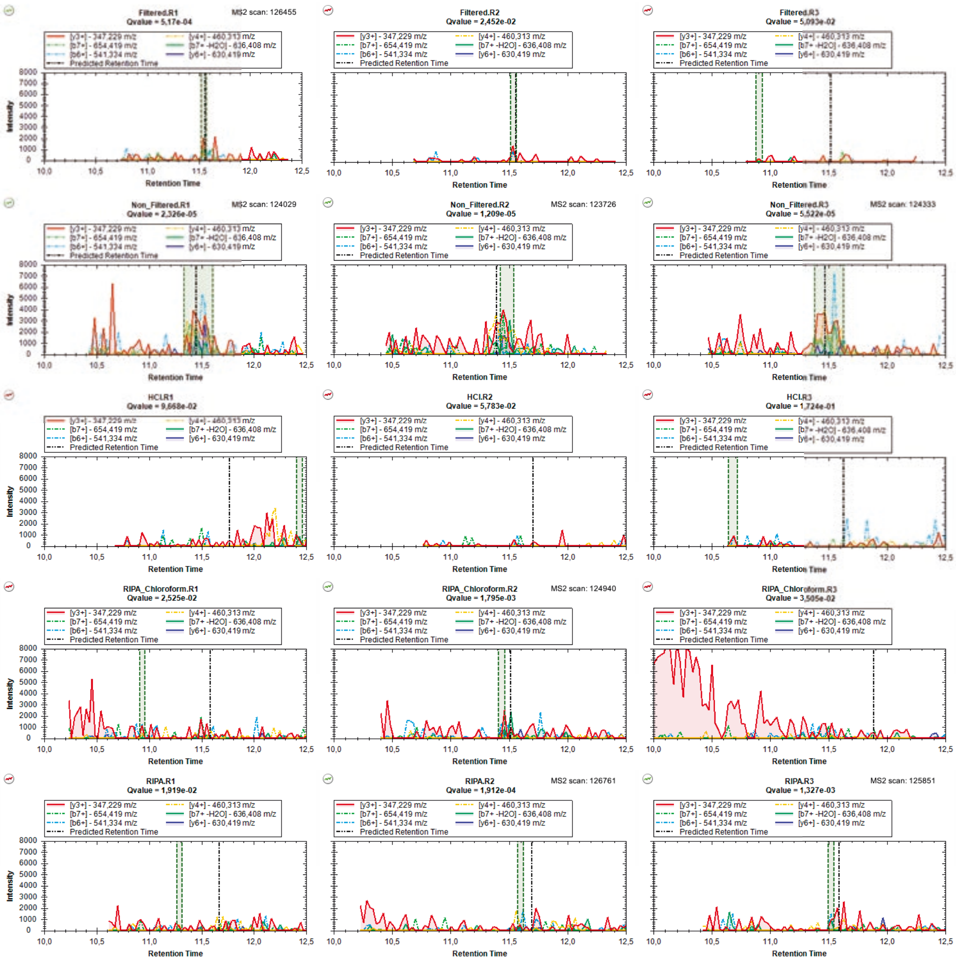

### mzspec_PXD066336_IP_594653_GIGILSLK_Charge_2

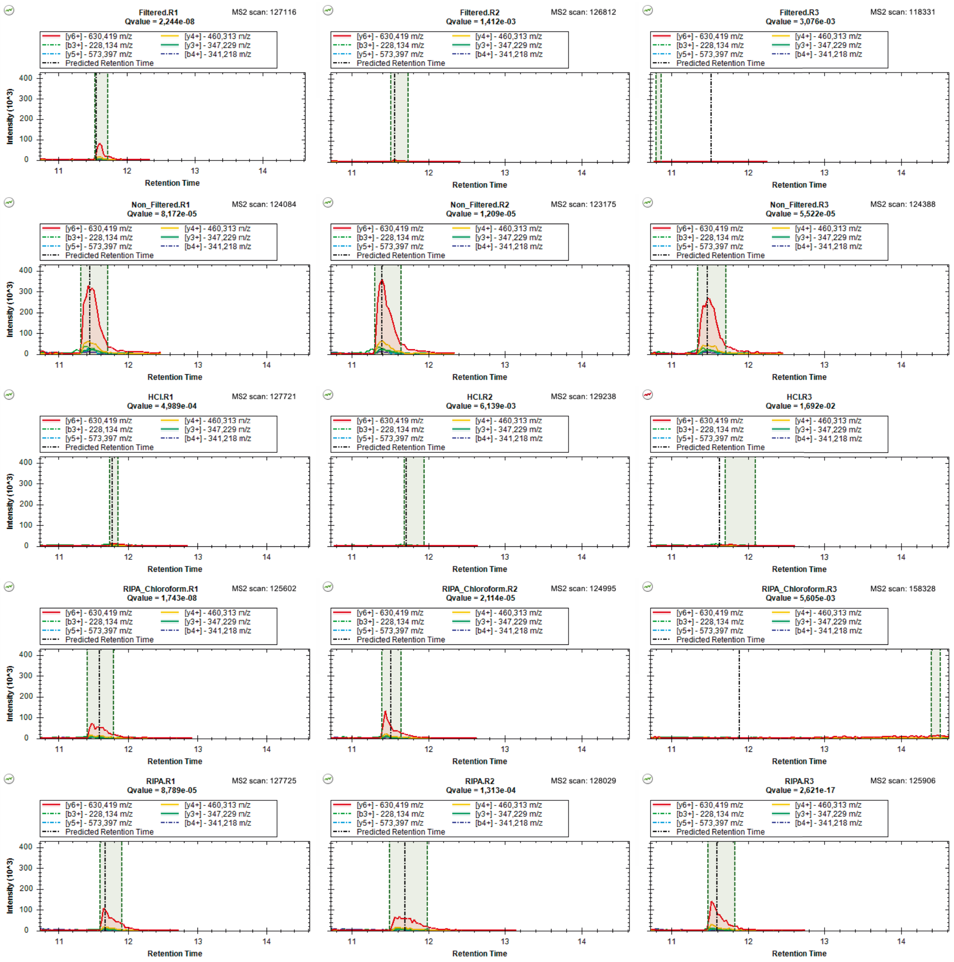

### mzspec_PXD066336_II_719305_GPGLLPLLAGEGR

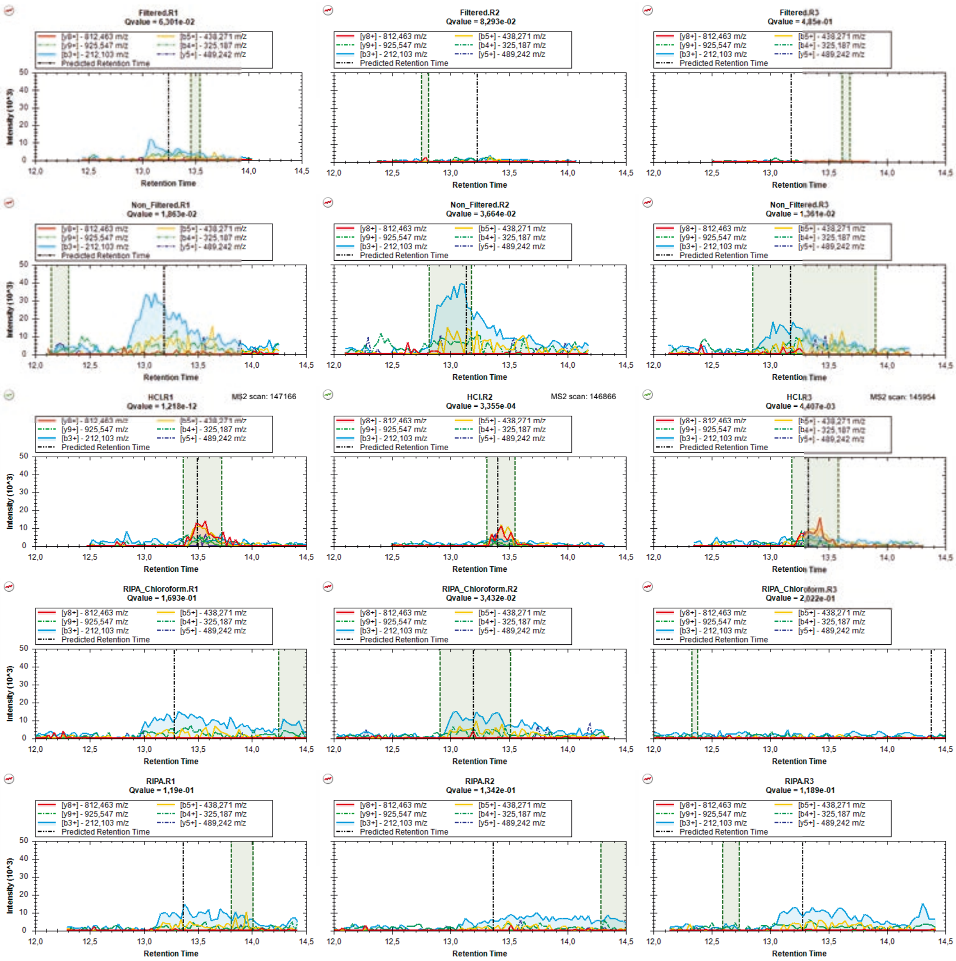

### mzspec_PXD066336_IP_668161_GPPDPILGVTEAFK

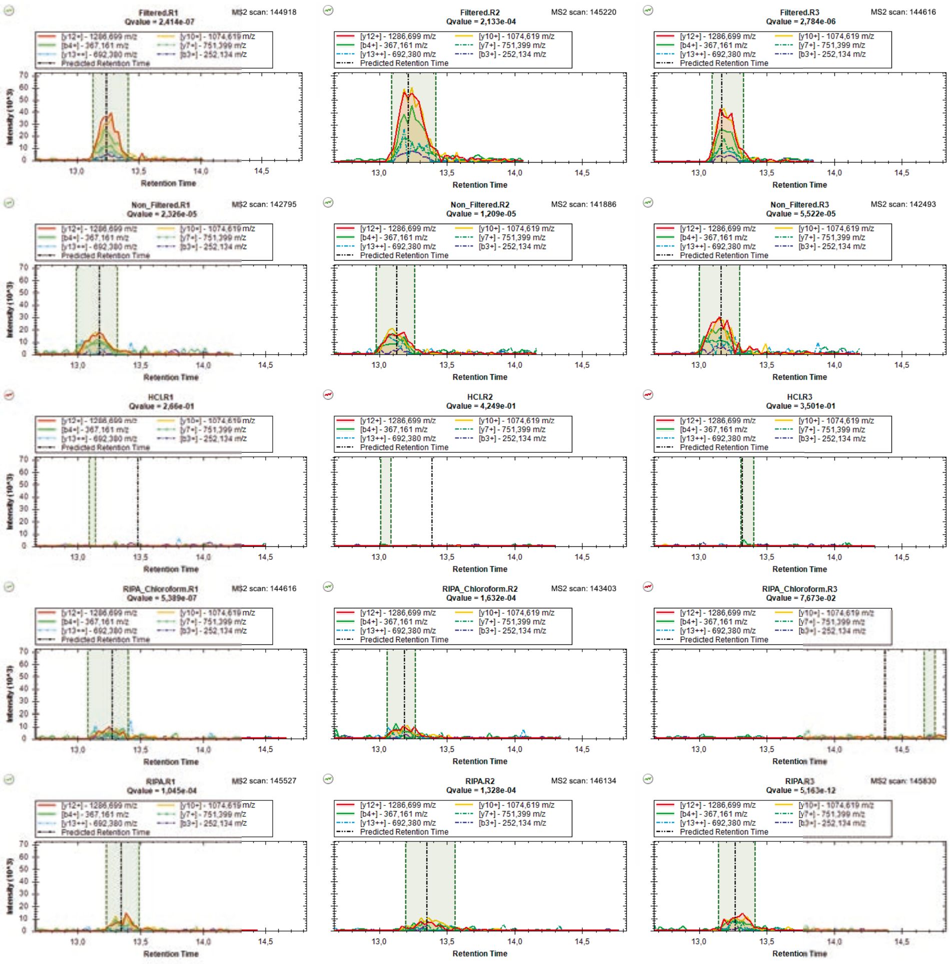

### mzspec_PXD066336_IP_668161_GPPDPILGVTEAFKR

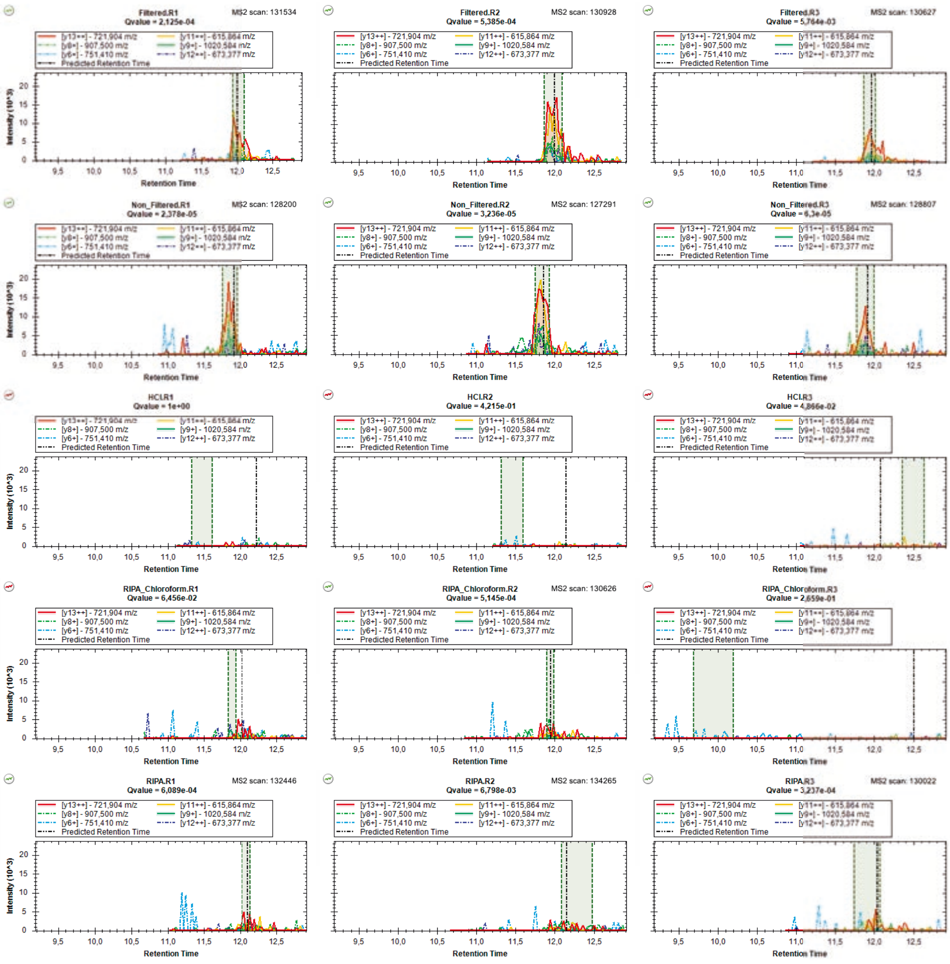

### mzspec_PXD066336_IP_4488681_GPTVLGQGEEGAAGSGR

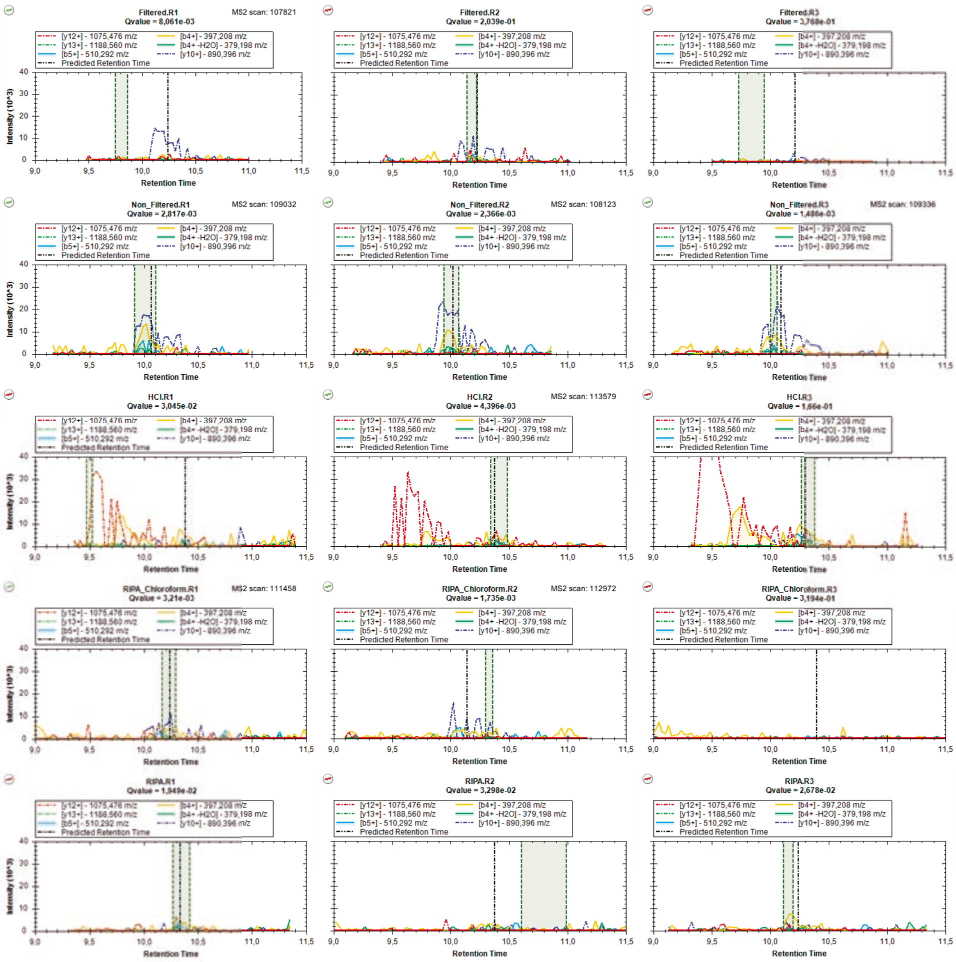

### mzspec_PXD066336_IP_3800027_GTLANFR

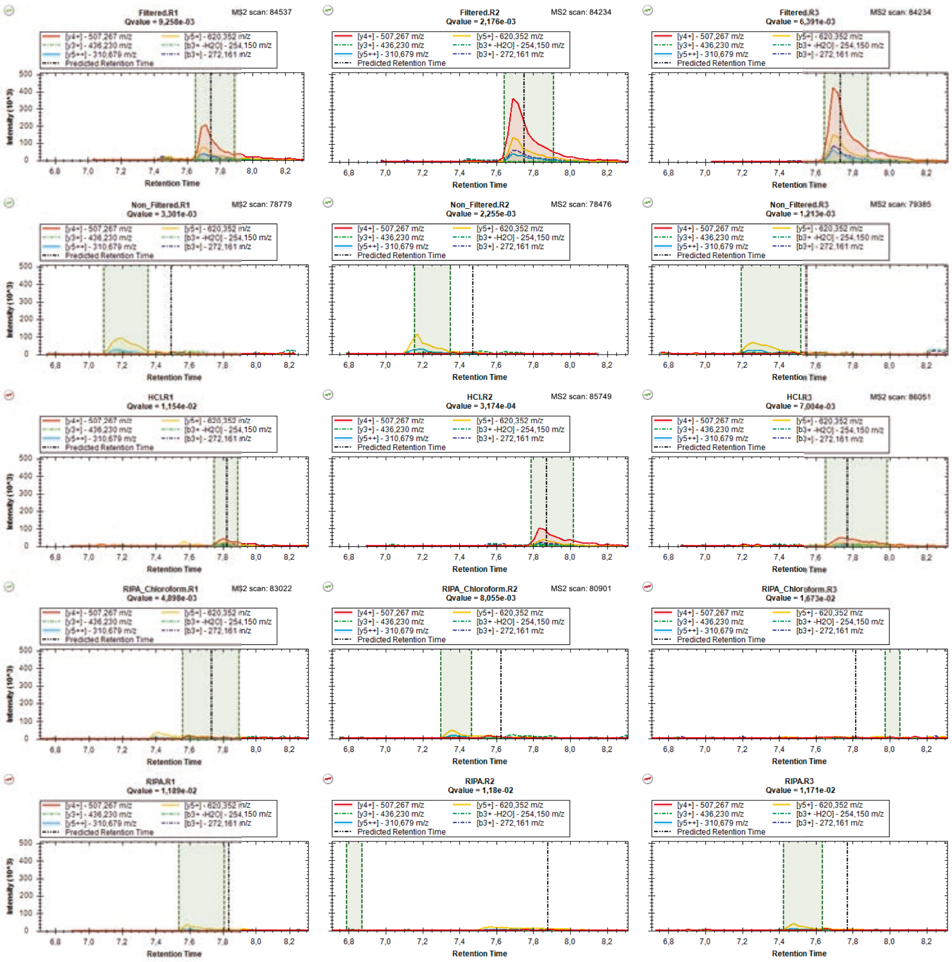

### mzspec_PXD066336_IP_557277_IEIPNHFMMRPDK_Charge_2

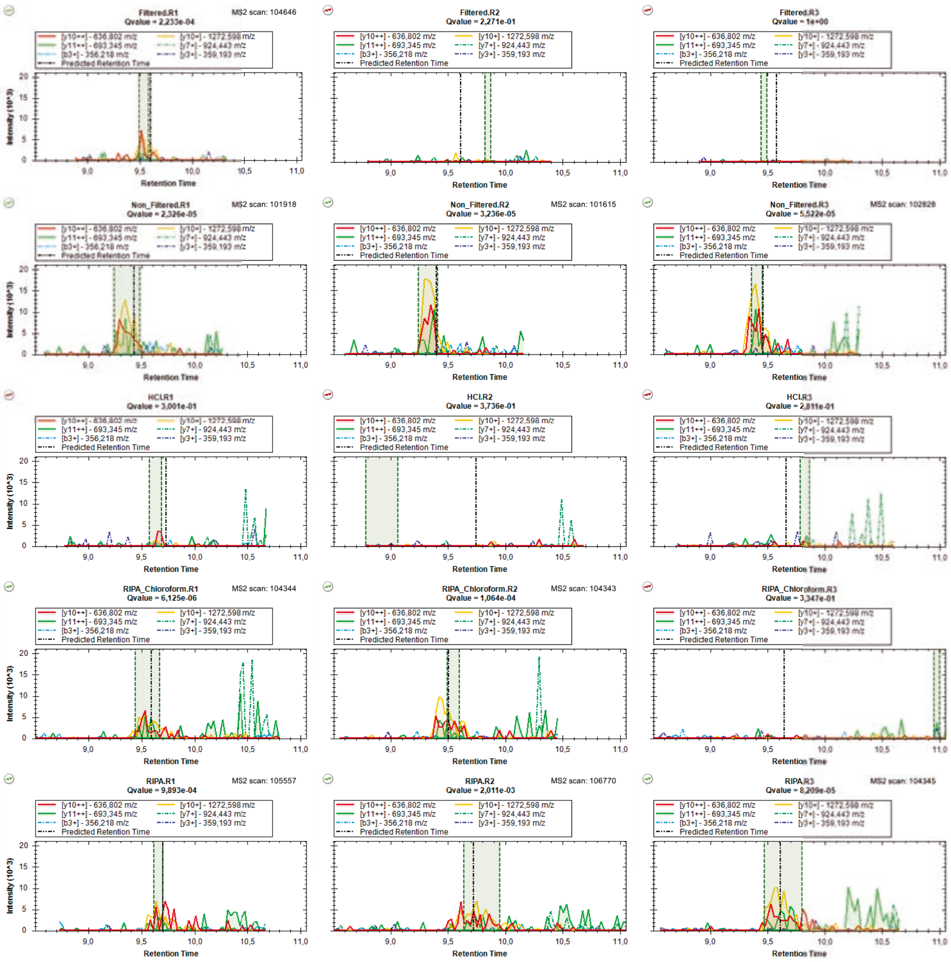

### mzspec_PXD066336_IP_557277_IEIPNHFMMRPDK_Charge_3

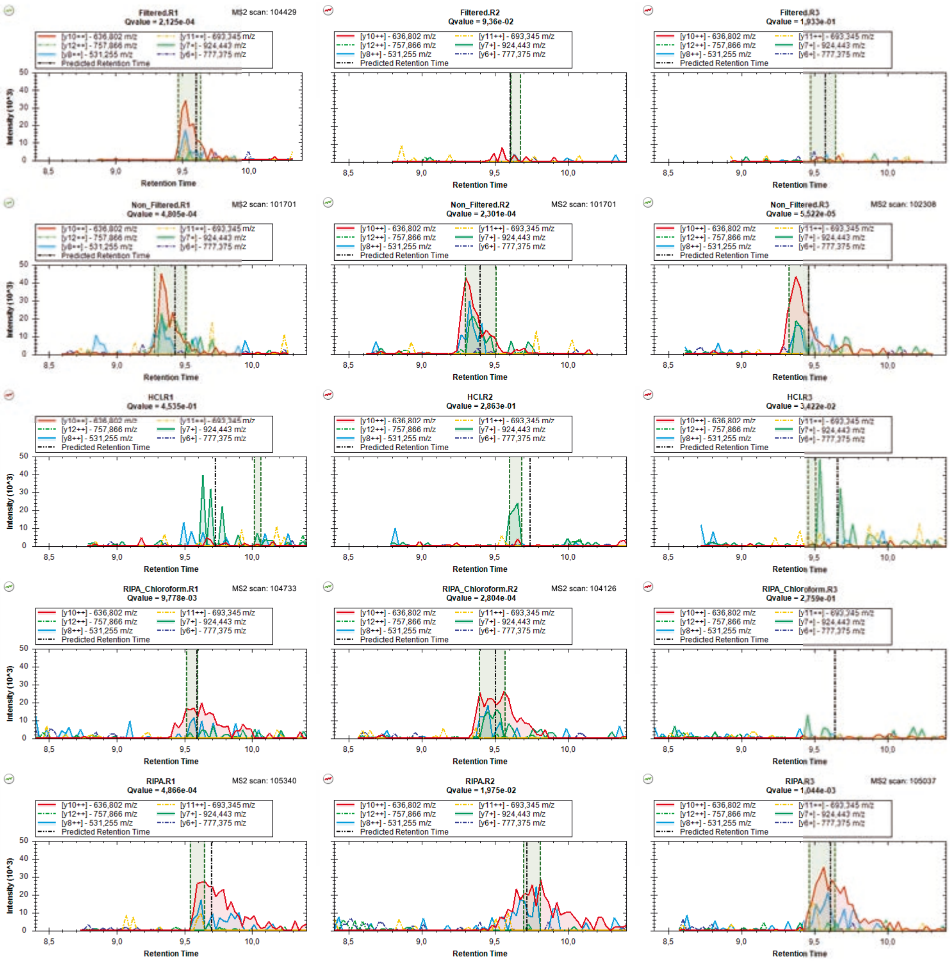

### mzspec_PXD066336_A0A1B0GW03_IFFINDEEK

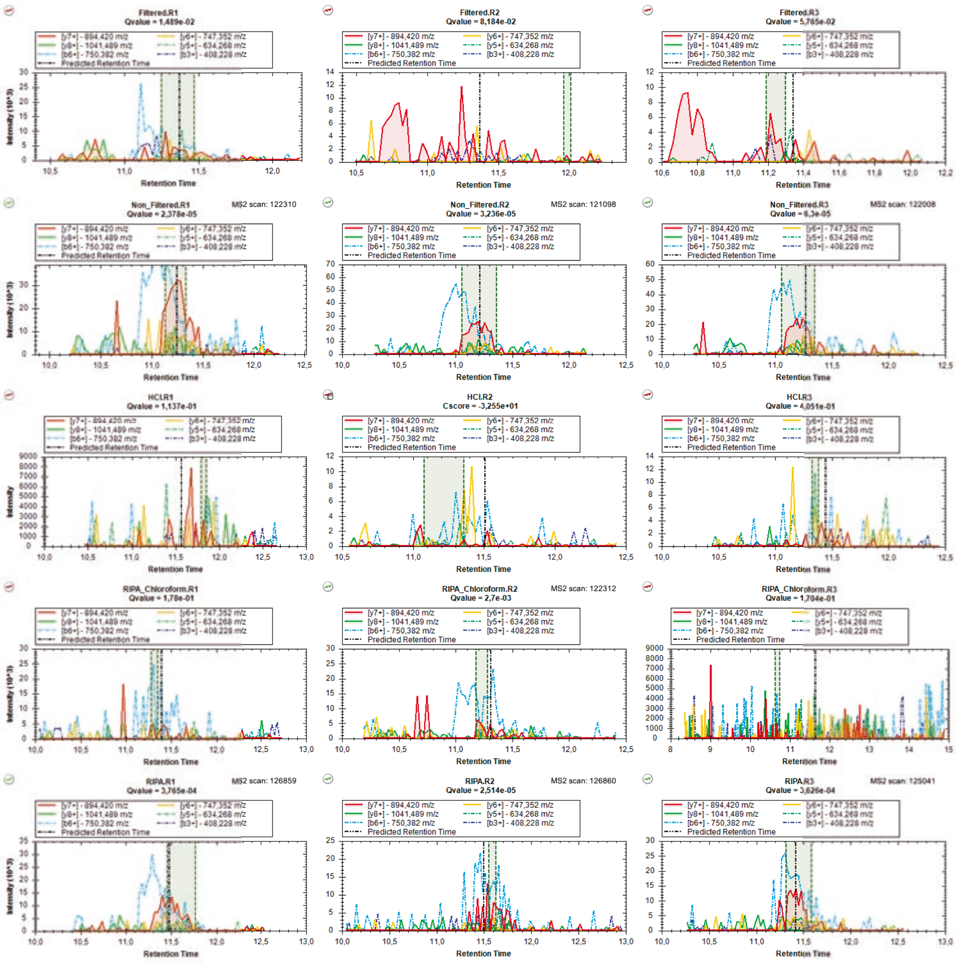

### mzspec_PXD066336_IP_725389_IFLLQACR

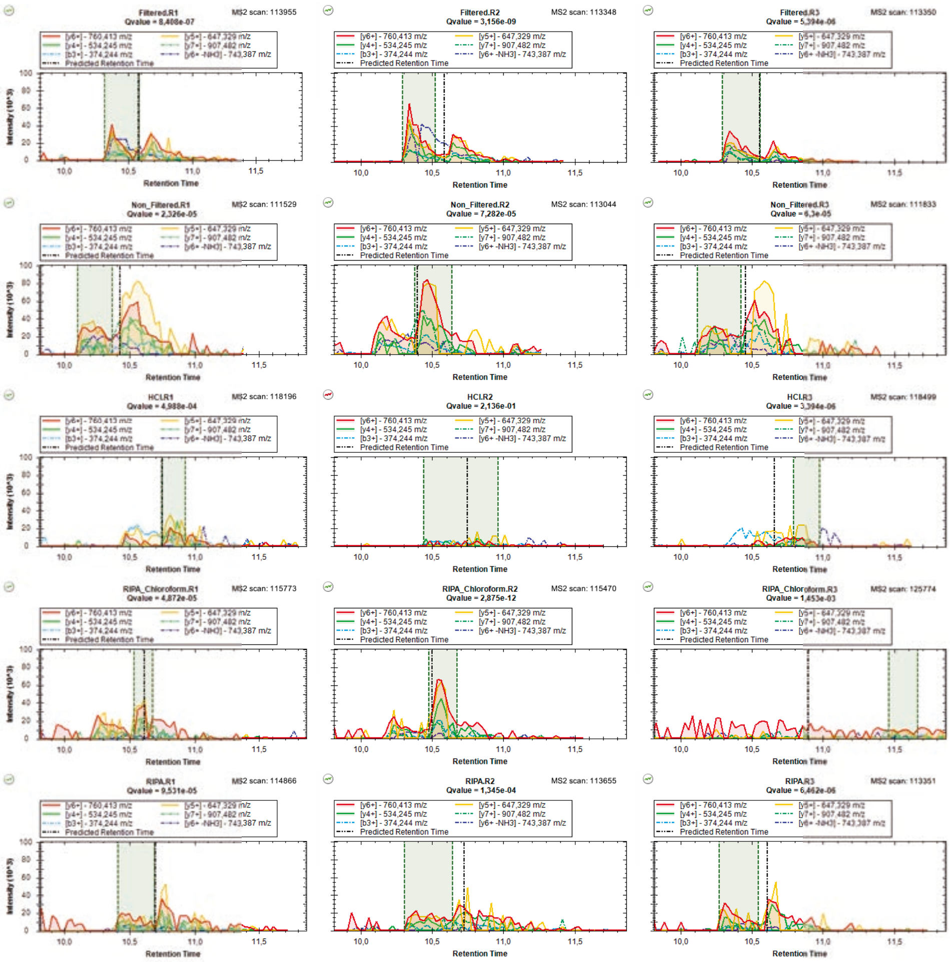

### mzspec_PXD066336_IP_602537_LCYVTLDFEQEMATVASSSSLEK

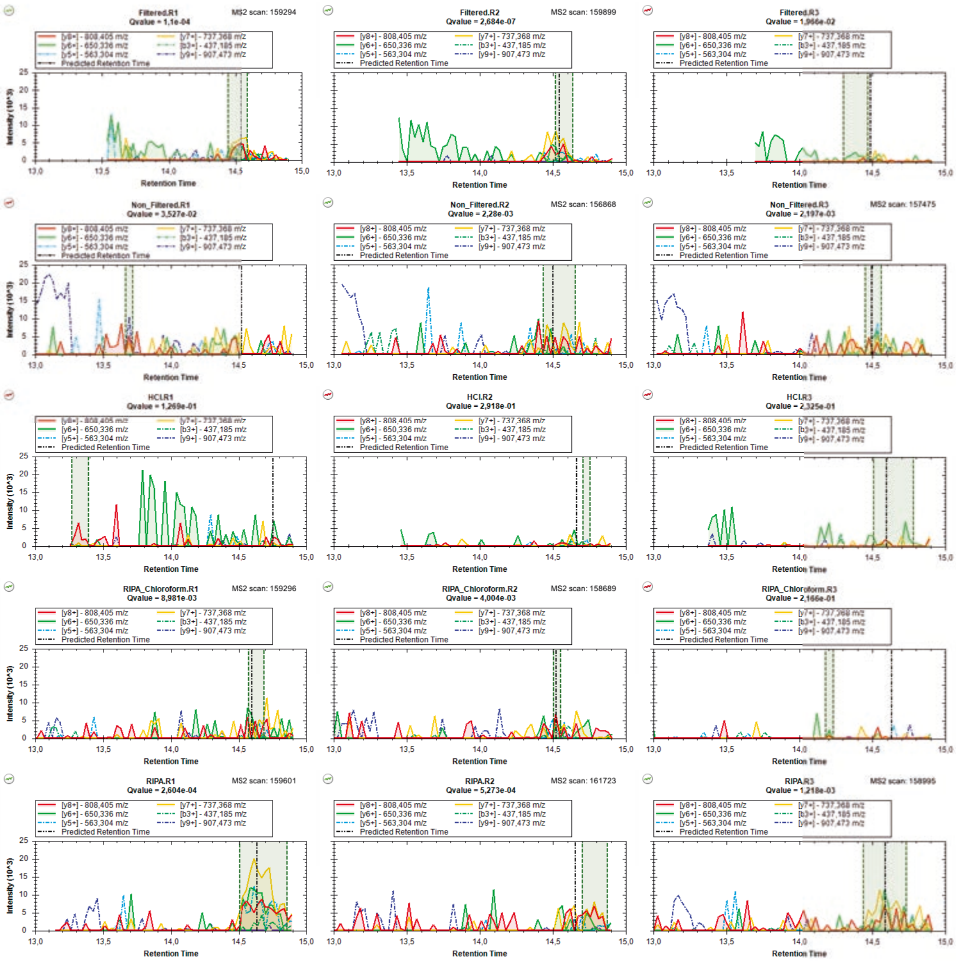

### mzspec_PXD066336_IP_620432_LDLSMSPK

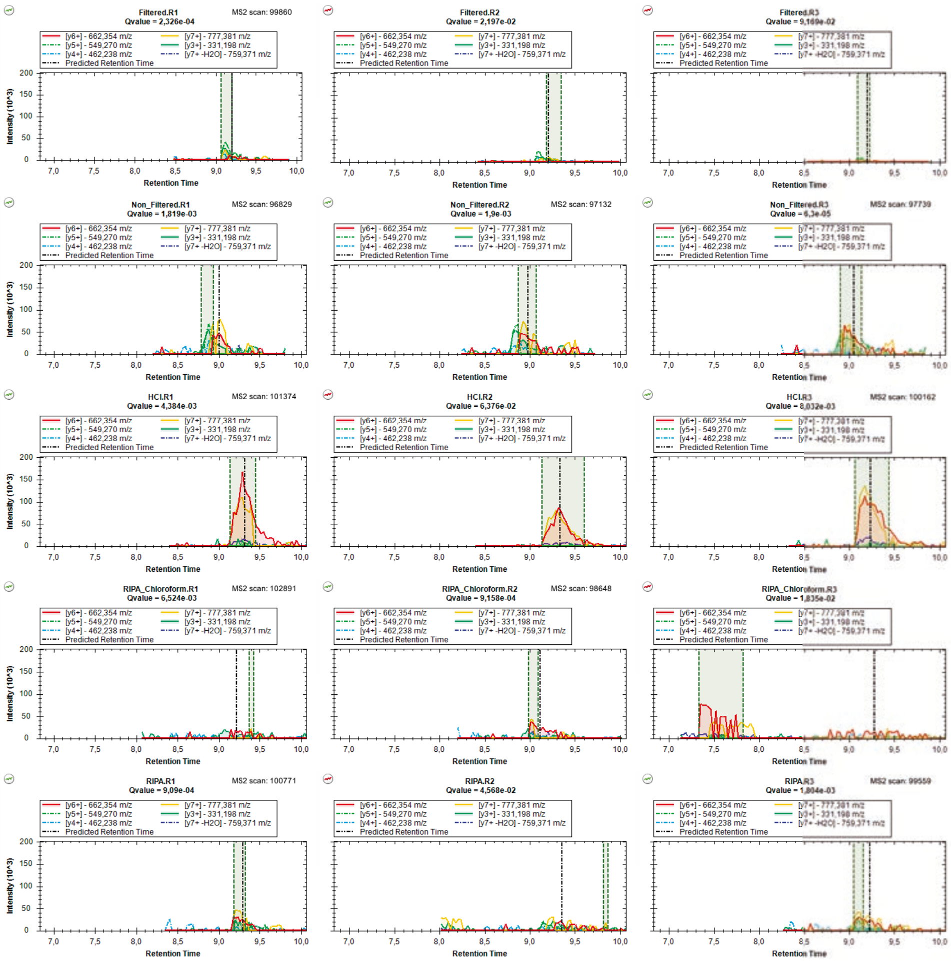

### mzspec_PXD066336_IP_3801301_LLLHLTSK

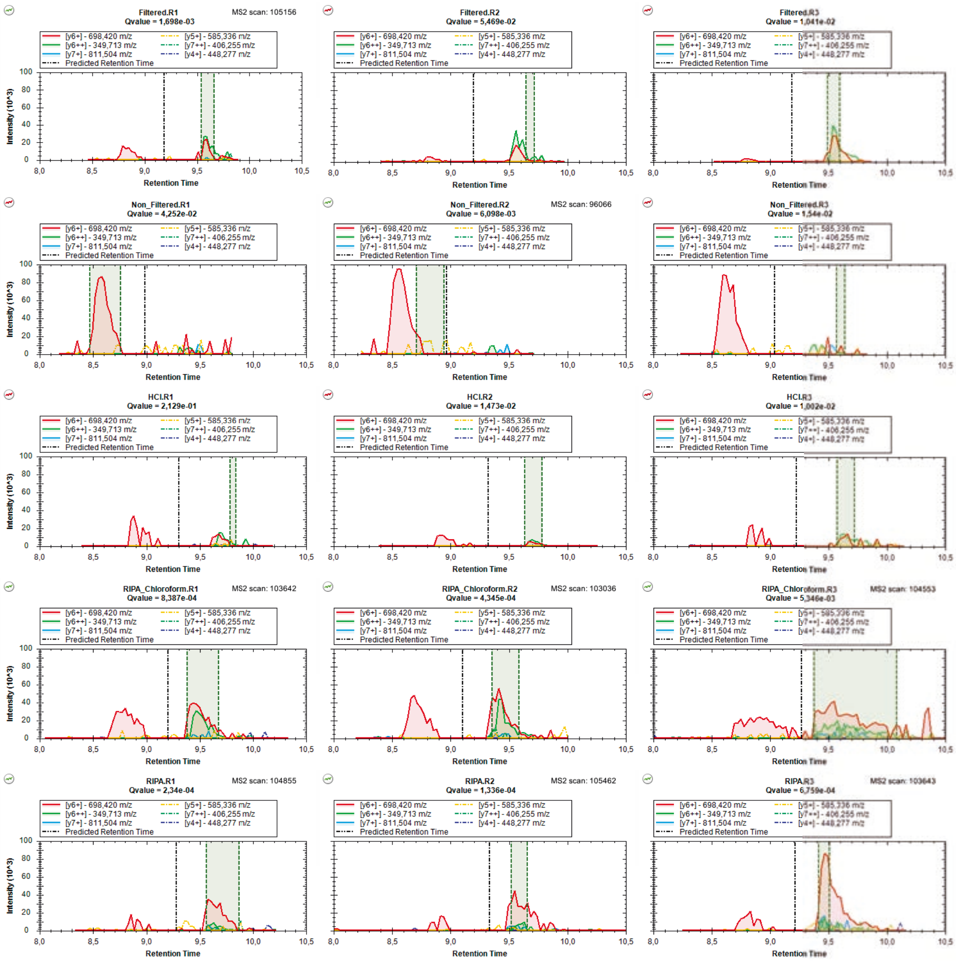

### mzspec_PXD066336_IP_782672_LPCGPLGSK

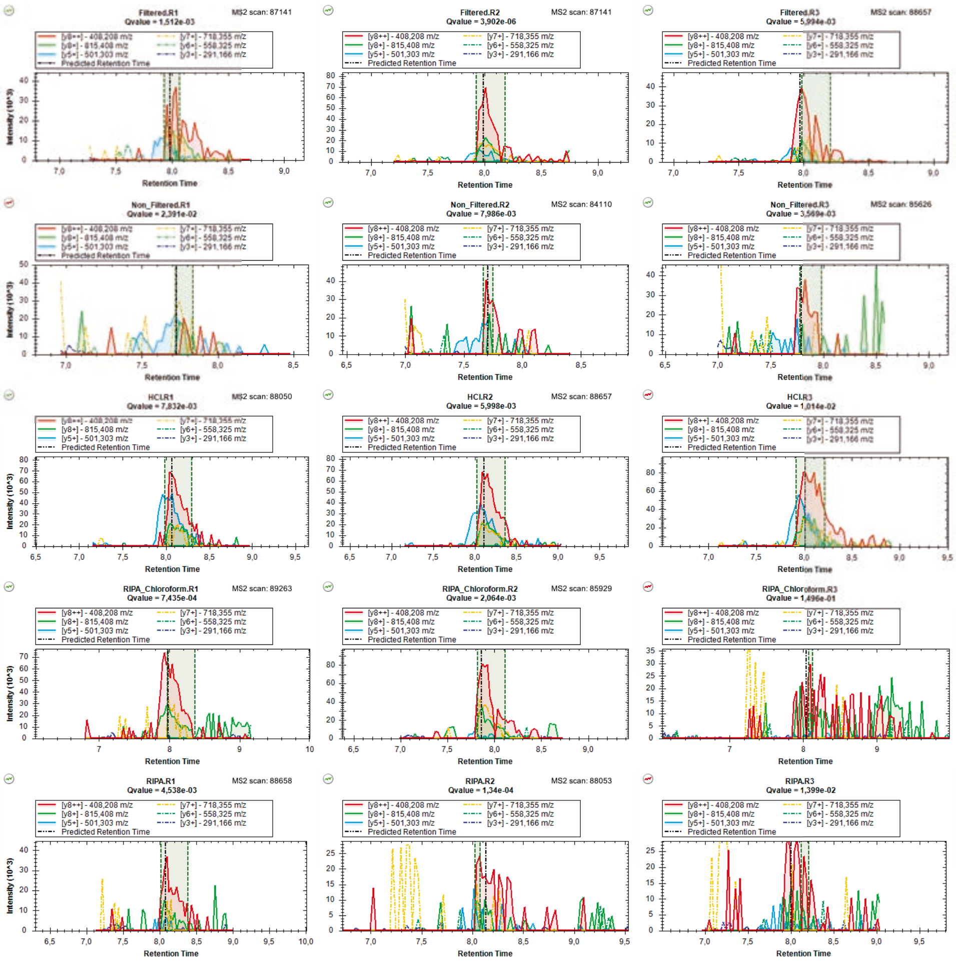

### mzspec_PXD066336_IP_580544_LPGNYLLEK

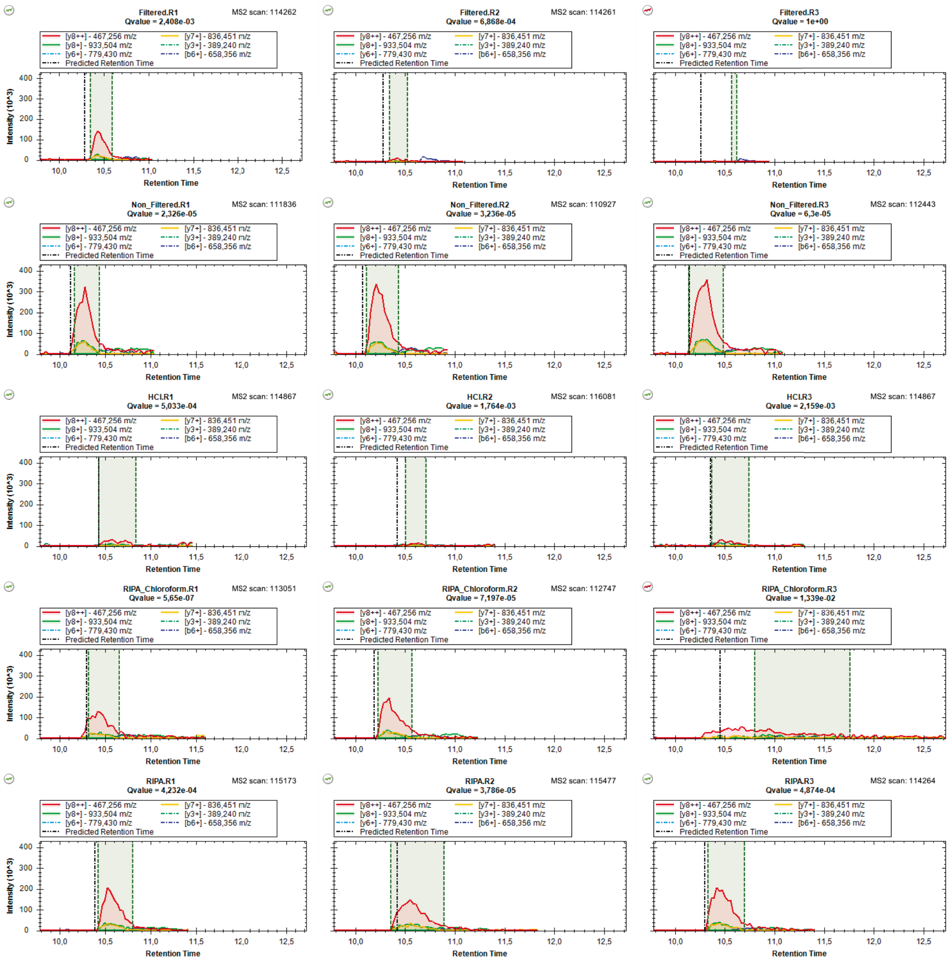

### mzspec_PXD066336_IP_674463_MALDALADGGVK

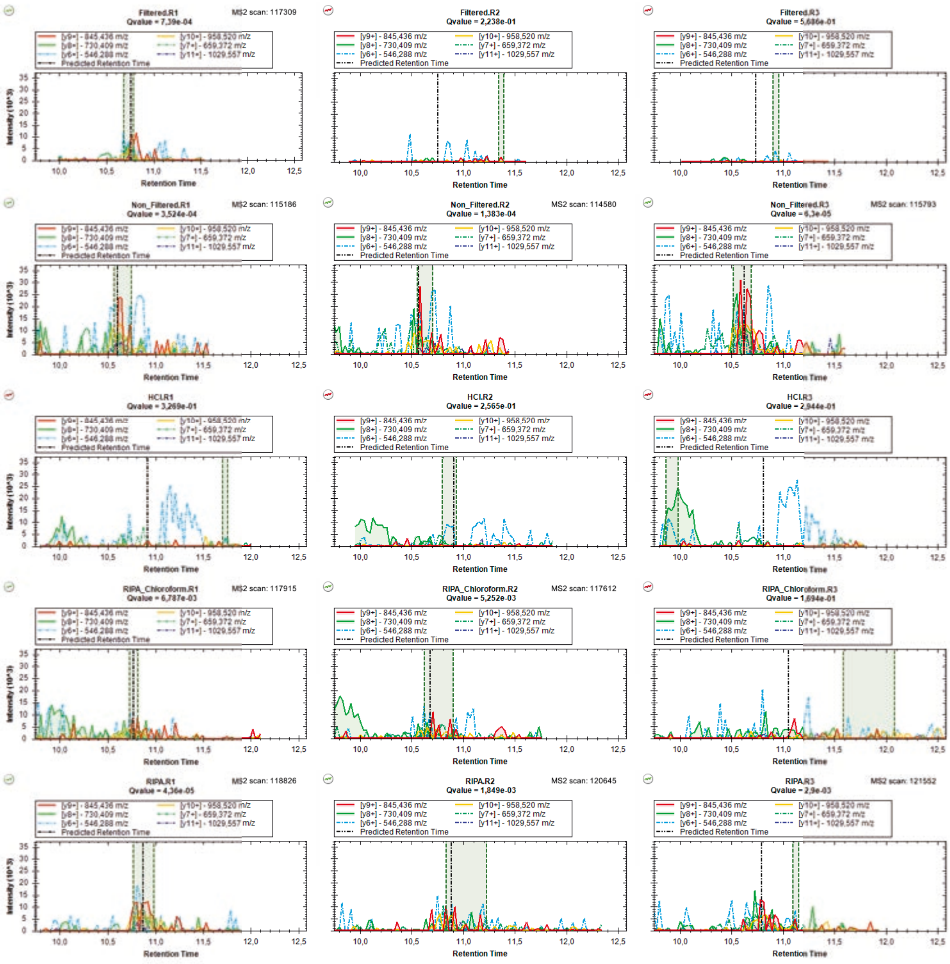

### mzspec_PXD066336_IP_566433_MAVNYVQHNR

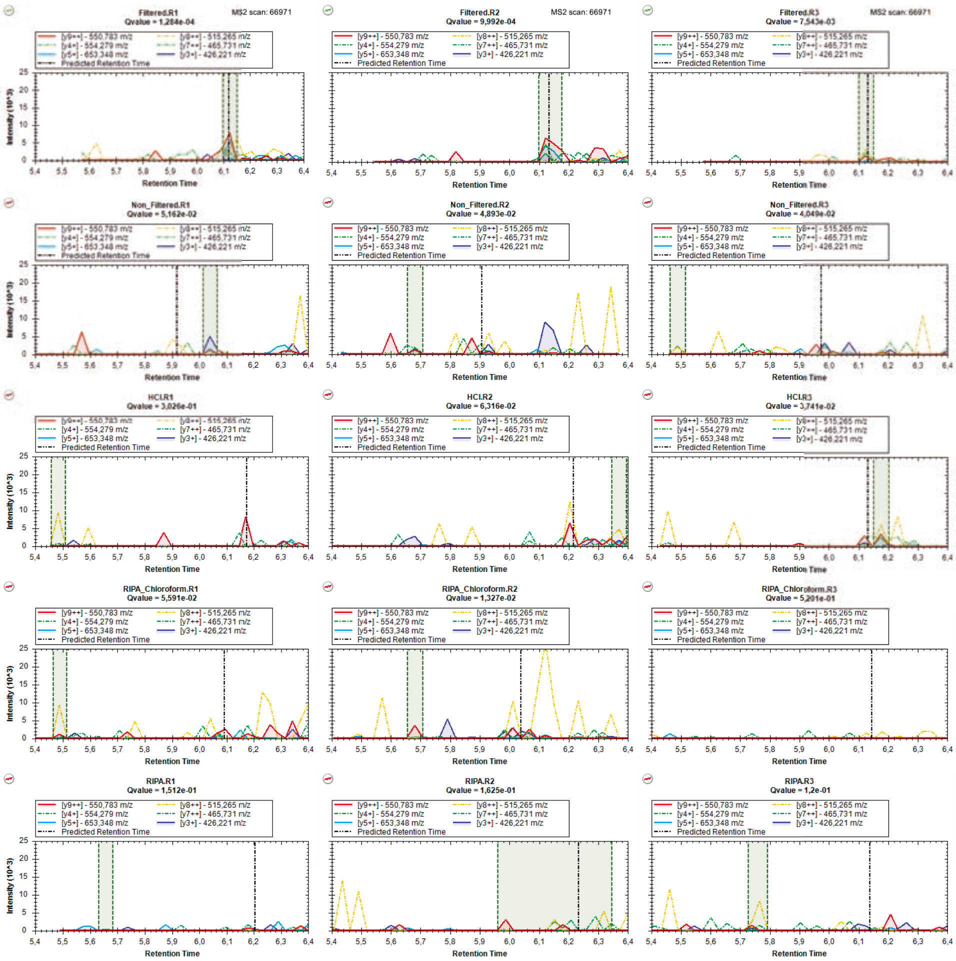

### mzspec_PXD066336_II_658785_MELKPNAGSDR_Charge_2

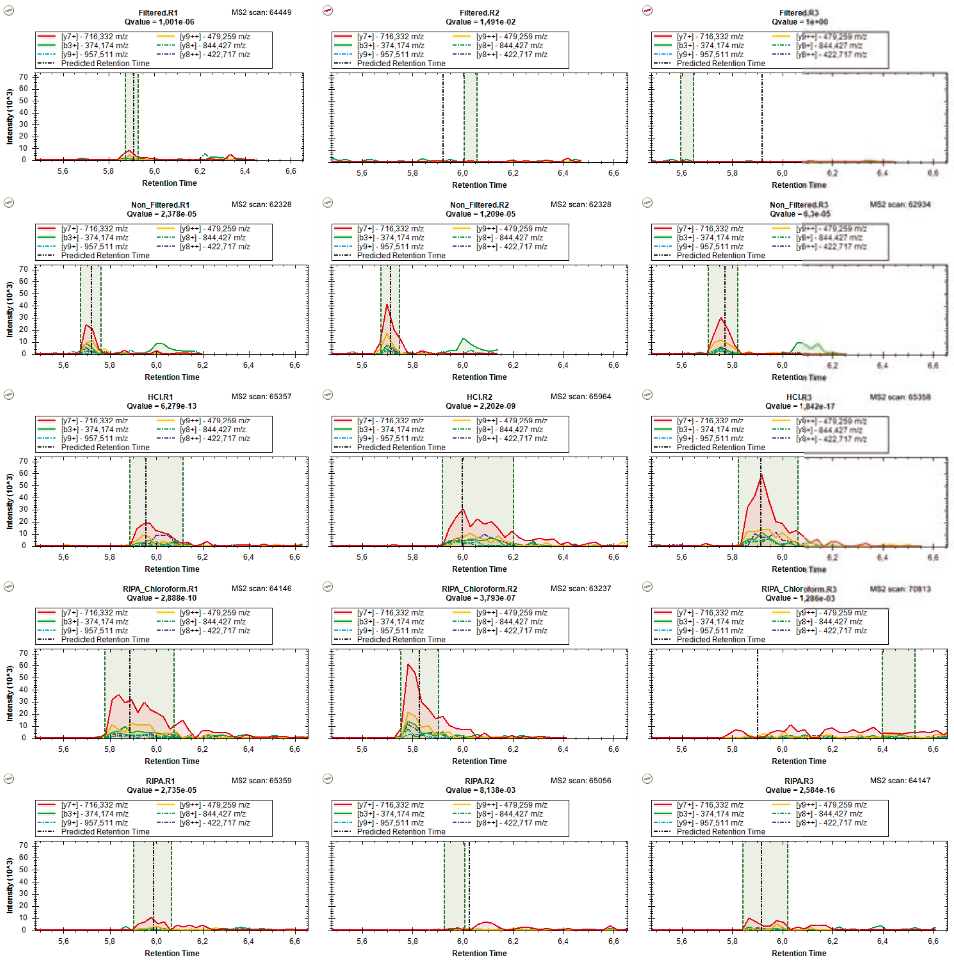

### mzspec_PXD066336_II_658785_MELKPNAGSDR_Charge_3

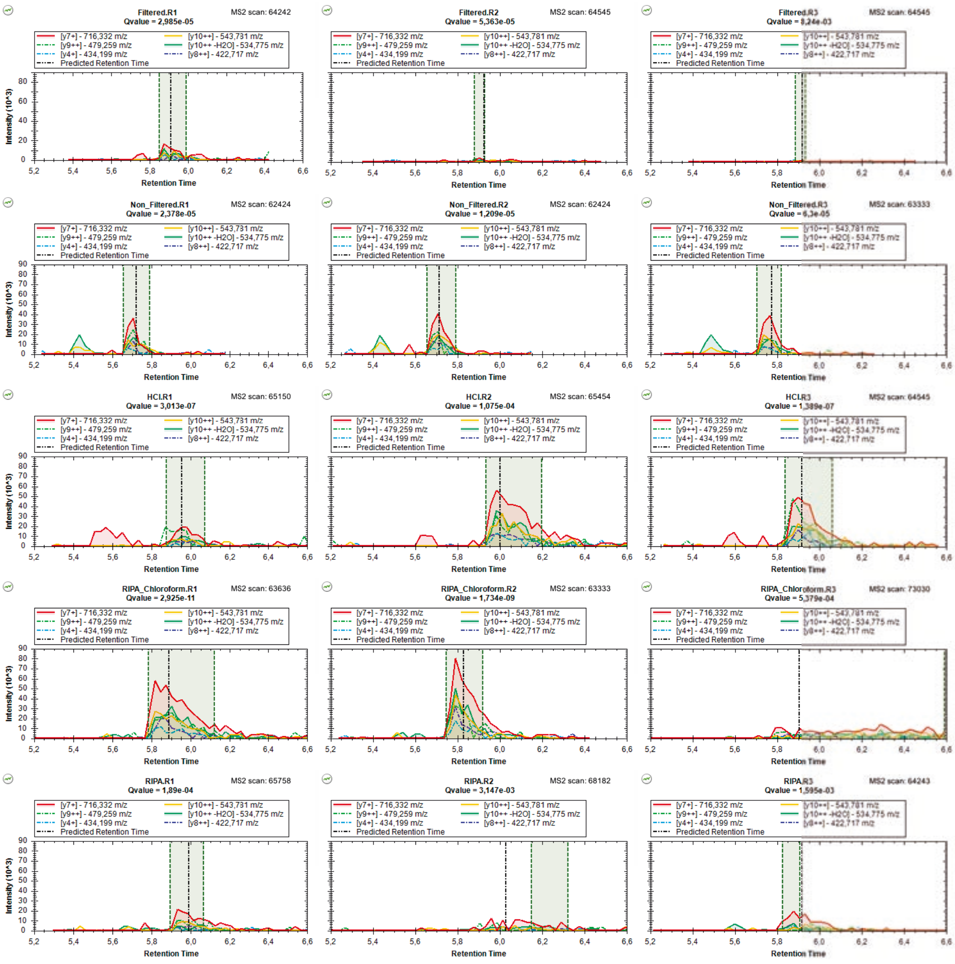

### mzspec_PXD066336_IP_741351_MLESMIK_Charge_1

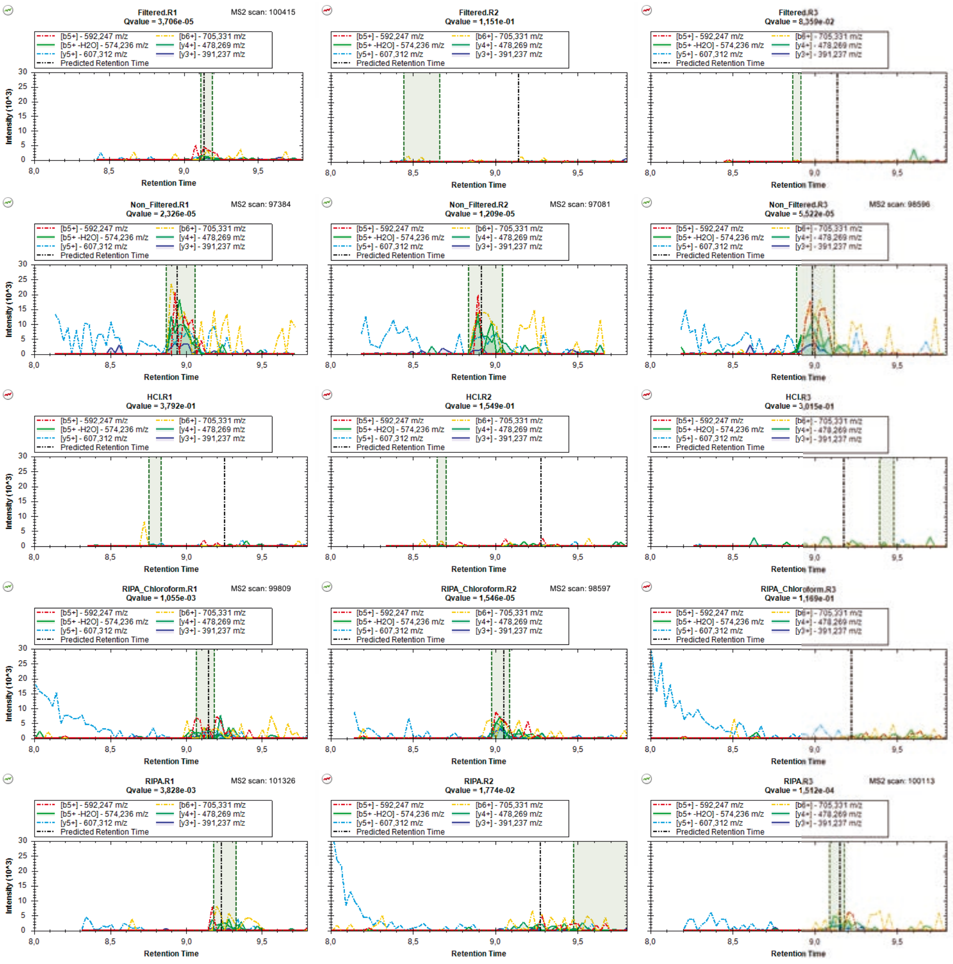

### mzspec_PXD066336_IP_741351_MLESMIK_Charge_2

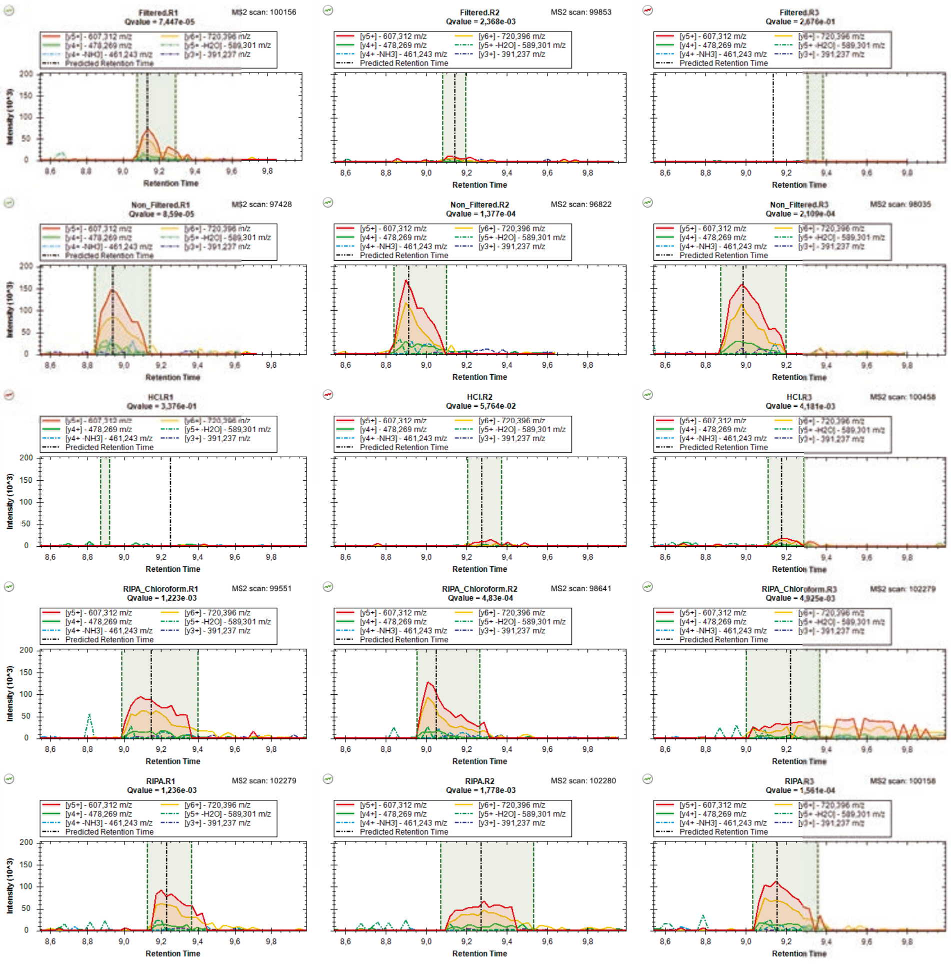

### mzspec_PXD066336_IP_647370_MLSNFHLEK

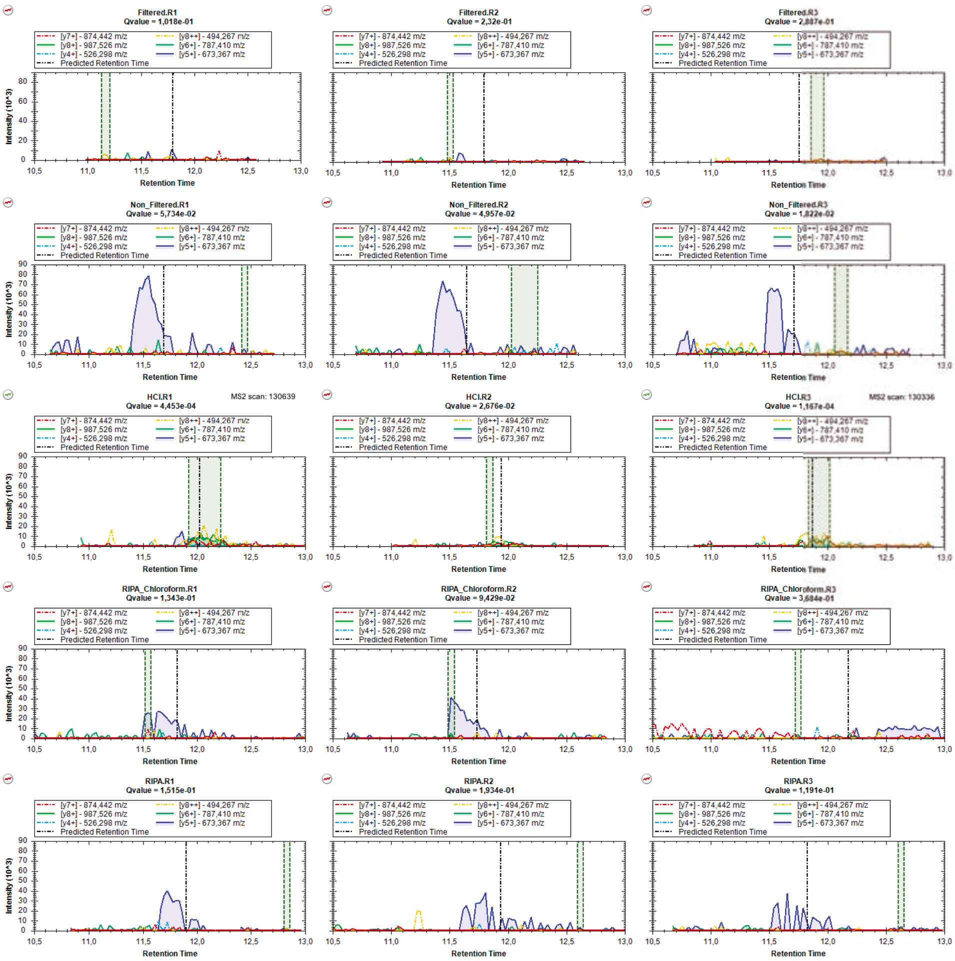

### mzspec_PXD066336_IP_577150_MVLLLER

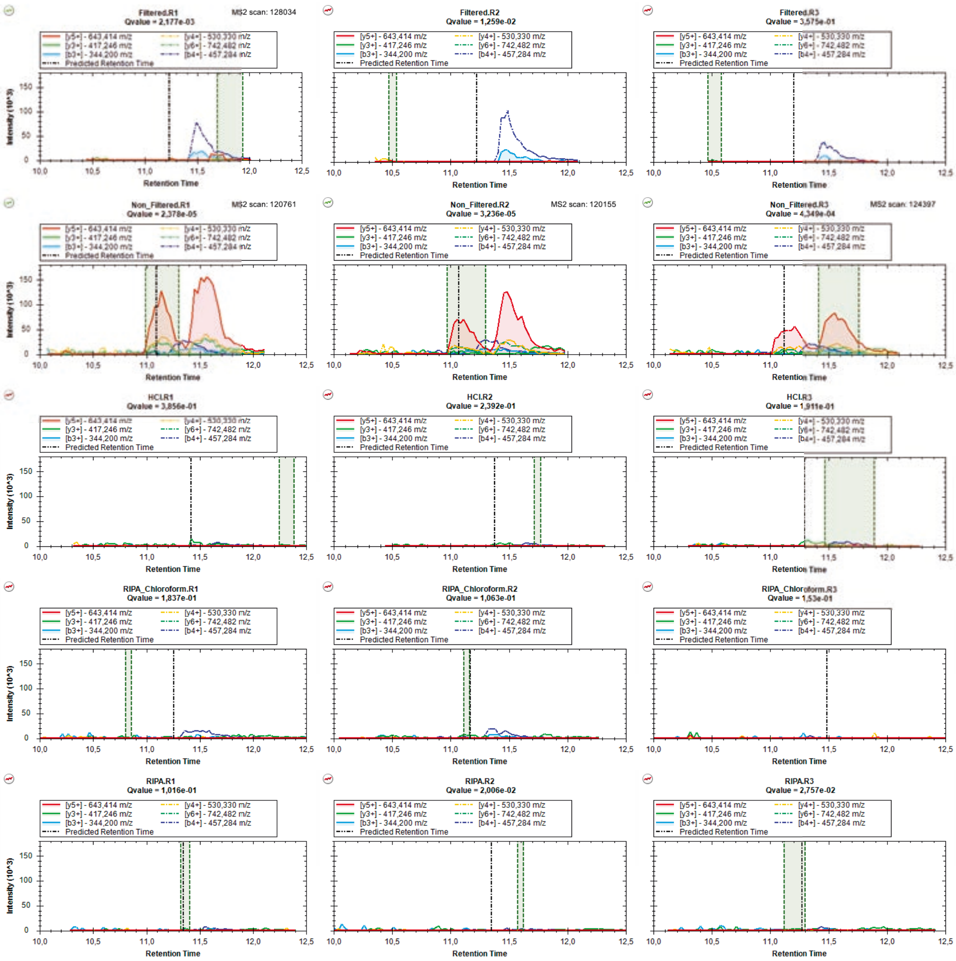

### mzspec_PXD066336_IP_568737_MVPTSQGFTEVR

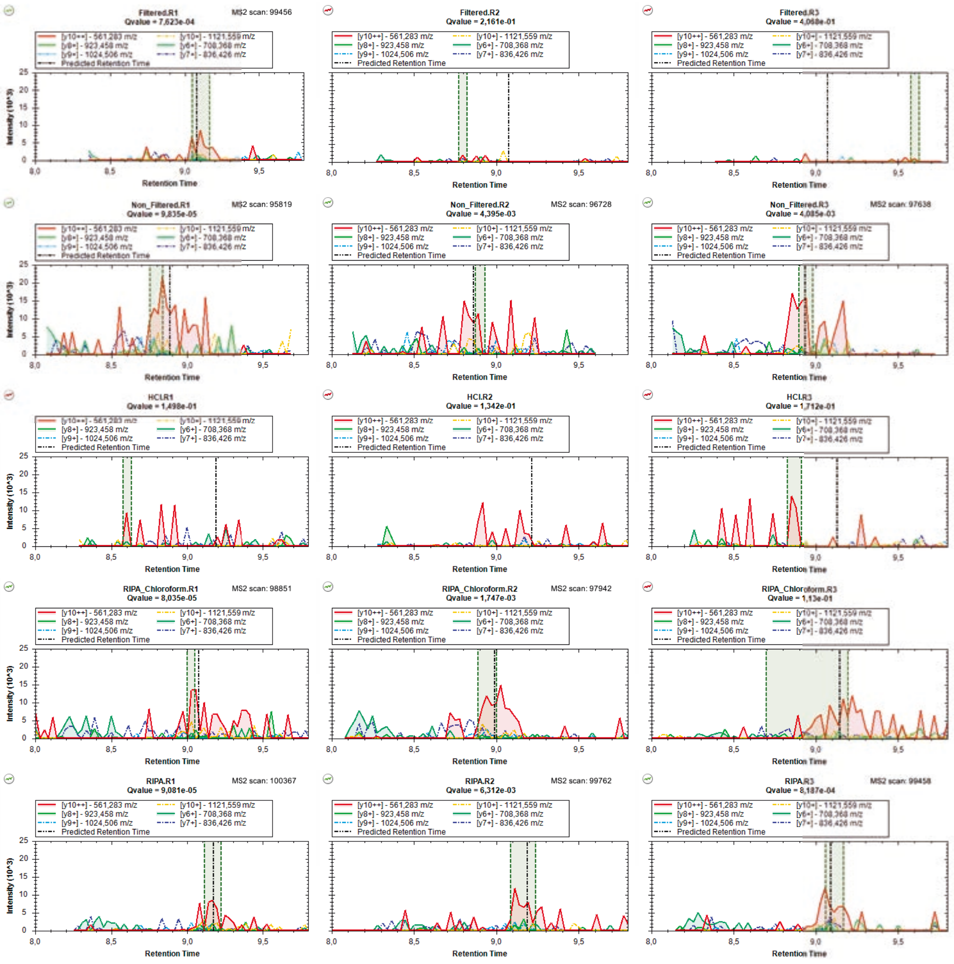

### mzspec_PXD066336_IP_674882_NISPGEISCLR

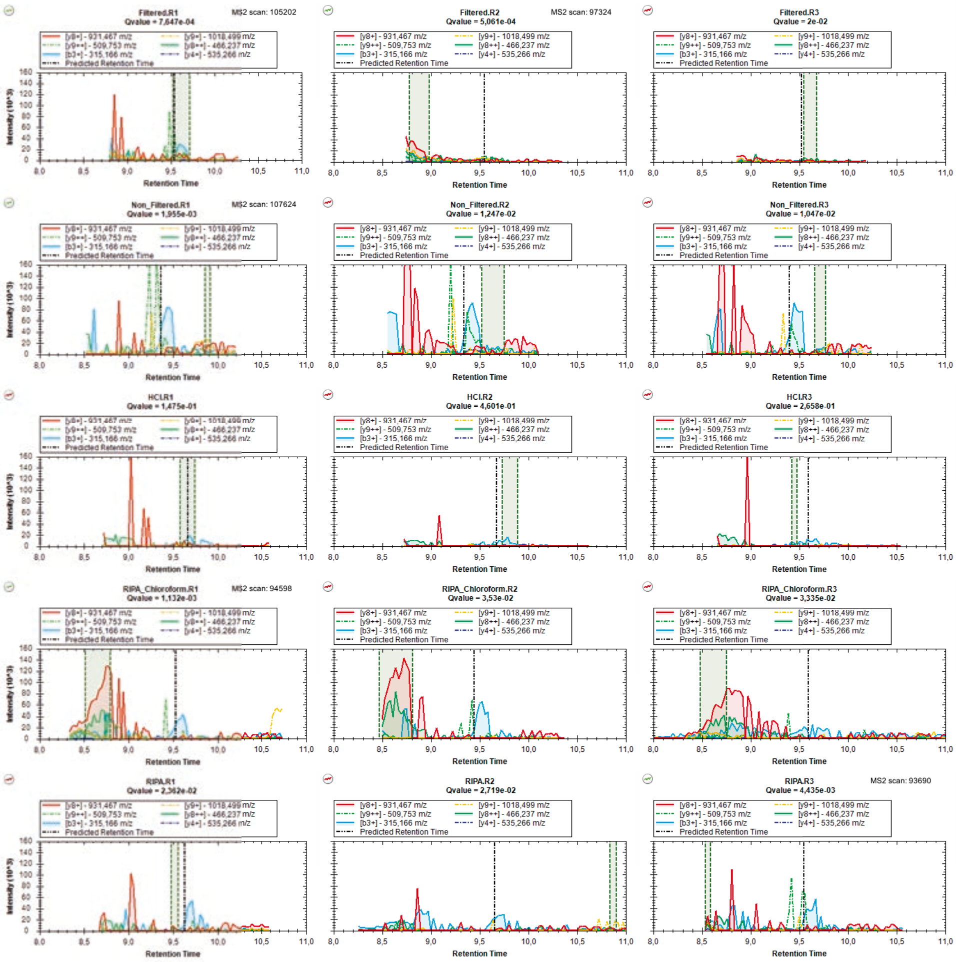

### mzspec_PXD066336_II_787098_NLTGTIPVPYR

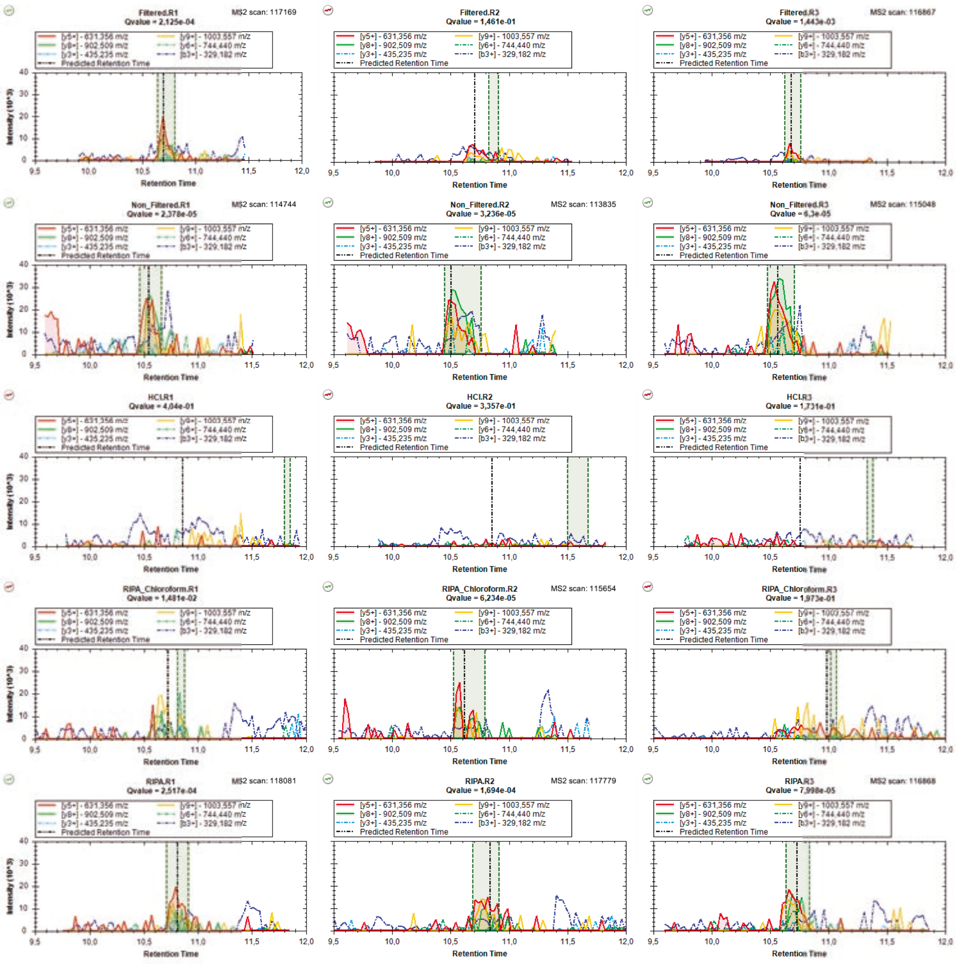

### mzspec_PXD066336_IP_3425048_PLLLLER

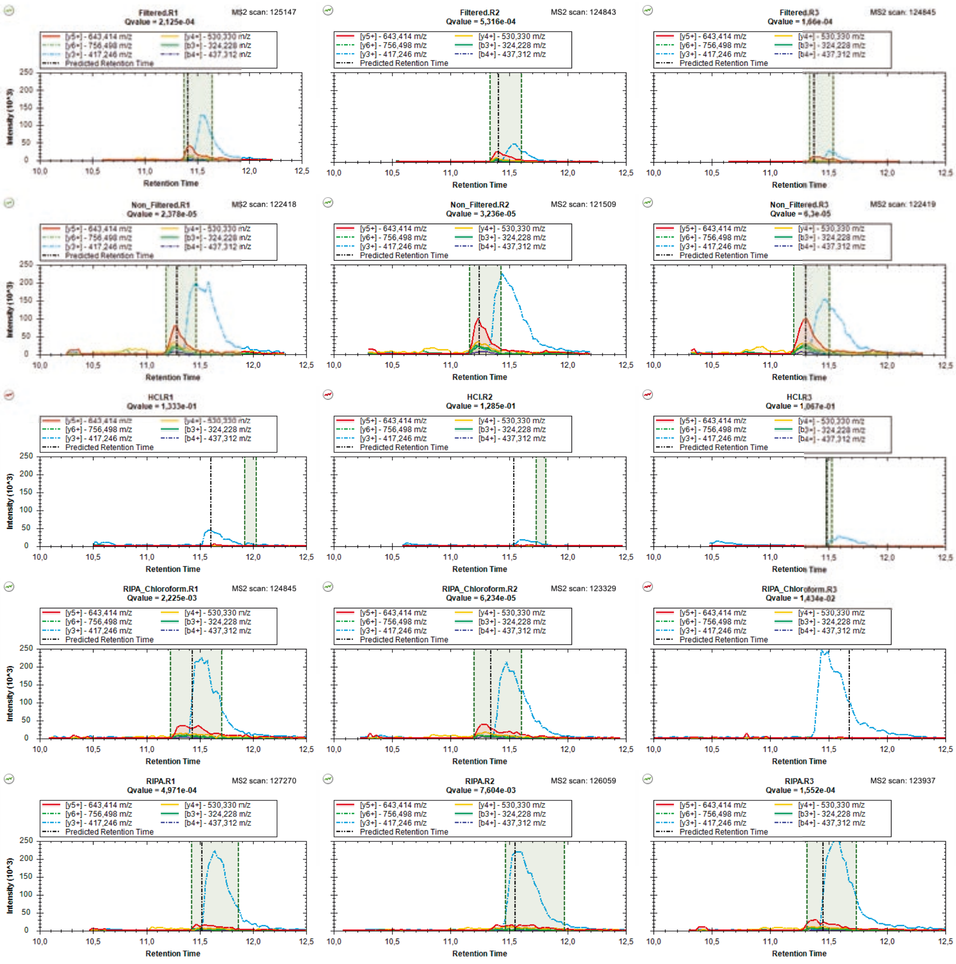

### mzspec_PXD066336_II_698681_PWNVDTLSK

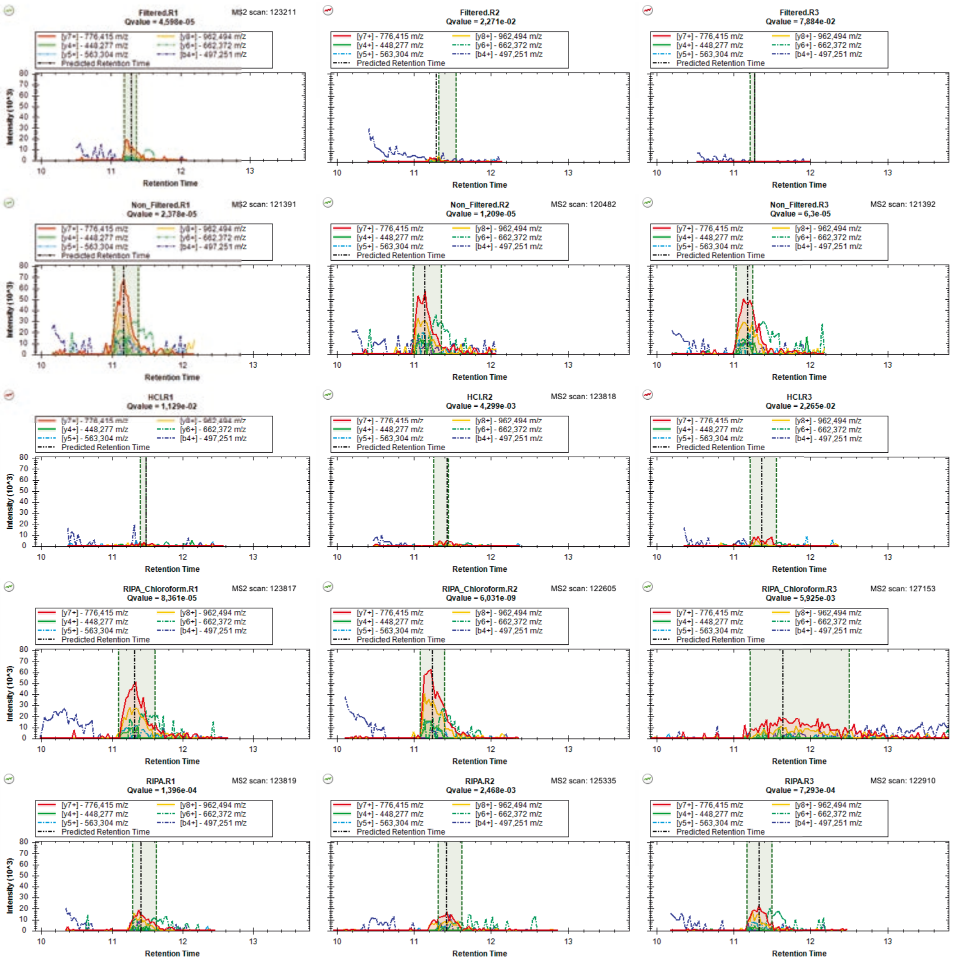

### mzspec_PXD066336_IP_762424_QAEVADQETK

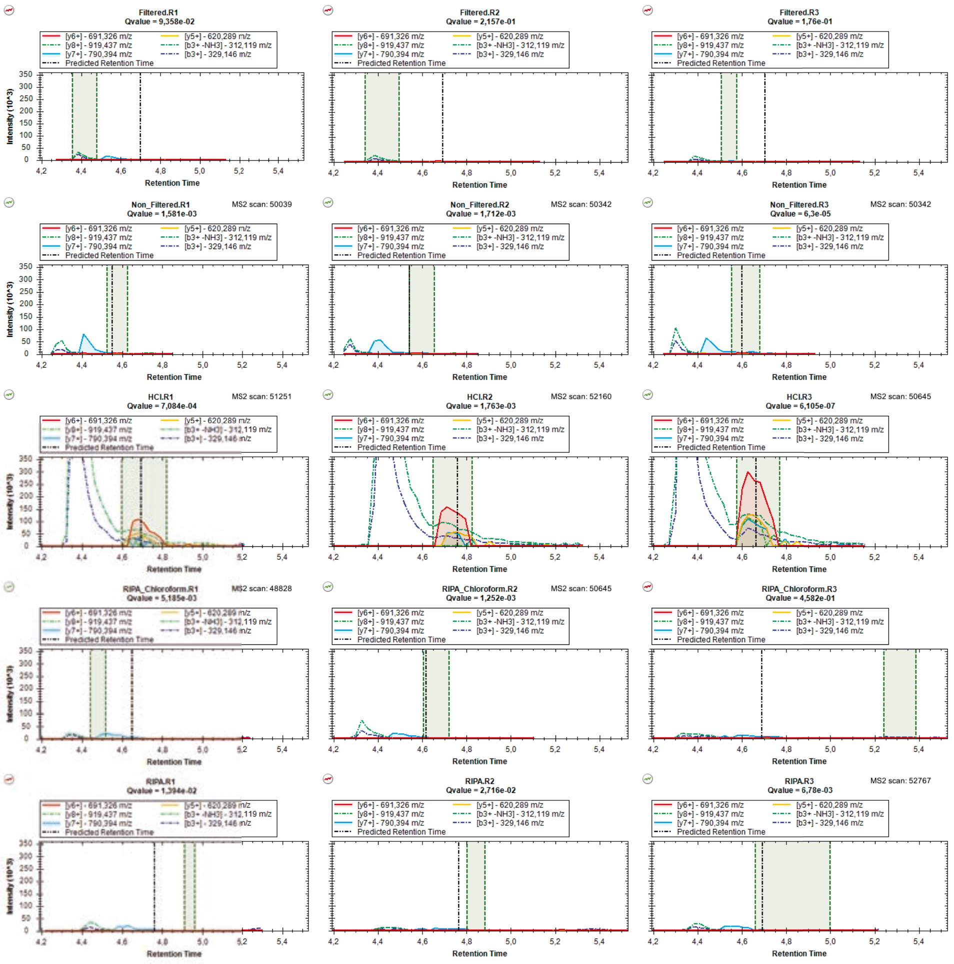

### mzspec_PXD066336_IP_660420_SCLICPQNPAFLSK

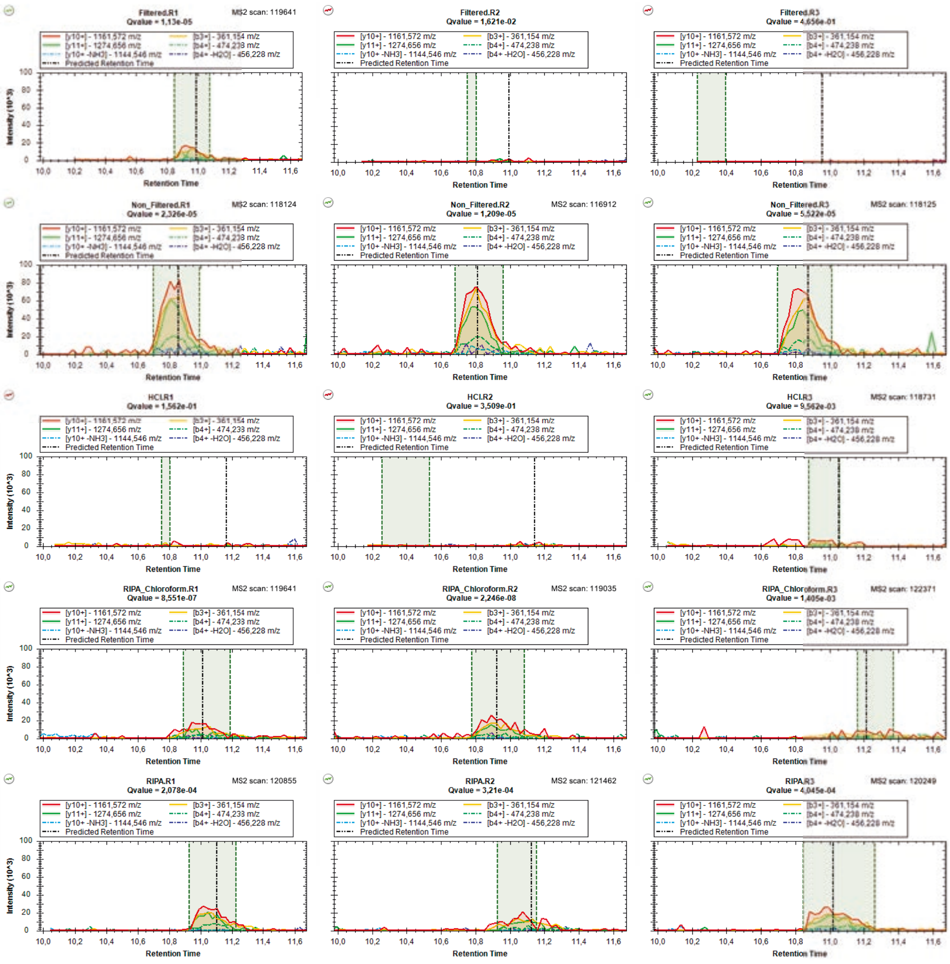

### mzspec_PXD066336_A0A1B0GW03_SPIQSGHTSCPGLPK_Charge_2

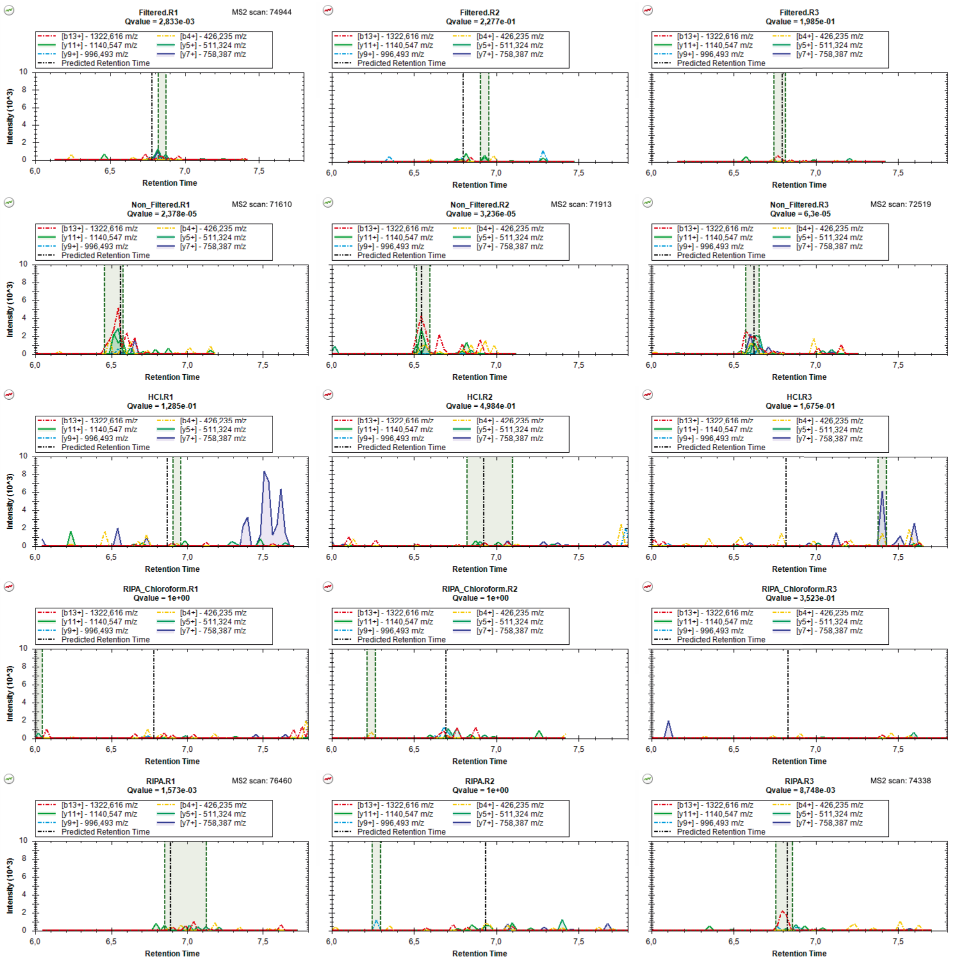

### mzspec_PXD066336_A0A1B0GW03_SPIQSGHTSCPGLPK_Charge_3

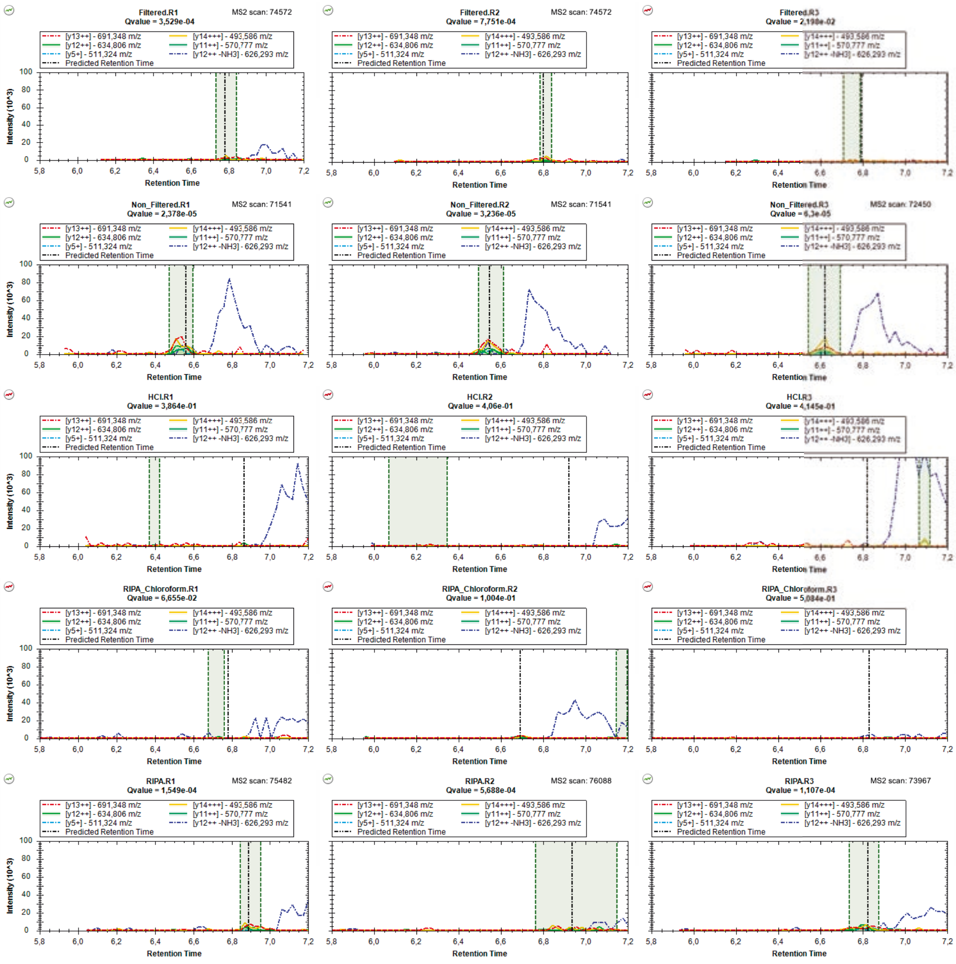

### mzspec_PXD066336_IP_3786303_SSLPSLGEK

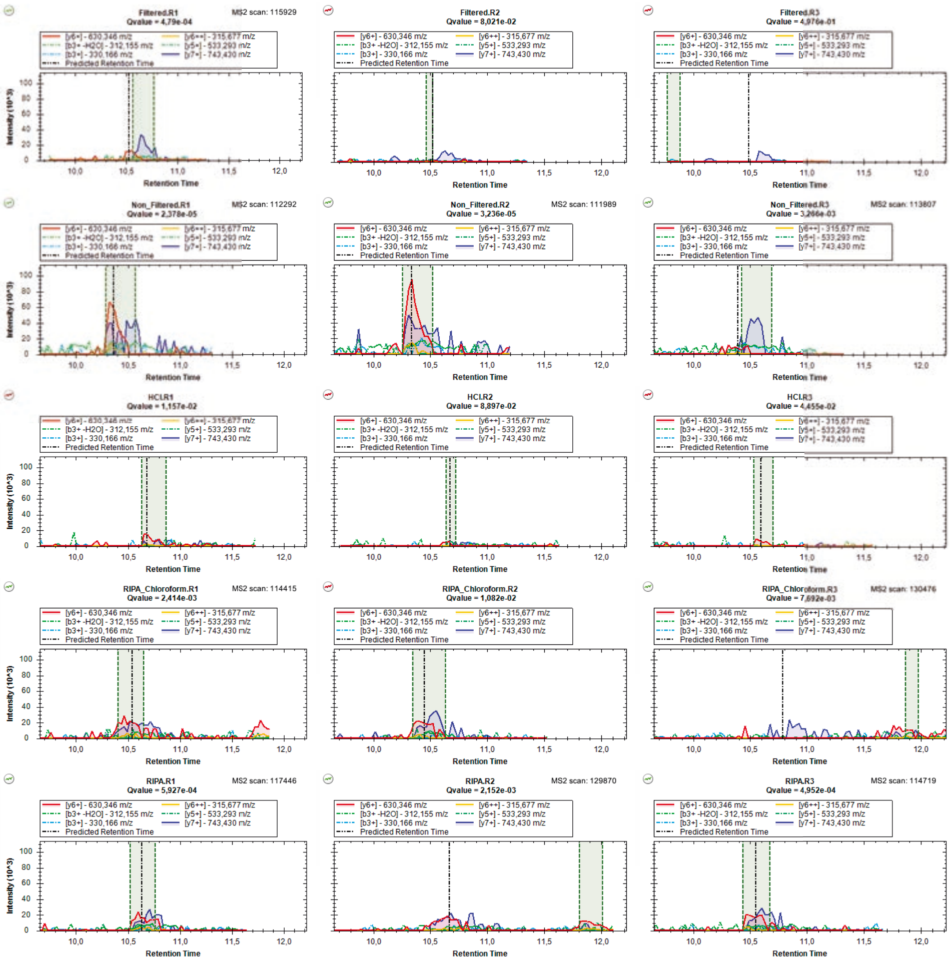

### mzspec_PXD066336_IP_573154_SSTPTPISR

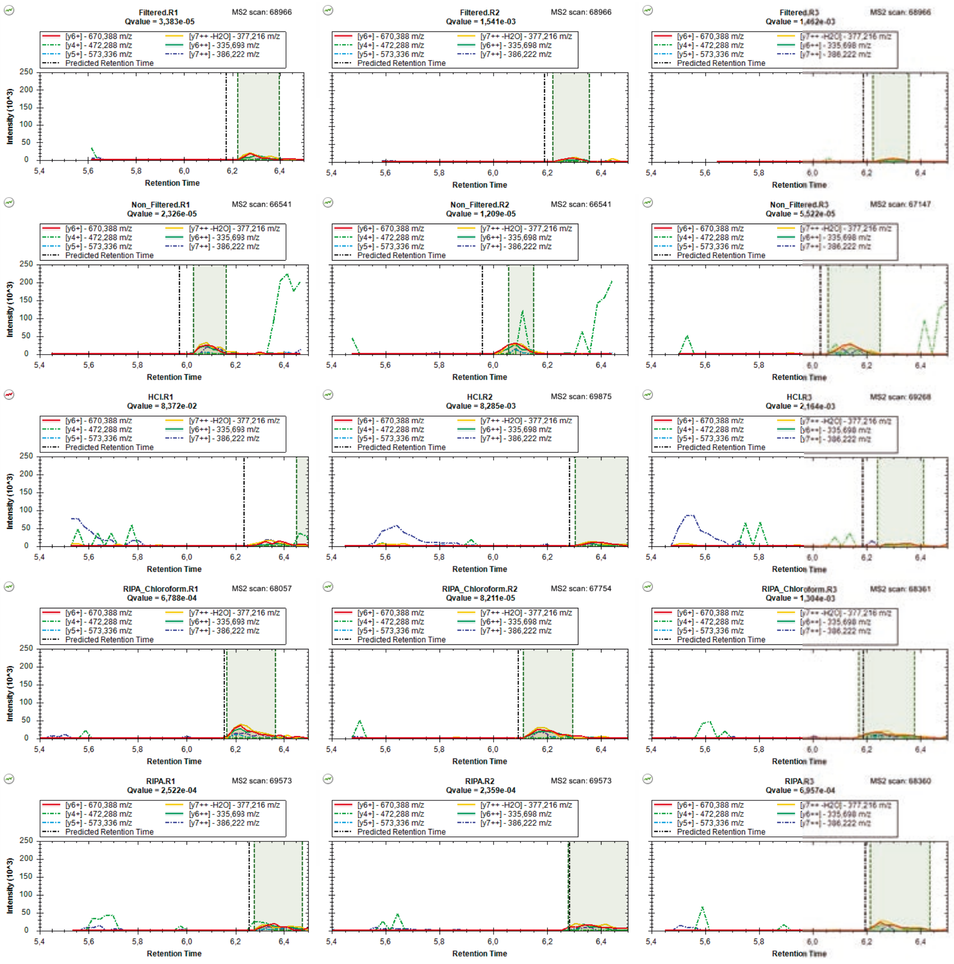

### mzspec_PXD066336_A0A1B0GW03_STSWVHPGTK

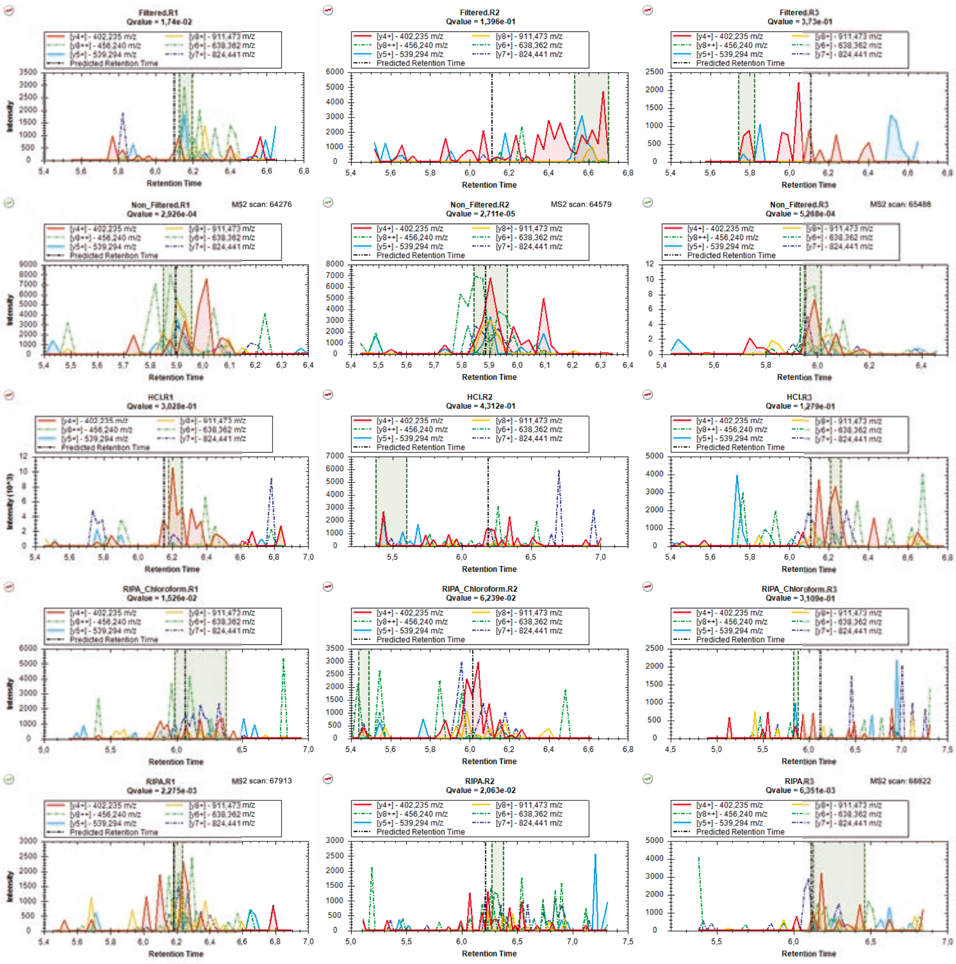

### mzspec_PXD066336_IP_773591_TQGFFGALNEQR

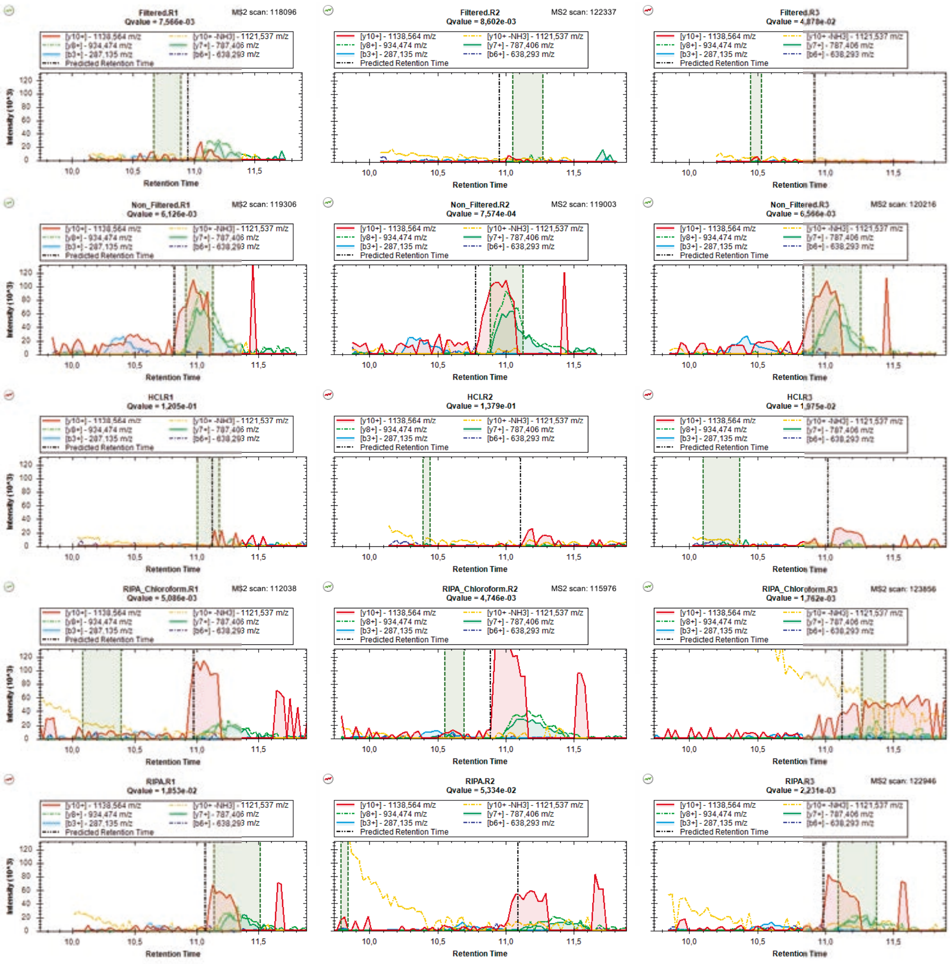
